# Polymorphic short tandem repeats make widespread contributions to blood and serum traits

**DOI:** 10.1101/2022.08.01.502370

**Authors:** Jonathan Margoliash, Shai Fuchs, Yang Li, Xuan Zhang, Arya Massarat, Alon Goren, Melissa Gymrek

## Abstract

Short tandem repeats (STRs), genomic regions each consisting of a sequence of 1-6 base pairs repeated in succession, represent one of the largest sources of human genetic variation. However, many STR effects are not captured well by standard genome-wide association studies (GWAS) or downstream analyses that are mostly based on single nucleotide polymorphisms (SNPs). To study the involvement of STRs in complex traits, we imputed genotypes for 445,720 autosomal STRs into genotype array data from 408,153 White British UK Biobank participants and tested for association with 44 blood and serum biomarker phenotypes. We used two fine-mapping methods, SuSiE and FINEMAP, to identify 119 high-confidence STR-trait associations across 93 unique STRs predicted as causal variants under all fine-mapping settings tested. Using these results, we estimate that STRs account for 5.2-7.6% of causal variants identifiable from GWAS signals for these traits. Our high confidence STR-trait associations implicate STRs in some of the strongest hits for multiple phenotypes, including a CTG repeat in *APOB* associated with circulating apolipoprotein B levels, a CGG repeat in the promoter of *CBL* associated with multiple platelet traits and a poly-A repeat in *TAOK1* associated with mean platelet volume. Replication analyses in additional population groups and orthogonal expression data further support the role of a subset of the candidate STRs we identify. Together, our study suggests that polymorphic tandem repeats make widespread contributions to complex traits, provides a set of stringently selected candidate causal STRs, and demonstrates the need to routinely consider a more complete view of human genetic variation in GWAS.

## Introduction

Genome-wide association studies (GWAS) have become an indispensable tool for identifying which genes and non-coding regions in the genome influence complex human traits. While GWAS routinely identify tens to hundreds of genomic regions associated with individual traits, biological interpretation of GWAS results remains challenging^1^. A major limitation is that typical GWAS pipelines only consider a subset of common genetic variants. The majority of GWAS have been based on common single nucleotide polymorphisms (SNPs) and short insertions or deletions (indels) either genotyped using microarrays or imputed from population reference databases based on whole genome sequencing (WGS) data. However, detailed follow-up of individual GWAS signals has often revealed complex variants that were absent from the original analysis, such as repeats^2,3^ or structural variants^4,5^, to be the causal drivers of those signals. Indeed, a recent study showed that polymorphic protein-coding variable number tandem repeats (VNTRs) are likely causal drivers of some of the strongest GWAS signals identified to date for multiple traits^2^.

Short tandem repeats (STRs) are a type of complex variant that consist of repeat units between 1-6bp duplicated many times in succession. Over one million STRs occur in the human genome^6^, each spanning from tens to thousands of base pairs. STRs undergo frequent mutations resulting in gain or loss of repeat units^7^, with per-locus mutation rates several orders of magnitude higher than average rates for SNPs^8^ or indels^9^. Large repeat expansions at STRs are known to result in Mendelian diseases such as Huntington’s disease, muscular dystrophies, hereditary ataxias and intellectual disorders^10,11^. Further, recent evidence suggests that more modest but highly prevalent variation at multi-allelic non-coding STRs can also be functionally relevant. We and others have found associations between STR length and both gene expression^3,12,13^ and splicing^14,15^. The impact of non-coding STRs on gene expression is hypothesized to be mediated by a variety of mechanisms including modulating nucleosome positioning^16^, altering methylation^12,17^, affecting transcription factor recruitment^3^ and impacting the formation of non-canonical DNA^18,19^ and RNA^20,21^ secondary structures. Together, these suggest that STRs potentially play an important role in shaping complex traits in humans.

Despite this potential, STRs are not well-captured by current GWAS. Because STRs are not directly genotyped by microarrays and are challenging to analyze from WGS, STRs have been largely excluded from widely used reference haplotype panels^22–24^ and downstream GWAS analyses. While some STRs are in high linkage disequilibrium (LD) with nearby SNPs, many highly multi-allelic STRs can only be imperfectly tagged by individual common SNPs, which are typically bi-allelic. Thus effects driven by variation in repeat length have likely not been fully captured, especially at those highly polymorphic STRs which cannot be well approximated by a bi-allelic indel or well-tagged by bi-allelic SNPs.

Recent technological advances can now enable incorporation of STRs into GWAS. We and others have created a variety of bioinformatic tools to genotype STRs directly from WGS by statistically accounting for the noise inherent in STR sequencing^6,25–29^. We previously used these tools to develop a reference haplotype panel consisting of both SNP and STR genotypes that allows for imputation of STRs from genotype array data^30^ in samples for which WGS is unavailable. In that study we found that all but the most highly polymorphic STRs are amenable to imputation in European cohorts, with an average per-locus imputation concordance of 97% with genotypes obtained from WGS.

Here, we leverage our SNP-STR reference haplotype panel to impute genome-wide STRs into SNP array data from 408,153 White British individuals obtained from the UK Biobank (UKB) for which deep phenotype information is available^31^. Whereas a recent publication studied the effects of protein-coding VNTRs (118 total VNTRs with repeat units of 7+ base pairs; total length of up to several kilobases) on complex traits^2^, our study focuses on a distinct set of repeats (namely 445,720 STRs with repeat units of 1bp to 6bp) which are mostly non-coding. We test for association between imputed STR lengths and 19 blood cell count and 25 biomarker traits. These traits provide multiple advantages: they are broadly and reliably measured, continuous, highly polygenic and have variants with relatively large effect sizes, thus enabling well-powered association testing.

We performed fine-mapping on these associations and estimate that STRs account for 5.2-7.6% of signals identified by GWAS for the traits we studied. We observed that fine-mapping results in some instances are substantially influenced by the choice of fine-mapper, and additionally are moderately sensitive to data-processing choices and fine-mapper instabilities, and thus require careful interpretation. After restricting to the signals which were consistently fine-mapped across multiple fine-mappers and fine-mapping settings, we identified 93 unique STRs strongly predicted to be causal for at least one trait. We highlight multiple STRs in this set which we predict contribute to some of the strongest hits for many traits, including apolipoprotein B, platelet crit and mean platelet volume. Overall, our study demonstrates the widespread role of polymorphic tandem repeats and the need to consider a broad range of variant types in GWAS and downstream analyses such as fine-mapping.

## Results

### Performing genome-wide STR association studies in 44 traits

We imputed genotypes for 445,720 autosomal STRs into phased genotype array data from 408,153 White British individuals from the UKB using Beagle^32^ in combination with our published SNP-STR reference haplotype panel^30^ (**Methods**; **Fig. 1a**; **Supplementary Fig. 1; URLs**). We find that this imputation yields broadly similar genotypes to WGS (see below). Compared to common SNPs, which are typically bi-allelic, many of the imputed STRs are highly multi-allelic (**Fig. 1b**). We tested STRs for association with 44 quantitative blood cell count and other biomarker traits (**Supplementary Table 1**) for which phenotype information was available for between 304,658-335,585 genetically unrelated subsets of individuals. To facilitate this and other STR association studies, we developed associaTR (**URLs**), an open-source software package for identifying associations between STR lengths (as measured by the number of repeat units) and phenotypes.

**Figure 1:**
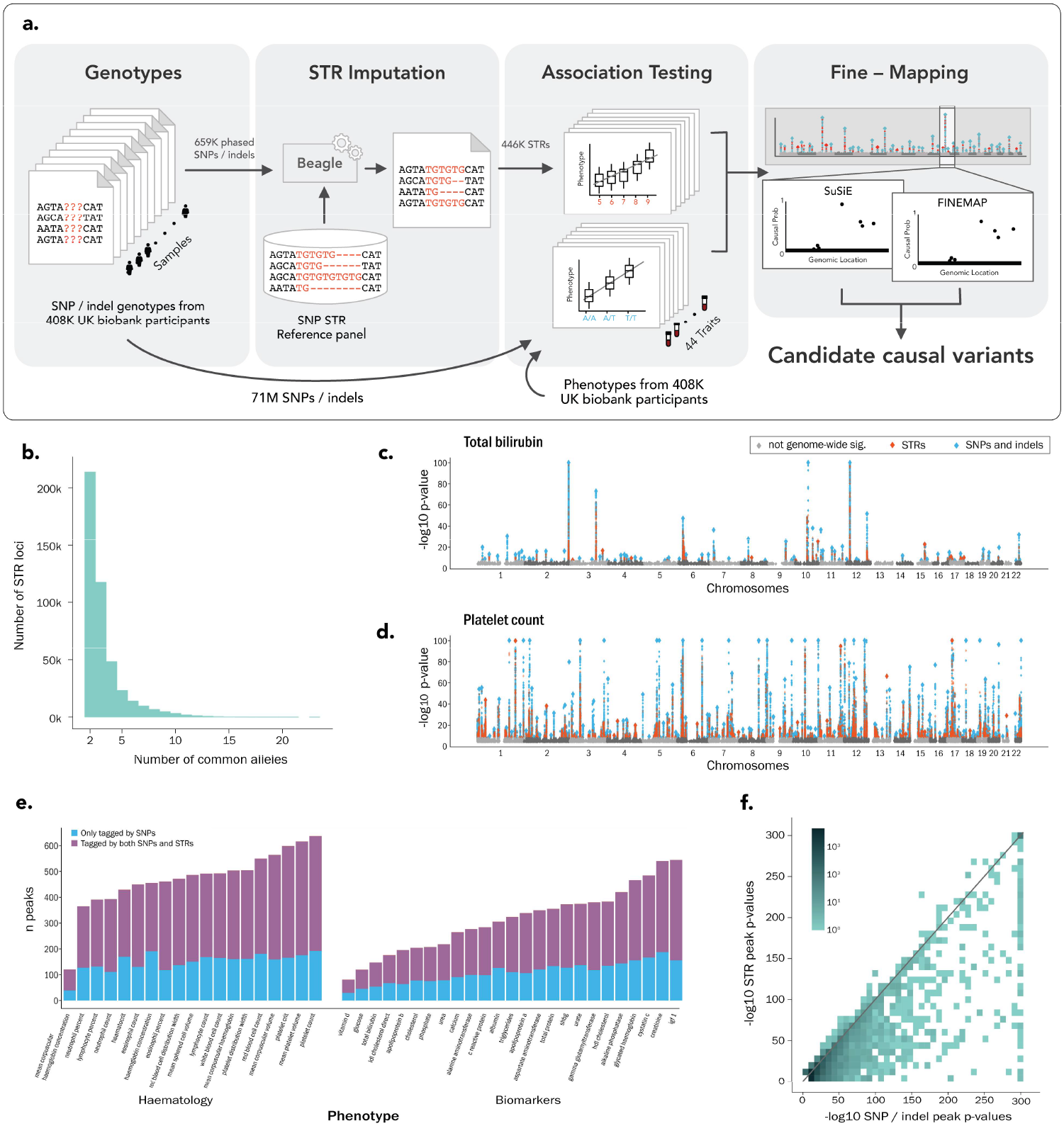
Genome-wide association tests identify STRs, SNPs and indels associated with blood and biomarker traits in the UKB. **(a) Schematic overview of this study**. STRs are imputed into phased hard-called variants obtained from genotype arrays. GWAS is performed on SNPs and STRs in parallel. Regions with significant signals are identified and then fine-mapped by two independent methods each under multiple scenarios, resulting in candidate causal STRs. **(b) Distribution of the number of common alleles at each imputed STR**. Common alleles are defined as alleles with estimated frequency ≥1% (**Methods**). For clarity we omitted from this figure the 237 imputed STRs with only a single imputed allele with frequency ≥1%. **(c-d) Representative association results**. Manhattan plots are shown for phenotypes (c) total bilirubin (an example moderately polygenic trait) and **(d)** platelet count (an example highly polygenic trait). Large diamonds represent the lead variants (pruned to include at most one lead variant per 10Mb for visualization). -log10 p-values are truncated at 100. Blue=SNPs and indels; orange=STRs. **(e) Summary of signals identified for each trait**. Bars show the number of peaks per phenotype. Blue denotes peaks only containing genome-wide significant SNPs and indels, purple denotes peaks containing both significant SNPs/indels and STRs. The number of peaks only containing significant STRs is too small to be visible in this display. **(f) Comparison between lead SNP and indel and STR p-values at each peak**. If there are no STRs in a peak, the y coordinate is set to zero (same for SNPs and indels). p-values are capped at 1e-300, the maximum precision of our pipeline. The shade represents the number of peaks falling at each position on the graph. The bottom left tile (which only contains peaks whose lead SNP/indel and STR variants fall in the least significant bin) has been removed so as not to not skew the scale of the color bar.

For each STR-trait pair, we used associaTR to test for a linear association between STR dosage (the sum of the imputed allele length dosages of both chromosomes) and the measured trait value (**Fig. 1c-d**). For comparison, we used plink^33^ to perform similar association tests using 70,698,786 SNP and short indel variants that were imputed into the same individuals. For all associations (STR, SNP and indel), we included as covariates SNP-genotype principal components, sex, and age (**Methods**). Additional covariates were included on a per-trait basis (**Supplementary Table 1**). We compared the output of our SNP analysis pipeline to previous results reported by Pan UKBB^34^ and found that our pipeline produced similar results, although had slightly weaker p-values overall. (**Supplementary Fig. 2**).

We then compared signals identified by SNPs and indels to those identified by STRs. For each trait we defined peaks as non-overlapping 250kb intervals centered on the lead genome-wide significant variant (a SNP, indel or STR with p<5e-8) in that interval (**Methods**). We identified an average of 389 peaks per trait, with blood cell count traits generally more polygenic than other biomarkers (**Fig. 1e**). Of these peaks, 65.9% contained both a significant STR and a significant SNP or indel, 32.5% contained only significant SNPs or indels, and 1.7% contained only significant STRs. The majority of strong peaks (containing any variant with p<1e-100) were identified by both STRs and SNPs and/or indels, in that they contain both an STR and a SNP or indel with p<1e-80. No new strong peaks were identified only by STRs (**Fig. 1f**), which is unsurprising given that SNP and indel genotypes were used to impute the STRs. Overall, p-values of the lead SNP or indel and lead STR were similar for most peaks. Thus, we focused on fine-mapping to determine which variants might be causally driving the identified signals.

### Fine-mapping suggests 5.2-7.6% of significant signals are driven by STRs

We applied statistical fine-mapping to identify candidate causal variants that may be driving the GWAS signals detected above. We used two fine-mapping methods: SuSiE^35^ and FINEMAP^36^. These methods differ in their modeling assumptions and thus provide partially orthogonal predictions. For each trait we divided its genome-wide significant variants (SNPs, indels and STRs) and nearby variants into non-overlapping regions of at least 250kb (**Methods**). This resulted in 14,491 fine-mapping trait-regions (**Supplementary Table 2**), with some trait-regions containing multiple nearby peaks. To compare outputs between fine-mappers in downstream analyses, we defined the causal probability (CP) of each variant to be a number between 0 and 1 that indicates the variant’s chance of causality. For FINEMAP we defined a variant’s CP to be the FINEMAP posterior inclusion probability (PIP) calculated for that variant. For SuSiE we defined a variant’s CP to be the maximal SuSiE alpha value for that variant across pure credible sets in the region (**Supplementary Figs. 3-4**). We explain the rationale behind this choice in **Supplementary Note 1**.

We used two approaches to study the contribution of STRs vs. SNPs and indels to fine-mapped signals. First, we focused on the genome-wide significant variants (STR, SNP, or indel) with CP ≥ 0.8. (These accounted for a minority of the total 21,045 pure signals detected by SuSiE and the total 33,756 signals detected by FINEMAP). SuSiE identified 4,494 such variants and FINEMAP identified 5,170. Of these, 7.4% (range 1.3-13.0% across traits; SuSiE) and 7.6% (range 1.4- 14.0%; FINEMAP) are STRs. Among the subset of variants identified by both methods (3,961), 5.4% (range 1.0-11.1%) are STRs. Second, we considered the sum of CPs from all genome-wide significant variants, thereby taking into account the many signals which were not resolved to a single variant. STRs make up 5.2% (range 1.1-6.8%) of the total SuSiE CP sum and 7.4% (range 2.9-9.0%) of the total FINEMAP CP sum. A potential limitation of this second metric is that variants with small CPs (CP ≤ 0.1) represent a large fraction (29.3% for SuSiE, 35.1% for FINEMAP) of these totals (**Supplementary Fig. 5**). Additionally, our results below suggest that a sizable subset of variant CPs are either discordant between fine-mappers or unstable, particularly for STRs (**Supplementary Notes 2-3**), impacting the totals in both metrics. Nevertheless, the results above suggest that between 5.2-7.6% of causal variants identifiable from GWAS can be attributed to an STR, regardless of the fine-mapping method or metric used. This is in line with the total percentage of non-major alleles per person and roughly half the total percentage of base pair variation per person accounted for by STR lengths compared to other variant types in our study (**Supplementary Table 3**). We report all 511 genome-wide significant STR associations across 409 distinct STRs with either FINEMAP or SuSiE CP ≥ 0.8 in **Supplementary Table 4**, and a subset of those which pass more stringent thresholding in **Supplementary Table 5** (see below).

To evaluate the reliability of our approach for determining the relative contributions of STRs vs SNPs and indels, we performed fine-mapping simulation analyses assuming a simple additive model. We used two strategies for simulating phenotypes, in each case simulating only causal SNPs and indels, and assessed to what extent STRs were incorrectly identified by SuSiE or FINEMAP as contributing to the underlying signals. For the first strategy, we chose between one and three causal SNPs or indels at random, weighting by minor allele frequencies, for a total of 1644 simulations. For the second, we chose causal variants to be those indicated by SuSiE as being potentially causal for a representative trait (platelet count), in an attempt to more closely capture properties of truly causal variants during simulation, for a total of 1374 simulations. These simulations procedures and rationales are described in more detail in the **Methods**, **Supplementary Table 6** and **Supplementary Fig. 6**.

When simulating from one to three randomly chosen causal SNPs or indels, of genome-wide significant variants that SuSiE or FINEMAP assigned CP ≥ 0.8, between 0% and 0.46% were STRs. In contrast, using phenotypes simulated from the second strategy, between 1.4% and 3.2% were STRs (**Supplementary Table 7**). Across all genome-wide significant variants, of the total CP assigned by SuSiE or FINEMAP, 0.50% to 0.95% and 3.1% to 3.2% were assigned to STRs in the first and second simulation strategies, respectively (**Supplementary Table 8**). These numbers are uniformly lower than the 5.2-7.6% STR contributions identified above. This suggests that if the 44 traits studied here have genetic architectures similar to the phenotypes we simulated, the results above are unlikely to be fully explained by systematic bias of these fine-mappers in favor of STRs. However, we expect there are complexities to the genetic architecture of blood traits that are not captured by our simulations, and we cannot rule out the possibility that they could result in such bias. These results also suggest that some of the fine-mapped STRs may be false positives, and thus that our estimates of STR contributions could be somewhat inflated. On the other hand, we also observed that a large fraction (66-81%) of simulated causal SNPs and indels are not assigned CP ≥ 0.8 by fine-mapping, and we see a similar lack of sensitivity in limited simulations performed with causal STRs (**Supplementary Table 7**). We expect this low sensitivity is a greater source of uncertainty regarding the relative contribution of each variant type than the false positive rates reported here.

We evaluated imputation quality at the 409 STRs in **Supplementary Table 4** by comparing imputed genotypes to genotypes obtained from recently released WGS data for 200,025 UKB individuals. At each locus, we computed the Pearson r^2^ between length dosages from imputation and summed lengths from WGS hard-calls, in addition to other per-locus metrics (**Supplementary Table 4**). For 78.7% of these STRs, per-locus r^2^ values are greater than 0.9 and for 92.7% they are greater than 0.8. Other measurements of imputation concordance perform comparably well (**Supplementary Fig. 7; Methods**). Overall, they suggest that fine-mapping results from imputed data are unlikely to systematically differ from fine-mapping of hard-called genotypes for these loci. Unless otherwise stated, all results reported below are based on imputed genotypes.

### Identifying and characterizing confidently fine-mapped STRs

We performed additional analyses to identify a high confidence set of causal STR candidates. First, we noticed that while SuSiE and FINEMAP tended to output similar results, they assigned highly discordant CPs to a subset of variants (**Supplementary Note 2**; **Supplementary Figs. 8- 10**). Thus we conservatively restricted our follow-up analyses to the 167 candidate STR associations with association p-values < 1e-10 and with CP ≥ 0.8 in both FINEMAP and SuSiE. Second, to confirm that the default fine-mapping settings did not appreciably dictate our results, we reran SuSiE and FINEMAP under a range of alternative settings (**Methods**), including using best-guess STR genotypes instead of dosages and varying the prior distribution of effect sizes. These additional fine-mapping runs tended to produce concordant results, but again for a subset of STRs, produced highly inconsistent CPs (**Supplementary Figs. 11-14**), which we mostly attribute to imputation uncertainty and FINEMAP instability (**Supplementary Note 3**). Thus, we further restricted to the 118 (70.7%) of the 167 STR-trait associations which also maintained CP ≥ 0.8 across these additional fine-mapping runs. We refer to the STR-trait associations meeting these criteria as confidently fine-mapped STR associations. Lastly, we added an association with an STR in the *APOB* gene to this set as this association only failed to meet the above criteria because the STR was simultaneously represented in both our STR reference panel and in the SNP and indel set generated by the UKB team (**Supplementary Note 4**). In sum, we were left with 119 confidently fine-mapped STR-trait associations corresponding to 93 distinct STRs, which we display in **Fig. 2** and **Supplementary Table 5**.

**Figure 2:**
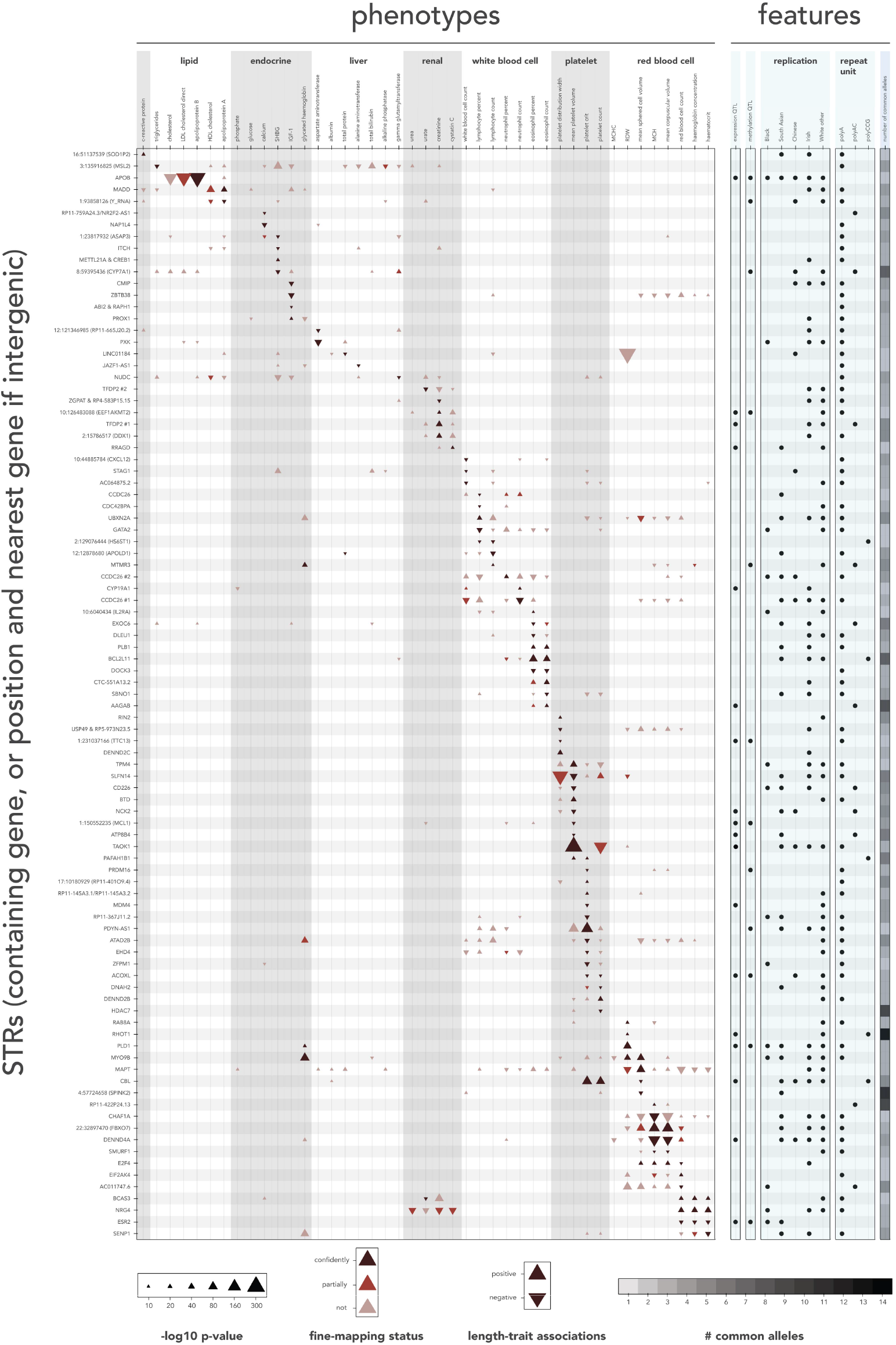
STRs are confidently fine-mapped to causally impact many traits. Only STRs with at least one confidently fine-mapped association are shown. Each triangle represents an STR-trait association with association p-value < 1e-10. Black=confidently fine-mapped, red-brown=CP ≥ 0.8 in either initial FINEMAP or SuSiE run, light-tan=all other associations with p-values < 1e-10. Triangle direction (up or down) indicates the sign of the association between STR length and the trait. Triangle size scales with association p-value. Similar traits are grouped on the x-axis by white and light-grey bands. STRs are grouped on the y-axis according to the traits they were confidently fine-mapped to. STRs are labeled by the genes they reside in (protein coding genes preferred) or by chromosomal location and the nearest gene for intergenic STRs. *CCDC26* and *TFDP2* each contain two confidently fine-mapped STRs and appear twice. Light blue rows indicate (from left to right): whether each STR is associated with expression of a nearby gene in the GTEx cohort (adjusted p<0.05; **Supplementary Table 12**), associated with the methylation of a nearby CpG island (**Supplementary Table 14**), replicates with the same direction of effect in other populations (adjusted p<0.05; **Methods**), repeat unit, and the number of common alleles for each STR (as defined in **Fig. 1**; see scale beneath). Additionally, we mark the STRs in *TAOK1* and *RHOT1* as being expression QTLs even though they did not pass WGS call rate filters in the GTEx dataset, as when we imputed the *TAOK1* STR into the GTEx dataset it was associated with *TAOK1* expression (**Methods**) and the *RHOT1* STR was associated with *RHOT1* expression in the Geuvadis dataset (**Methods**). The data summarized here is available in **Supplementary Tables 4, 5, 12** and **14**.

We evaluated these fine-mapping results by measuring their replication rates in populations besides White British individuals, with the expectation that causal associations will replicate at higher frequencies in other populations than non-causal associations due to having common biological functionality. The UKB includes self-identified groups of 8,043 Black, 7,952 South Asian, 1,568 Chinese, 12,957 Irish and 16,051 Other White participants that passed quality control, with about 40% of each population having WGS data (**Methods**). Using that WGS data we validated the imputed genotypes of those populations for the STRs in **Supplementary Table 4** and found that 78.2% and 93.9% of per-locus dosage r^2^ values are greater than 0.8 and 0.6 in the South Asian population, respectively, 45.5% and 84.8% in the Black population, and 64.3% and 84.8% in the Chinese population (**Supplementary Fig. 7**). These metrics are weaker in the non-White populations; this is expected given the largely European origin of our reference haplotype panel. Nevertheless, our results suggest imputed genotypes are sufficiently accurate across these groups for downstream analysis.

For each of these populations we performed follow-up STR association testing. Specifically, for each trait, for each fine-mapping region for that trait identified among White British individuals, we tested each STR in that region for association with that trait in each of the other populations (**Supplementary Table 9**; data for individual loci in **Supplementary Tables 4** and **5**). As expected, signals replicate at a higher rate in groups most closely related to our discovery cohort (i.e. Irish and Other White). Encouragingly, fine-mapped associations replicate at higher rates than non-fine-mapped associations in the Black, South Asian, and Chinese populations, even after stratifying by the discovery p-value (**Fig. 3**, **Supplementary Fig. 15**). To quantitatively measure this trend, for each population we fit a logistic regression model using whether signals replicated in that population as the outcome, the fine-mapping status of those associations as the independent variable, and their -log_10_(p-value) in the discovery cohort as a covariate (**Supplementary Table 10)**. This analysis further supports the conclusion that fine-mapped associations replicate at higher rates. Additionally, the model consistently predicts that confidently fine-mapped STR associations replicate at higher rates than STR associations fine-mapped by either fine-mapper alone, although only a subset of those predictions reached nominal significance, likely due to the small number of fine-mapped STR associations.

**Figure 3:**
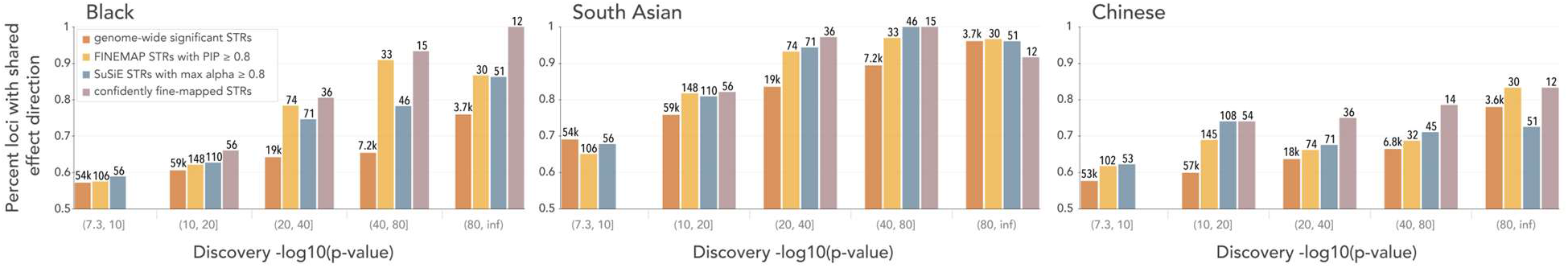
Concordance of White British STR effect directions in Black, South Asian and Chinese populations. The y-axis gives the fraction of STR associations measured in the discovery cohort that have the same direction of effect when measured in the replication population regardless of p-value. Brackets beneath the x-axis denote the binning of discovery -log_10_(p-values). Brown=genome-wide significant associations (discovery p<5e-8), orange=FINEMAP fine-mapped STR associations (discovery p<5e-8 and FINEMAP CP ≥ 0.8), teal=SuSiE fine-mapped STR associations (discovery p<5e-8 and SuSiE CP ≥ 0.8) and purple=confidently fine-mapped STR associations. Annotations above each bar indicate the number of STR-trait associations considered. We required confidently fine-mapped STR associations to have p-value < 1e-10, thus they do not appear in the left-most bin. The trends in these figures are somewhat sensitive to the choice of p-value bin boundaries so we additionally analyze this data using logistic models (**Supplementary Table 10**).

Next, we sought to characterize the set of confidently fine-mapped STRs. This set contains 62 poly-A repeats, 11 poly-AC, 5 poly-CCG, and 15 repeats with other units. Twelve of these STRs overlap coding or untranslated regions (UTRs) (**Table 1**; **Supplementary Table 11;** the two protein-coding repeats are further described in **Supplementary Note 4**; **Supplementary Fig. 16**). Compared to all genome-wide significant STRs, confidently fine-mapped STRs were more likely to be exonic trinucleotide STRs, in 5’ UTR regions or in genes that are not protein-coding (two-sided two-sample test of difference between proportions p=2e-26, 1e-3 and 2e-4 respectively). No other annotation categories that we tested showed significant enrichment or depletion after multiple hypothesis correction, likely due to low power arising from the small number of confidently fine-mapped STRs (**Methods**; **Supplementary Fig. 17**). Lastly, we observed that 18 of these confidently fine-mapped STRs are significant *cis* expression quantitative trait loci (QTLs) and 12 are significant *cis* DNA methylation QTLs in the Genotype-Tissue Expression (GTEx) dataset^37^ (**Fig. 2; Supplementary Tables 12-14**; **Supplementary Fig. 18; Methods**). We note that the expression and methylation analyses were underpowered because of low sample sizes, particularly for relevant tissue types (e.g. kidney, liver).

**Table 1:** Confidently fine-mapped STRs are identified in coding regions and untranslated regions (UTRs). Imputed alternate alleles and rsIDs are provided in **Supplementary Table 11**. Repeat units here are calculated as described in the **Methods**, except that they are required to be on the strand in the direction of transcription of the overlapping gene. We denote with asterisks the three STRs in this list which only appear in non-canonical transcripts for their genes from Ensembl release 106. Additionally, two STRs in this list only appear in transcripts or genes which are not protein coding. For all those STRs, we provide Ensembl transcript numbers followed by parentheses containing the Ensembl transcript support level, a number from 1 to 5, with 1 being the highest level of evidence and 5 being the lowest. Additional analysis and discussion of the potential impact of the protein-coding repeats in *APOB* and *E2F4* are given in **Supplementary Note 4**.

| STR coordinate (hg19 chr:pos) | Reference allele | Repeat unit | Trait | Association p-value | Association Z-score | Gene (annotation) | Transcription direction |
| --- | --- | --- | --- | --- | --- | --- | --- |
| 1:204527033 | (TAA)<sub>9</sub> | AAT | platelet crit | 5.76e-17 | -8.37 | *MDM4* (3' UTR) | + |
| 2:21266752 | (CAG)<sub>6</sub>(CGCAGGCAG)<br>[CGC(CAG)<sub>2</sub>]<sub>2</sub>CGC | CTG (poly-leucine) | Apolipo-protein B | 1.37e-279 | -35.76 | *APOB* (Coding) | - |
| 2:106510441 | (AC)<sub>6</sub>GTG(CA)<sub>10</sub>C<br>(TA)<sub>7</sub>T | AC | mean platelet volume | 6.93e-29 | -11.15 | *NCK2* (3' UTR) | + |
| 2:111878544 | (CGC)(CGCTGC)<sub>2</sub><br>(CGC)<sub>13</sub>C | CCG | eosinophil count | 4.96e-58 | +16.06 | *BCL2L11* (5' UTR) | + |
|  |  |  | eosinophil percent | 5.88e-75 | +18.32 |  |  |
| 2:204311891 | T<sub>4</sub>CT<sub>4</sub>CT<sub>3</sub>CT<sub>18</sub> | T | IGF-1 | 3.97e-11 | -6.61 | *ABI2* (3' UTR*)<br>ENST00000295851.10 (1) | + |
| 6:90121977 | (TC)<sub>7</sub> | TC | cystatin C | 1.24e-16 | +8.28 | *RRAGD* (3' UTR) | - |
| 11:119077000 | (CGG)<sub>11</sub>C | CGG | mean sphered cell volume | 6.88e-16 | -8.07 | *CBL* (5' UTR*)<br>ENST00000634586.1 (5) | + |
|  |  |  | platelet count | 3.77e-83 | +19.32 |  |  |
|  |  |  | platelet crit | 6.07e-103 | +21.55 |  |  |
| 15:40312923 | T<sub>16</sub> | T | red blood cell count | 2.27e-20 | +9.25 | *EIF2AK4* (Not protein coding*)<br>ENST00000558743.1 (2) | + |
| 16:67229794 | (CAG)<sub>13</sub>(CAA)(CAG)<br>(TAA)(CAG)<sub>3</sub> | AGC (poly-serine) | mean sphered cell volume | 3.07e-23 | +9.93 | *E2F4* (Coding) | + |
|  |  |  | red blood cell count | 1.08e-13 | -7.43 |  |  |
|  |  |  | mean corpuscular haemoglobin | 2.83e-23 | +9.94 |  |  |
|  |  |  | mean corpuscular volume | 9.27e-26 | +10.49 |  |  |
| 17:30469471 | (CCG)<sub>16</sub>CC | CCG | red blood cell distribution width | 6.57e-13 | +7.19 | *RHOT1* (5' UTR) | + |
| 17:33871548 | T<sub>17</sub> | A | mean platelet volume | 4.30e-62 | -16.63 | *SLFN14* (3' UTR) | - |
| 20:32971954 | A<sub>20</sub> | A | shbg | 6.37e-15 | -7.80 | Y RNA<br>ENST00000364628.1 (3) | - |

### Fine-mapped STRs capture known associations

We identified multiple fine-mapped STRs that were previously demonstrated to have functional roles, providing supporting evidence for the validity of our pipeline. For instance, our confidently fine-mapped set implicates a protein-coding CTG repeat (**Supplementary Table 11**) to be the causal variant for one of the strongest signals for apolipoprotein B (p=1e-279; in one of the four peaks for this trait with minimal p-value exceeding our numeric precision) which forms the backbone of LDL cholesterol lipoproteins^38^. This locus is also one of the strongest signals for LDL cholesterol levels (two-sided association t-test p-value = 6e-236; fifth most-significant genome-wide peak) and the STR was marked as causal in 8 of 9 fine-mapping conditions. This repeat is bi-allelic in the UKB cohort with an alternate allele corresponding to deletion of three residues (Leu-Ala-Leu) in the signal peptide coded in the first exon of the apolipoprotein B (*APOB*) gene^39^. This deletion occurs in an imperfect region of the CTG repeat, with sequence CTGGCGCTG. In agreement with a previous study^40^, we found that the short allele is associated with higher levels of both analytes. We discuss this locus further in **Supplementary Note 4**.

As another example, our initial fine-mapping implicates a multi-allelic AC repeat (**Supplementary Table 11**) 6bp downstream of exon 4 of *SLC2A2* (also known as *GLUT2*, a gene that is most highly expressed in liver) as causally impacting bilirubin levels (p=9e-18). However, this repeat was not included in the final confidently fine-mapped set due to its FINEMAP CP of 0.61 not passing our 0.8 threshold, despite its SuSiE CP of 0.99. The potential link between GLUT2 and bilirubin is described in **Supplementary Note 5.** Previous studies in HeLa and 293T cells showed that inclusion of exon 4 of *SLC2A2* is repressed by the binding of mRNA processing factor hnRNP L to this AC repeat^41,42^, implicating this STR in *SLC2A2* splicing. Notably, these studies did not investigate the impact of varying repeat copy number. We examined this STR in GTEx liver samples and did not find a significant linear association between repeat count and the splicing of exon 4, though we did find evidence for association with the splicing of exon 6 (**Supplementary Fig. 19**).

### A trinucleotide repeat in CBL regulates platelet traits

Most of the confidently fine-mapped STR associations identified by our pipeline have, to our knowledge, not been previously reported. For example, this set includes positive associations between the length of a highly polymorphic CGG repeat in the promoter of the gene *CBL* and both platelet count (p=4e-83) and platelet crit (p=6e-103; 11^th^ most-significant platelet crit peak; **Supplementary Table 11**; **Fig. 4a-b**; **Supplementary Fig. 20**). This finding fits with observations at other loci, as CG-rich repeats in promoter and 5’ UTR regions have been strongly implicated in transcriptomic regulation^13^, often via epigenomic regulation^43,44^. This repeat in *CBL* is also confidently fine-mapped to an association with mean sphered cell volume (p=7e-16; **Supplementary Fig. 21**), but that is comparatively weaker and we do not discuss it here. For both the platelet crit and platelet count phenotypes, SuSiE and FINEMAP identify two genome-wide significant signals in this region, one of which they both localize to this STR. After conditioning on a lead variant from the other signal (rs2155380), this STR becomes the lead variant in the region by a wide margin (**Fig. 4c-d**). Conditioning on both rs2155380 and this STR accounts for all the signal in the region (**Fig. 4e**). This supports the fine-mappers’ prediction that there are two signals in this region, one of which is driven by this STR.

**Figure 4:**
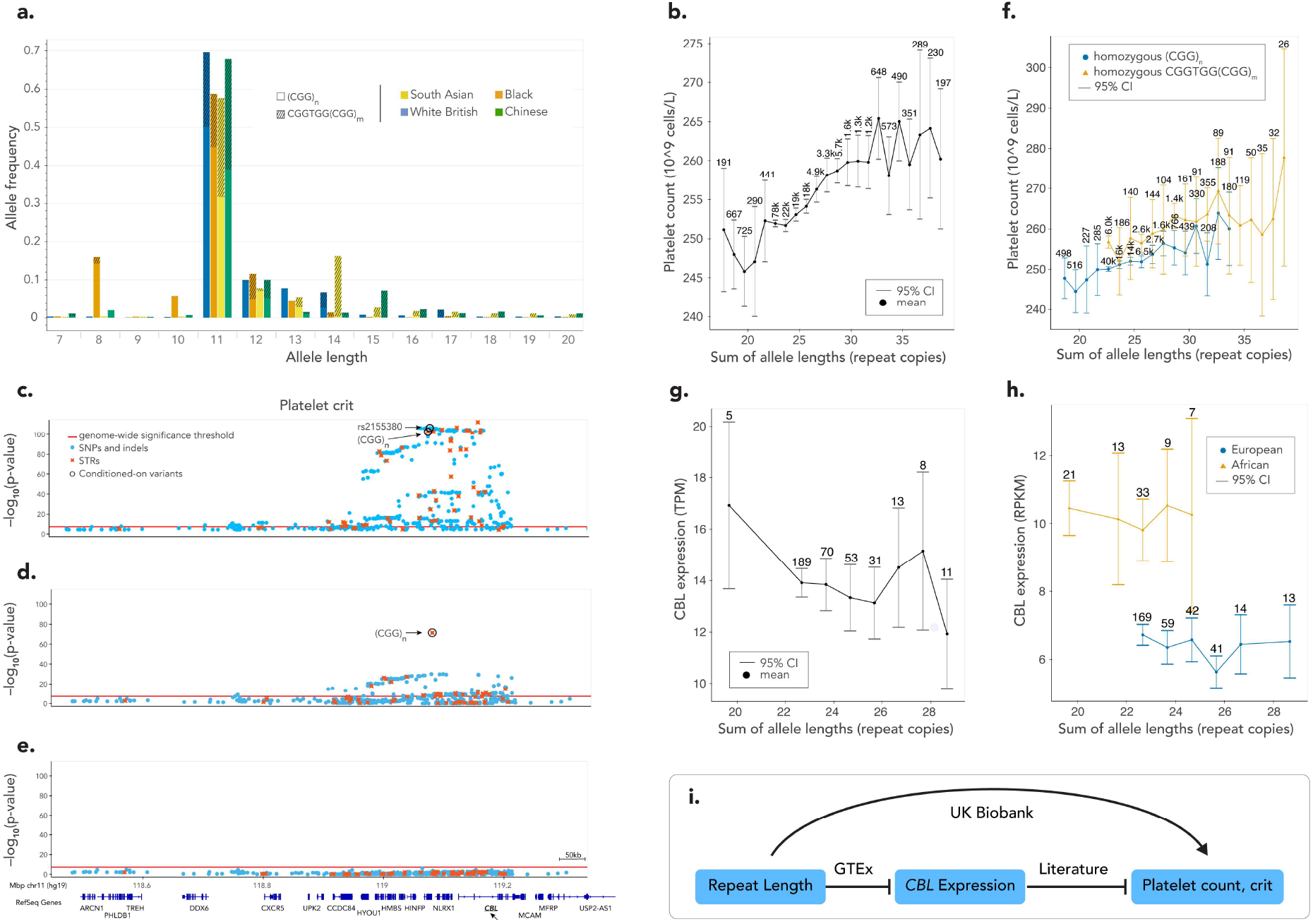
A highly polymorphic CGG repeat in the promoter of *CBL* influences platelet traits. **(a) Distribution of STR alleles across populations.** The x-axis gives STR length (in number of full repeat unit lengths, using WGS data) and y-axis gives the population frequency. The hatched portion of each bar corresponds to the alleles of that length that include a “TGG” imperfection at the second repeat (rs7108857). The ultra-rare alleles with T imperfections at other locations we label as ‘perfect’ for the sake of these analyses. Colors denote different UKB populations. Extreme allele lengths 3-6, 21-33, 36 and 37 each have frequency less than 1% in all populations and have been omitted. **(b) STR length vs. platelet count.** STR length sums were calculated from WGS data on (potentially related) White British participants that passed quality control. Error bars correspond to 95% confidence intervals. Only allele length sums with a population frequency of 0.1% or greater are displayed. Rounded population-wide counts are displayed for each sum. **(c-e) Association of variants at the *CBL* locus with platelet crit.** Association plots in the White British population are shown before conditioning **(c)**, after conditioning on rs2155380 **(d)**, and after conditioning on both rs2155380 and STR length **(e)**. Light blue=SNPs and indels; orange=STRs. Red line=genome-wide significance threshold, black circles=the (CGG)_n_ STR and rs2155380. **(f) STR length vs. platelet count conditioned on the TGG imperfection rs7108857.** STR length sums were calculated from WGS data on (potentially related) White British individuals that passed quality control. Blue=individuals homozygous for no imperfection (n=86,974); orange=individual homozygous for the imperfection (n=11,778). For each category, only length sums with a frequency of 0.2% or greater in that category are displayed. **(g-h) STR length vs. *CBL* expression.** Associations are shown for Cultured Fibroblasts from GTEx (n=393) **(g)** and LCLs from Geuavdis (n=447) **(h).** Yellow=African, blue=European. For **(g-h)** only allele length sums with at least 5 corresponding participants are displayed. **(i) Proposed pathway for effect of STR length on platelet traits.** The arrow denotes a positive association, the capped lines denote negative associations. Interactions are captioned by their information sources.

This STR contains a common imperfection (rs7108857, which changes the second CGG copy to TGG). That variant is in weak linkage disequilibrium (LD) with the length of the STR (r^2^ in imputed genotypes ranging between 0.023 (White British) and 0.175 (Chinese)) (**Fig. 4a**) and in strong LD with the lead variant of the other signal in this region (rs2155380, White British r^2^=97.8%). While rs7108857 is more strongly associated with the platelet traits than the STR’s length (platelet count p=9e-86, platelet crit p=4e-98) and is associated with CBL expression in the GTEx cohort (minimum p=2.04e-18 in Esophagus Muscularis), given the fine-mappers’ results that the STR length association is an independent signal, it is unsurprising that the STR-length association remains after stratifying on the presence of this imperfection (**Fig. 4f**). This suggests that imperfections and repeat lengths are different characteristics of repeats that may have distinct associations.

The imputation of this STR displays more modest levels of concordance with WGS data than less polymorphic loci (r^2^=0.582 between length dosages from imputation and summed lengths from WGS; additional statistics reported in **Supplementary Table 5**). Yet reassuringly, hard-called genotypes from WGS show similar trends with both platelet traits (**Fig. 2b**, **Supplementary. 20a,c-d**). Further, population-specific allele length distributions at this STR observed in UKB are highly concordant with those observed in the 1000 Genomes Project (**Fig. 4a**, **Supplementary Fig. 22**).

*CBL* codes for a protein in the RING finger subfamily of E3 ubiquitin ligases – a class of proteins, each with specific target molecule(s), that ubiquitinate their targets and thus prime them for downstream degradation. CBL targets the thrombopoietin receptor MPL^45^, thereby downregulating thrombopoietin signaling^46^. As thrombopoietin is the primary positive regulator of platelet production^47^, this implicates *CBL* as a negative regulator of platelet production. As further evidence of this link, controlled experiments in mice demonstrate that loss of *CBL* function in megakaryocytes, the bone marrow platelet progenitor cells, results in increased platelet counts^48^. Further, we observed that increased CCG length was negatively associated with *CBL* expression in three tissues in the GTEx cohort^37^ (each with p-value < 0.05 after multiple hypothesis correction; **Supplementary Table 12**; **Fig. 4g**). This association replicated (p=0.007) in European individuals in the Geuvadis cohort^49^ (**Fig. 4h**). This data fits with the STR association with platelet traits and the connection between *CBL* and platelet production, leading to an overall hypothesis that longer CCG repeat alleles contribute to increased platelet count by decreasing CBL expression (**Fig. 4i**).

### Additional examples of confidently fine-mapped STR-trait associations

We observe another 5’ UTR CCG repeat in *BCL2L11* (also known as *BIM*) that is confidently fine-mapped to eosinophil percentage (p=6e-75) and eosinophil count (p=5e-58) (**Supplementary Table 11**). This repeat is the most strongly associated variant in the region for both traits, and conditioning on it accounts for the entire signal in this region (**Supplementary Fig. 23**). Proteins in the BCL-2 family are known to act as anti-or pro-apoptotic regulators. *BCL2L11* in particular is pro-apoptotic and is required in the tightly regulated lifespan of myeloid lineage cells^50^, which include eosinophils. One mouse-model study showed that loss of repression of *BCL2L11* is associated with lowered eosinophil counts^51^ and another showed that *BCL2L11* knockout increases counts of granulocytes, a class of cells including eosinophils^52^. This implicates *BCL2L11* in the regulation of eosinophil count, supporting the connection we observe between eosinophil count and the length of this STR.

While exonic repeats are potentially easier to interpret, a majority of our confidently fine-mapped STRs fall in intronic regions. We resolve one of the strongest signals for mean platelet volume (p<1e-300; one of twelve such peaks with p-values exceeding our numeric precision) to a multi-allelic poly-A STR in an intron of the gene *TAOK1* (**Supplementary Table 11**; **Supplementary Fig. 24a**). Conditioning on the length of this STR demonstrates that it explains the majority of the signal in this region (**Supplementary Fig. 24b**). The same STR also shows a strong association with platelet count, with p=2e-181 and a SuSiE CP of 1.

*TAOK1* is a protein kinase that plays a role in regulating microtubule dynamics^53^ and microtubule function is known to be critical to platelet generation^54^. The STR is in an intron of the canonical *TAOK1* transcript but lies immediately downstream of a non-protein coding transcript of *TAOK1* (ENST00000577583; which contains a retained intron) and is approximately 2.4kb upstream of a differentially spliced exon. The STR also bears the hallmarks of a regulatory element: it is located in a DNase hypersensitivity cluster and overlaps a transcription factor binding site for ESR1 (**Methods**). Although this STR was filtered from our initial GTEx callset due to low call rate (11%), we used our haplotype panel to impute it into available SNP data for that cohort (**Methods**). While we did not identify significant associations between repeat length and splicing of any nearby exons, STR lengths showed modest but significant negative correlation with *TAOK1* expression in 5 tissues (strongest p-value 8e-6 in thyroid; **Supplementary Fig. 24c-d**). The repeat also showed associations with the expressions of nearby genes *ANKRD13B* (negative association; strongest p-value 5e-05 in heart) and *TP53I13* (positive association; strongest p-value 6e-5 in tibial nerve), although their potential role in platelet regulation is less clear.

In another confidently fine-mapped example we identify a previously unreported association between a GTTT repeat in an intron of estrogen receptor beta (*ESR2*) and haemoglobin concentration (p=1e-24), red blood cell count (p=3e-24) and haematocrit (p=1e-26), where additional repeat copies correspond to lower measurements of all three traits (**Supplementary Table 11**; **Supplementary Note 5; Supplementary Fig. 25**). Despite the relatively weak discovery p-value and differing allele distributions between the White British and Black populations (**Supplementary Fig. 25c**), all three associations replicate in the Black population with p-values < 0.05. Consistent with these associations, *ESR2* ligand 17β-estradiol has been implicated in the regulation of red blood cell production^55,56^. We found a significant negative association between STR length and *ESR2* expression in two tissues in GTEx (each with p-value < 0.05 after multiple hypothesis correction; **Supplementary Table 12**). While evidence suggests a link between *ESR2* and red blood cell production, the expected direction of effect is unclear given the highly tissue-specific isoform usage and functions of this gene (**Supplementary Note 5**). Nevertheless, our results support a role of this STR in red blood cell production through regulation of *ESR2*.

We observed many additional associations of interest amongst the confidently fine-mapped STRs. For example, we find multiple AC repeats in our set that are significantly associated with expression of nearby genes. This includes a polymorphic AC repeat located in the 3’ UTR of *NCK2* which is associated with mean platelet volume (p=7e-29; **Supplementary Table 11; Supplementary Fig. 26**). This repeat overlaps a binding site for the transcription factor PABPC1 and has a significant negative association with *NCK2* expression in multiple GTEx tissues (strongest p=5e-7; **Supplementary Table 12**). We also find a highly polymorphic CCG repeat in the 5’ UTR of *RHOT1* that is associated with red blood cell distribution width (p=7e-13; **Supplementary Table 11**). WGS data shows our imputation of this locus is poor, which is unsurprising given the highly polymorphic nature of this STR. Nonetheless, the effect of this STR is biologically plausible – this repeat overlaps a CTCF binding site, is located within a nucleosome depleted region of a H3K27ac peak in LCLs, and shows a strong association with the expression of *RHOT1* in LCLs from the Geuvadis dataset (p=2e-44 in Europeans, p=0.035 in Africans; **Supplementary Fig. 27**). Finally, many STRs in our fine-mapped set consist of poly-A repeats. While traditionally these have been particularly challenging to genotype^57^, many such STRs, including poly-A repeats in *MYO9B*, *DENND4A*, and *NRG4*, show strong statistical evidence of causality and replicate across multiple population groups (**Fig. 2**). Taken together, these loci exemplify the large number of confidently fine-mapped STRs our analysis provides for future study.

## Discussion

In this study, we imputed 445,720 STRs into the genomes of 408,153 participants in the UK Biobank and associated their lengths with 44 blood cell and other biomarker traits. Using fine-mapping, we estimate that STRs account for 5.2-7.6% of causal variants for these traits that can be identified by GWAS. We stringently filtered the fine-mapping output to produce 119 confidently fine-mapped STR-trait associations with strong evidence for causality across 93 distinct STRs including some of the strongest signals for apolipoprotein B, platelet crit and mean platelet volume. These confidently fine-mapped STRs replicated in the Black, South Asian and Chinese UKB populations at higher rates than non-fine-mapped STRs (p<0.02 in each). A subset of these STRs were associated with expression of nearby genes, providing evidence for their impact on regulatory processes and explanations for their effects on the studied traits.

Broadly, our study highlights the importance of considering a more complete set of genetic variants in complex trait analysis. It has been proposed that STRs may represent an important source of the “missing heritability” associated with SNP-based GWAS^58,59^. Indeed, STRs, as well as other complex variant types such as VNTRs^2^, copy number variants^4^, HLA types^60^, and some structural variants^61^ are often highly multi-allelic and only imperfectly tagged by individual common SNPs. This suggests that omitting these variants from analysis pipelines may overlook important sources of causal variants and heritability. Further, we expect incorporation of this additional source of causal variants, which we observe often exhibit population-specific allele distributions, will improve downstream applications such as polygenic risk scores, particularly in constructing scores that are more applicable across diverse populations.

While our results uncover many novel candidate causal STR variants, these findings are not exhaustive. Our fine-mapping procedure was exceptionally conservative and excluded hundreds of STR-trait associations strongly predicted to be causal in some but not all settings tested. Further, whereas here we performed association tests with a fixed effects model, analysis with a linear mixed model would increase power to detect additional associations. Additionally, compute constraints limited our analysis to a small number of traits. In follow up studies we plan to extend this analysis to a wide variety of medically-actionable traits.

Another limitation of our study is that it is based on imputed genotypes. Our SNP-STR reference panel only included 27.5% of the 1.6 million STRs in the original HipSTR reference panel (**URLs**), due to the exclusion of the many STRs with low heterozygosity, but also due to the exclusion of non-autosomal STRs, most long repeats such as those implicated in pathogenic expansion disorders, and many STR alleles that are common only in non-European populations^30^. Further, imputed genotypes are inherently subject to some amount of noise, especially in non-European populations. Despite these limitations, analysis of WGS data released for 200,025 UKB participants^62^ during the course of this study validated associations seen in imputed data. Subsequently, popSTR calls at 2.5 million STRs were released for 150,000 of the participants with WGS data^62^. Future studies which perform STR-based GWAS solely using WGS datasets such as this should ameliorate the limitations mentioned above, enabling many future discoveries.

Methodological advances are also needed to support the study of STRs. Here we developed associaTR, an open-source reproducible pipeline that enables future studies to conduct STR length-based association tests. However, we envision that integrating support for STR length-based tests and other complex variant associations into widely used GWAS toolkits would enable more routine analysis of the full spectrum of human genetic variation. Improvements to our association testing models are also likely to reveal new insights. In this study we only modeled linear associations between STR lengths and trait values. Visualization of some of the associations we identify suggests that these linear models only partially approximate those signals. Many STR effects may be best described by non-linear models, such as quadratic or sigmoid relationships between repeat copy numbers and traits. Yet fitting non-linear models would require modeling the effects of the two alleles at each locus separately while simultaneously controlling for the possibility of over-fitting, and is a topic of ongoing work. We also only tested traits for association with the number of repeat units. However, inspection of individual loci reveals that complex repeat structures are common (**Table 1**). Systematic evaluation of the potential for epistasis between repeat imperfections and STR lengths, as well as between the lengths of neighboring repeats, would potentially enable better understanding of the phenotypic impact of STRs.

Importantly, our results highlight current challenges in performing statistical fine-mapping. We found that fine-mapping results were in some cases highly sensitive to the choice of fine-mapping tool, and to a lesser extent to data-processing choices and fine-mapper instabilities, where one fine-mapping run would identify a variant as highly likely to be causal but a second would identify it as having no causal impact. Further, our simulations suggest that fine-mappers have low sensitivity rates even when sample sizes are large and all model assumptions are met. This suggests statistical fine-mapping results should be interpreted with caution and evaluated for sensitivity to model choices, and that further work is needed to make the process of fine-mapping more robust.

Although fine-mapping inconsistencies were identified for SNPs and indels as well as STRs, they were most prevalent for STRs. While this may in part be due to issues with imputing STR genotypes, more research is needed to further evaluate the performance of current fine-mapping tools on regions containing STRs. Additionally, there is a need for fine-mapping tools that can model effects of multi-allelic variants. Existing fine-mapping frameworks in theory can accurately model linear repeat-length associations, but we hypothesize that more detailed modeling of LD between SNPs and individual STR alleles may enable more accurate model fitting procedures. Similarly, during model fitting, existing tools often compare models which trade one causal variant for another variant in close LD, but greater accuracy may be obtained by comparing models which trade off a single, potentially causal, multi-allelic variant for multiple simultaneously-causal bi-allelic variants.

Overall, our study provides a statistical framework for incorporating hundreds of thousands of tandem repeat variants into GWAS studies, either via imputation, or using STR genotypes called from WGS such as the newly released^62^ UKB callset. Our study identifies dozens of novel candidate variants for future mechanistic studies and demonstrates that STRs likely make a widespread contribution to complex traits.

## Methods

### Selection of UK Biobank participants

We downloaded the fam file and sample file for version 2 of the phased SNP array data (referred to in the UKB documentation as the ‘haplotype’ dataset) using the ukbgene utility (ver Jan 28 2019 14:09:15 - using Glibc2.28(stable)) described in UKB Data Showcase Resource ID 664 (**URLs**). The IDs from the sample file already excluded 968 individuals previously identified as having excessive principal component-adjusted SNP array heterozygosity or excessive SNP array missingness after call-level filtering^31^ indicating potential DNA contamination. We further removed withdrawn participants, indicated by non-positive IDs in the sample file as well as by IDs in email communications from the UKB access management team. After the additional filtering, data for 487,279 individuals remained.

We downloaded the sample quality control (QC) file (described in the sample QC section of UKB Data Showcase Resource ID 531 (**URLs**)) from the European Genome-Phenome Archive (accession EGAF00001844707) using pyEGA3^63^. We subsetted the non-withdrawn individuals above to the 408,870 (83.91%) participants identified as White-British by column `in.white.British.ancestry.subset` of the sample QC file. This field was computed by the UKB team to only include individuals whose self-reported ethnic background was White British and whose genetic principal components were not outliers compared to the other individuals in that group^31^. In concordance with previous analyses of this cohort^31^ we additionally removed data for:

● 2 individuals with an excessive number of inferred relatives, removed due to plausible SNP array contamination (participants listed in sample QC file column `excluded.from.kinship.inference` that had not already been removed by the UKB team prior to phasing)
● 308 individuals whose self-reported sex did not match the genetically inferred sex, removed due to concern for sample mislabeling (participants where sample QC file columns `Submitted.Gender` and `Inferred.Gender` did not match)
● 407 additional individuals with putative sex chromosome aneuploidies removed as their genetic signals might differ significantly from the rest of the population (listed in sample QC file column `putative.sex.chromosome.aneuploidy`)

Following these additional filters the data for 408,153 individuals remained (99.82% of the White British individuals considered above).

### SNP and indel dataset preprocessing

We obtained both phased hard-called and imputed SNP and short indel genotypes made available by the UKB. These variants were provided in reference genome hg19 coordinates, and all analyses in this study, unless otherwise specified, were performed with hg19 coordinates.

#### Phased hard-called genotypes

We downloaded the bgen files containing the hard-called SNP and indel haplotypes (release version 2) and the corresponding sample and fam files using the ukbgene utility (UKB Data Showcase Resource 664 (**URLs**)). These variants had been genotyped using microarrays and phased using SHAPEIT3^64^ with the 1000 genomes phase 3 reference panel^23^. Variants genotyped on the microarray were excluded from phasing and downstream analyses if they failed QC on more than one microarray genotyping batch, had overall call-missingness rate greater than 5% or had minor allele frequency less than 0.01%. Of the resulting 658,720 variants, 99.5% were single nucleotide variants, 0.2% were short indels (average length 1.9bp, maximal length 26bp), and 0.2% were short deletions (average length 1.9bp, maximal length 29bp).

#### Imputed genotypes

We similarly downloaded imputed SNP data using the ukbgene utility (release version 3). Variants had been imputed with IMPUTE4^31^ using the Haplotype Reference Consortium panel^22^, with additional variants from the UK10K^24^ and 1000 Genomes phase 3^23^ reference panels. The resulting imputed variants contain 93,095,623 variants, consisting of 96.0% single nucleotide variants, 1.3% short insertions (average length 2.5bp, maximum length 661bp), 2.6% short deletions (average length 3.1bp, maximum length 129bp). This set does not include the 11 classic human leukocyte antigen alleles imputed separately.

We used bgen-reader^65^ 4.0.8 to access the downloaded bgen files in python. We used plink2^33^ v2.00a3LM (build AVX2 Intel 28 Oct 2020) to convert bgen files from both hard-called and imputed SNPs to the plink2 format for downstream analyses. For hard-called genotypes, we used plink to set the first allele to match the hg19 reference genome. Imputed genotypes already matched the reference. Unless otherwise noted, our pipeline worked with imputed genotypes as non-reference allele dosages, i.e. Pr(heterozygous) + 2 ∗ Pr(homozygous alternate) for each individual.

### STR imputation

We previously published a reference panel containing phased haplotypes of SNP variants alongside 445,720 autosomal STR variants in 2,504 individuals from the 1000 Genomes Project^23,30^ (**URLs**). This panel focuses on STRs ascertained to be highly polymorphic and well-imputed in European individuals. Notably, this excludes many STRs known to be implicated in repeat expansion diseases, STRs that are primarily polymorphic only in non-European populations, or STRs that are too mutable to be in strong linkage disequilibrium (LD) with nearby SNPs.

The IDs listed in the ‘str’ column of Supplementary Table 2 at that URL describe which variants in the reference panel are STRs and which are other types of variants. That produces a list of 445,715 unique variant IDs and 5 IDs which are each assigned to four separate variants in the reference panel VCFs. For the IDs with multiple assignments, we selected the variant that appeared first in the VCF and discarded the others, leaving 445,720 unique STR variants each with unique IDs.

While our analyses with these STRs were performed using hg19 coordinates unless otherwise stated, we also provide hg38 reference coordinates for these STRs in the supplementary tables. We obtained those coordinates using liftover (**URLs**) which resulted in identical coordinates as in HipSTR’s^6^ hg38 STR reference panel (**URLs**). All STRs successfully lifted over to hg38 coordinates.

To select shared variants for imputation, we note that 641,582 (97.4%) of SNP and indel variants that were hard-called and phased in the UKB participants were present in our SNP-STR reference panel. As a quality control step, we filtered variants that had highly discordant minor allele frequencies between the 1000 Genomes European subpopulations (**URLs**) and White British individuals from the UKB. We first took a maximal unrelated set of the White British individuals (see **Phenotype Methods** below) and then visually inspected the alternate allele frequency of the overlapping variants (**Supplementary Fig. 1**) and chose to remove the 110 variants with an alternate allele frequency difference of more than 12%.

We used Beagle^32^ v5.1 (build 25Nov19.28d) with the tool’s provided human genetic maps (**URLs**) and non-default flag `ap=true` to impute STRs into the remaining 641,472 SNPs and indels from the SNP-STR panel into the hard-called SNP haplotypes. Though we performed the above comparison between reference panel Europeans and UKB White British individuals, we performed this STR imputation into all UKB participants using all the individuals in the reference panel. We chose Beagle because it can handle multi-allelic loci. Due to computational constraints, we ran Beagle per chromosome on batches of 1000 participants at a time with roughly 18GB of memory. We merged the resulting VCFs across batches and extracted only the STR variants. Lastly, we added back the `INFO` fields present in the SNP-STR reference panel that Beagle removed during imputation.

Unless otherwise noted, our pipeline worked with these genotypes as length dosages for each individual, defined as the sum of length of each of the two alleles, weighted by imputation probability. Formally, *dosage* = Σ_*a* ∈*A*_ *len*(*a*) ∗ [*Pr*(*hap*_1_ == *a*) + *Pr*(*hap*_2_ == *a*)], where *A* is the set of all possible STR alleles at the locus, *len*(*a*) is the length of allele *a*, and *Pr*(*hap_i_* == *a*) is the probability that the allele on the *i*th haplotype is *a*, output by Beagle in the `AP1` and `AP2 FORMAT` fields of the VCF file.

Estimated allele frequencies (**Fig. 1b**) were computed as follows: for each allele length *L* for each STR, we summed the imputed probability of the STR on that chromosome to have length *L* over both chromosomes of all unrelated participants. That sum is divided by the total number of chromosomes considered to obtain the estimated frequency of each allele.

### Inferring repeat units

Each STR in the SNP-STR reference panel was previously annotated with a repeat period - the length of its repeat unit - but not the repeat unit itself. We inferred the repeat unit of each STR in the panel as follows: we considered the STR’s reference allele and given period. We then took each k-mer in the reference allele where k is the repeat period, standardized those k-mers, and took their counts. We define the standardization of a k-mer to be the sequence produced by looking at all cyclic rotations of that k-mer and choosing the first one lexicographically. For example, the standardization of the k-mer CAG would be AGC. If the most common standardized k-mer was less than twice as frequent as the second most common standardized k-mer, we did not call a repeat unit for that STR (11,962 STRs; 2.68%). Otherwise, the most common standardized k-mer was labeled as the forward-strand (based on the reference genome) repeat unit for that STR. To infer the strand-independent repeat unit for the STR we looked at all rotations of the forward-strand repeat unit in both the forward and reverse-complement directions and chose whichever comes first lexicographically. For example the repeat unit for the STR TGTGTGTG would be AC, while the forward-strand repeat unit would be GT. In the large majority of cases the repeat unit identified by this approach is the unit which is duplicated or deleted in alternate alleles, but this method of identifying repeat units does not consider alternate alleles and so does not make that guarantee.

### Phenotypes and covariates

IDs listed in this section refer to the UKB Data Showcase (**URLs**).

We analyzed a total of 44 blood traits measured in the UKB. 19 phenotypes were chosen from Category Blood Count (Data Field ID 100081) and 25 from Category Blood Biochemistry (Data Field ID 17518). We refer to them as blood cell count and biomarker phenotypes respectively. The blood cell counts were measured in fresh whole blood while all the biomarkers were measured in serum except for glycated haemoglobin which was measured in packed red blood cells (details in Resource ID 5636). The phenotypes we analyzed are listed in **Supplementary Table 1**, along with the categorical covariates specific to each phenotype that were included during association testing.

We analyzed all the blood cell count phenotypes available except for the nucleated red blood cell, basophil, monocyte, and reticulocyte phenotypes. Nucleated red blood cell percentage was omitted from our study as any value between the bounds of 0% and 2% was recorded as exactly either 0% or 2% making the data inappropriate for study as a continuous trait. Nucleated red blood cell count was omitted similarly. Basophil and monocyte phenotypes were omitted as those cells deteriorate significantly during the up-to-24-hours between blood draw and measurement. This timing likely differed consistently for different clinics, and different clinics drew from distinct within-White British ancestry groups, which could lead to confounding with true genetic effects. See Resource ID 1453 for more information. Reticulocytes were excluded from our initial pipeline. This left us with 19 blood cell count phenotypes. For each blood cell count phenotype we included the machine ID (1 of 4 possible IDs) as a categorical covariate during the association tests to account for batch effects.

Biomarker measurements were subject to censoring of values below and above the measuring machine’s reportable range (Resource IDs 1227, 2405). **Supplementary Table 1** includes the range limits and the number of data points censored in each direction. Five biomarkers (direct bilirubin, lipoprotein(a), oestradiol, rheumatoid factor, testosterone) were omitted from our study for having >40,000 censored measurements across the population (approximately 10% of all data), since those would require analysis with models that take censoring into account. The remaining biomarkers had less than 2,000 censored measurements. We excluded censored measurements for those biomarkers from downstream analyses as they consisted of a small number of data points. For each serum biomarker we included aliquot number (0-3) as a categorical covariate during association testing as an additional step to mediate the dilution issue (described in Resource ID 5636). Glycated haemoglobin was not subject to the dilution issue, being measured in packed red blood cells and not serum, so no aliquot covariate was published in the UKB showcase or included in our analysis.

For each phenotype we took the subset of the 408,153 individuals above that had a measurement for that phenotype during the initial assessment visit or the first repeat assessment visit, preferentially choosing the measurement at the initial assessment when measurements were taken at both visits. We include a binary categorical covariate in association testing to distinguish between phenotypes measured at the initial assessment and those measured at the repeat assessment. Each participant’s age at their measurement’s assessment was retrieved from Data Field ID 21003.

The initial and repeat assessment visits were the only times the biomarkers were measured. The blood cell count phenotypes were additionally measured for those participants who attended the first imaging visit. We did not use those measurements and for each phenotype excluded the <200 participants whose only measurement for that phenotype was taken during the first imaging visit as we could not properly account for the batch effect of a group that small (**Supplementary Table 1**).

No covariate values were missing. Before each association test we checked that each category of each categorical covariate was obtained by at least 0.1% of the tested participants. We excluded the participants with covariate values not matching this criterion, as those quantities would be too small to properly account for batch effects. In practice, this meant that for each biomarker phenotype we excluded the <100 participants that were measured using aliquot 4, and that for 8 of the biomarker phenotypes we additionally excluded the ≤125 participants that were measured using aliquot 3 (**Supplementary Table 1**).

For each phenotype we then selected a maximally-sized genetically unrelated subset of the remaining individuals using PRIMUS^66^ v1.9.0. When multiple such maximal subsets existed (for instance, wherever a single individual needed to be chosen from a family of two), one subset was chosen arbitrarily, thus introducing some lack of reproducibility. Precomputed measures of genetic relatedness between participants (described in UKB paper supplement section 3.7.1^31^) were downloaded using ukbgene (Resource ID 664). We ran PRIMUS with non-default options `- -no_PR -t 0.04419417382` where the `t` cutoff is equal to 0.5^9^, chosen so that two individuals are considered to be related if they are relatives of third degree or closer. This left between 304,658 and 335,585 unrelated participants per phenotype (**Supplementary Table 1**).

Sex and ancestry principal components (PCs) were included as covariates for all phenotypes. Participant sex was extracted from fam file available with hard called genotypes (see above). The top 40 ancestry PCs were extracted from the corresponding columns of the sample QC file (see the Participants **Methods** section above).

We then rank-inverse-normalized phenotype values for association testing. The remaining unrelated individuals for each phenotype were ranked by phenotype value from least to greatest (ties broken arbitrarily) and the phenotype value for association testing for each individual was taken to be 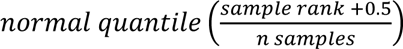. We use rank-inverse normalization as it is standard practice, though it does not have a strong theoretical foundation^67^ and only moderate empirical support^68–71^.

For each phenotype and its remaining unrelated individuals we standardized all covariates to have mean zero and variance one for numeric stability.

### Association testing

We performed STR and SNP association testing separately. We developed associaTR to streamline performing association tests between STR length and quantitative traits. While our approach relies on a standard linear model, linear mixed models based on STR length dosages would likely result in increased power and will be considered in future studies. As our downstream analyses required STR and SNP associations to be comparable, we also used a standard linear model for SNP association testing.

For STR association testing, the imputed VCFs produced by Beagle were accessed in python with cyvcf2^72^ 0.30.14 and v4.2.1 of our TRTools library^73^. In line with plink’s recommendation for SNP GWAS (**URLs**), 6 loci with non-major allele dosage < 20 were filtered. For each STR, we fit the linear model 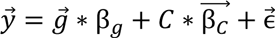 where 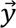 is the vector of rank-inverse-normalized phenotype values per individual, 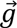 is the vector of STR length dosage genotypes per individual, β*_g_* is the effect size of this STR, *C* is the matrix of standardized covariates, 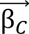 is the vector of covariate effect sizes, and 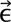 is the vector of errors between the model predictions and the outcomes. Models were fit using the `regression.linear_model.OLS` function of the Python statsmodels library v0.13.2 (**URLs**). Per GWAS best-practices, we used imputation dosage genotypes instead of best-guess genotypes^74^.

We used plink2^33^ v2.00a3LM (build AVX2 Intel 28 Oct 2020) for association testing of imputed SNPs and indels. For each analysis, plink first converts the input datasets to its pgen file format. To avoid performing this operation for every invocation of plink, we first used plink to convert the SNP and indel bgen files to pgen files a single time. We invoked plink once per chromosome per phenotype. We used the plink flag `--mac 20` to filter loci with minor allele dosage less than 20 (**URLs**). Plink calculates minor allele counts across all individuals before subsetting to individuals with a supplied phenotype, so this uniformly filtered 22,396,837 (24.1%) of the input loci from each phenotype’s association test leaving 70,698,786 SNPs and indels. Plink fit the same linear model described above in the STR associations, except that 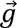 is the vector of dosages of the non-reference SNP or indel allele.

For conditional regressions, we fit the model 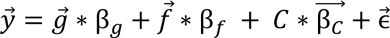 where all the terms are as described above, except 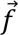 is the vector of per-individual genotypes of the variant being conditioned on, and β*_f_* is its effect size.

### Comparison with Pan-UKB pipeline

We compared the results of our pipeline to results available on the Pan UKBB^34^ website (see **URLs**) using bilirubin as an example trait. We matched variants between datasets on chromosome, position, reference and alternate alleles, excluding variants not present in both pipelines. We found our pipeline produced largely similar but somewhat less significant p-values than those reported for European participants in Pan UKBB (**Supplementary Fig. 2**).

### Defining significant peaks

Given a peak width *w* (bp), we selected variants to center peaks on in the following manner:

1. Order all variants (of all types) from most to least significant. For variants which exceed our pipeline’s precision (p<1e-300), order them by their chromosome and base pair from first to last. (These variants will appear at the beginning of the list of all variants).
2. For each variant: If the variant has p-value ≥ 5e-8, break. If there is a variant in either direction less than *w* bp away which has a lower p-value, continue. Otherwise, add this variant to the list of peak centers.

We define peaks to be the *w* (base pair) width regions centered on each selected variant. The statistics given in the **Results** are calculated using *w* = 250*kb*. The identification of peaks in **Fig. 1c-d** was made with *w* = 20*mb* for visualization purposes. Note that peaks within *w* bp of the end of a chromosome will necessarily be smaller than *w* bp in width.

#### Identifying indels which are STR alleles

Some STR variant alleles are represented both as alleles in our SNP-STR reference panel and as indel variants in the UKB imputed variants panel. We excluded the indel representations of those alleles from fine-mapping, as they represent identical variants and could confound the fine-mapping process. For each STR we constructed the following interval:

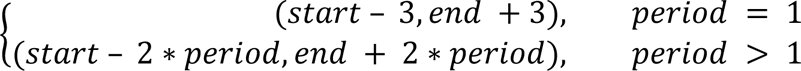

where *period* is the length of the repeat unit. and *start* and *end* give the coordinates of the STR in base pairs. We call an indel an STR-indel if it only represents either a deletion of base pairs from the reference or an insertion of base pairs into the reference (not both), overlaps only a single STR based on the interval above, and represents an insertion or deletion of full copies of that STR’s repeat unit. We conservatively did not mark any STR-indels for STRs whose repeat units were not called (see above) or for which the insertion or deletion was not a whole number of copies of any rotation of the repeat unit.

### Fine-mapping

For each phenotype, we selected contiguous regions to fine-map in the following manner:

1. Choose a variant (SNP or indel or STR) with p-value < 5e-8 not in the major histocompatibility complex (MHC) region (chr6:25e6-33.5e6).
2. While there is a variant (SNP or indel or STR) with p-value < 5e-8 not in the MHC region and within 250kb of a previously chosen variant, include that variant in the region and repeat.
3. This fine-mapping region is (min variant bp – 125kb, max variant bp + 125kb).
4. Start again from step 1 to create another region, starting with any variant with p-value < 5e-8 not already in a fine-mapping region.

This is similar to the peak selection algorithm above but is designed to produce slightly wider regions so that we could fine-map nearby peaks jointly. We excluded the MHC because it is known to be difficult to effectively fine-map. Note that peaks within 125kb of the end of a chromosome will necessarily be smaller than the minimum 125kb width in that direction.

This produced 14,494 trait-regions. Due to computational challenges during fine-mapping (see below), we excluded three regions (urate 4:8165642-11717761, total bilirubin 12:19976272-22524428 and alkaline phosphatase 1:19430673-24309348) from downstream analyses (see below), leaving 14,491 trait-regions.

We used two fine-mapping methods to analyze each region:

*SuSiE*^35^: For each fine-mapping trait-region, for each STR and SNP and indel variant in that region that was not filtered before association testing, was not an STR-indel variants (see above) and had p-value ≤ 5e-4 (chosen to reduce computational burden), we loaded the dosages for that variant from the set of participants used in association testing for that phenotype. For those regions we also loaded the rank-inverse-normalized phenotype values and covariates corresponding to that phenotype. We separately regressed the covariates out of the phenotype values and out of each variant’s dosages and streamed the residual values to HDF5 arrays using h5py v3.6.0 (**URLs**). We used rhdf5 v2.38.0 (**URLs**) to load the h5 files into R. We used an R script to run SuSiE v0.11.42 on that data with non-default values `min_abs_corr=0` and `scaled_prior_variance=0.005`. `min_abs_corr=0` forced SuSiE to output all credible sets it found so that we could determine the appropriate minimum absolute correlation filter threshold in downstream analyses. We set `scaled_prior_variance` to 0.005 which we considered is a more realistic guess of the per-variant percentage of signal explained than the default of 20%, although we determined that this parameter had no effect on the results (**Supplementary Note 3**). The SuSiE results for some regions did not converge within the default number of iterations (100) or produced the default maximum number of credible sets (10) and all those credible sets seemed plausible (minimum pair-wise absolute correlation ≥ 0.2 or size ≤ 50). We reran those regions with the additional parameters `L=30` (maximum number of credible sets) and `max_iter=500`. No regions failed to converge in under 500 iterations. We re-analyzed several loci that produced 30 plausible credible sets again with `L=50`. No regions produced 50 plausible credible sets. SuSiE failed to finish for two regions (urate 4:8165642-11717761, total bilirubin 12:19976272-22524428) in under 48 hours; we excluded those regions from downstream analyses. A prior version of our pipeline had applied a custom filter to some SuSiE fine-mapping runs that caused SNPs with total minor allele dosage less than 20 across the entire population to be excluded. For consistency, any regions run with that filter which produced STRs included in our confidently fine-mapped set were rerun without that filter. Results from the rerun are reported in **Supplementary Table 4**.

SuSiE calculates credible sets for independent signals and calculates an alpha value for each variant for each signal – the probability that that variant is the causal variant in that signal. We used each variant’s highest alpha value from among credible sets with purity ≥ 0.8 as its casual probability (CP) in our downstream analyses (or zero if it was in no such credible sets). See **Supplementary Note 1**.

*FINEMAP*^36^: We selected the STR and SNP and indel variants in each fine-mapping region that were not filtered before association testing and had p-value < 0.05 (chosen to reduce computational burden). We excluded STR-indels (see above). We constructed a FINEMAP input file for each region containing the effect size of each variant and the effect size’s standard error. All `MAF` values were set to `nan` and the `ref` and `alt` columns were set to `nan` for STRs as this information is not required. We then took the unrelated participants for the phenotype, loaded their dosage genotypes for those variants and saved them to an HDF5 array with h5py v3.6.0 (**URLs**). To construct the LD input file required by FINEMAP, we computed the Pearson correlation between dosages of each pair of variants. We then ran FINEMAP v1.4 with non-default options `--sss --n-causal-snps 20`. In regions which FINEMAP gave non-zero probability to their being 20 causal variants, we reran FINEMAP with the option `–n-causal-snps 40` and used the results from the rerun. FINEMAP did not suggest 40 causal variants in any region. FINEMAP caused a core dump when running on the region alkaline phosphatase 1:19430673- 24309348 so we excluded that region from downstream analyses. (For convenience, for the regions containing no STRs, we directly ran FINEMAP with `--n-causal-snps 40`, unless those regions contained less than 40 variants in which case we ran FINEMAP with `--n-causal-snps` <#variants>).

We used the FINEMAP’s PIP output for each variant in each region as its CP in downstream analyses.

### Alternative fine-mapping conditions

We reran SuSiE and FINEMAP using alternative settings on trait-regions that contained one or more STRs with p-value < 1e-10 and CP ≥ 0.8 in both the original SuSiE and FINEMAP runs. Each new run differed from the original run in exactly one condition. We restricted our set of high-confidence fine-mapped STRs (**Supplementary Table 5**) to those that had p-value < 1e-10 and CP ≥ 0.8 in the original runs and maintained CP ≥ 0.8 in a selected set of those alternate conditions.

For SuSiE, we evaluated using best-guess genotypes vs. genotype dosages as input. For FINEMAP, we tested varying the p-value threshold, choice of non-major allele frequency threshold, effect size prior, number of causal variants per region, and stopping threshold. Additionally, we reran FINEMAP with no changed settings to examine potential FINEMAP instability.

See **Supplementary Note 3** for a more detailed discussion of these various settings and their impact on fine-mapping results.

### Fine-mapping simulations

We simulated phenotypes under additive genetic models and fine-mapped those phenotypes separately at individual regions. Our simulations used real genotypes from White British UKB participants and focused on regions originally identified by our GWAS to maintain realistic LD patterns observed at regions with true signals. We used regions associated with platelet count as it was the phenotype with the maximal number of fine-mapping regions (n=548).

#### Strategies for choosing causal variants and effect sizes

We applied three different strategies for choosing causal variants from these regions and choosing their effect sizes. For each strategy, we simulated phenotypes from those variants, ran SuSiE and FINEMAP on all SNPs and STRs in the region against the simulated phenotypes and determined whether the fine-mappers correctly identified the variants simulated to be causal.

For the first strategy we chose causal SNPs and indels at random, weighting by minor allele frequency (MAF). For this strategy, we did not simulate causal STRs. To begin, we took all SNPs/indels in all platelet count regions that had either FINEMAP CP ≥ 0.5 or SuSiE CP ≥ 0.5 and binned them by MAF (bin boundaries = [0.01%, 0.1%, 10%, 50%]), excluding all variants with MAF < 0.01%. We assigned each bin a relative weight by the proportion of causal variants in that bin vs in all bins as compared to the proportion of all variants in that bin vs all bins, noting that these weights were relatively consistent across bins (within a factor of 2, **Supplementary Table 6**). Using those bin weights, for each fine-mapping region, we then drew causal SNPs/indels at random from all SNPs/indels in the region, with each variant’s chance of being drawn weighted by the bin that its MAF corresponds to. For each bin, we also collected all observed effect sizes of all variants falling in that bin, noting that as expected the effect sizes for common variants were smaller than those for rarer variants (**Supplementary Fig. 6**). For each variant chosen to be causal, we drew an effect size from the corresponding MAF bin. This strategy is designed so that the distributions of MAFs and effect sizes of causal variants in our simulations are similar to those observed for fine-mapped variants for the real phenotype. We repeated this strategy nine times for each simulation region, three times each choosing sets of one, two and three causal variants.

While the first strategy allows for a wide range of simulations by drawing causal variants at random, it may not capture systematic differences between the LD patterns of causal variants and the LD patterns of non-causal variants in causal regions. To address this, for the second strategy we chose variants fine-mapped by SuSiE for platelet count for simulating as causal as these may more closely capture LD patterns of truly causal variants. Specifically, we ran SuSiE on all the SNPs and indels in the fine-mapping region with p < 0.0005 against real platelet count data. Note that by only running SuSiE against the SNP and indel variants in the region, we forced SuSiE to give us the most plausibly causal set of SNPs/indels in the region under the condition that no STRs are causal. We discarded non-pure credible sets (those with variants in less than 0.8 r^2^) as we expect them to be less reliable in identifying truly causal variants. In the 458/548 regions where there were any pure credible sets remaining, we took the top variant from each of the remaining credible sets to use as causal for simulations, using their effect sizes measured against the real platelet count trait as their effect sizes for simulation. For each region, we used its causal variant set to simulate three phenotypes (which are distinct due to different noise terms).

While this second strategy may capture more realistic causal LD patterns compared to choosing causal variants at random, it has the drawback that it relies on the accuracy of fine-mapping to choose the causal variants, which is what we are trying to assess. Strategy one relies on fine-mappers as well, but to a much lesser extent, using them only to identify causal variant MAF and effect size distributions. A second caveat to strategy two is that by restricting to pure credible sets, we likely omit real signals which SuSiE could not resolve well.

For our third strategy, we paralleled our second strategy, except instead of fine-mapping platelet count against only SNPs and indels, we fine-mapped it against all the variants in the region (including STRs), thus allowing it to select STRs as causal for simulation. We continued with simulations as in the second strategy for the 52/548 regions where SuSiE identified a causal STR. This third strategy is the only strategy we performed which simulated causal STRs. As the number of simulations performed with this third strategy was limited, we only use it to contribute briefly to our discussion in the main text.

#### Simulating phenotypes

Let *V* represent the set of causal variants for a region. For each variant *ν* ∈ *V* let 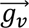 represent a vector of participant genotype dosages and β*_v_* denote the variant’s chosen effect size. Assuming additive and independent contributions of each variant, we simulated a vector of phenotypes (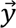) as 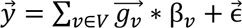, where 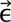∼N(0, diag(1 − 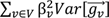)) so that similarly to the real, normalized, phenotypes used for our GWAS, the resulting phenotypes have mean 0 and variance 1.

#### Evaluating fine-mapping on simulated phenotypes

For each simulated phenotype and region we performed association testing of the variants in that region using the same methods as in the main analysis, excepting that we included no covariates and that the phenotypes were not subjected to rank-inverse normalization. We then ran FINEMAP and SuSiE against the variants in the region as described above (in particular, FINEMAP runs were restricted to variants with p<0.05, SuSiE runs to variants with p<0.0005), with the difference that the fine-mapping region was not recalculated from the simulated phenotype GWAS statistics but instead exactly matched to the causal region determined from the platelet count GWAS. Once fine-mappers were run, we calculated STR contribution statistics as for fine-mapping runs on the UKB blood traits (**Supplementary Tables 7-8**).

#### Simulation caveats

Many choices in the design of these simulations affect the interpretation of their results. Notably, these simulated phenotypes make standard assumptions of additive genetic architectures, including no non-linear effects, no epistasis between variants, and that the environmental contribution to each phenotype is both independent of an individual’s genotypes and normally distributed. These simulations also assume that there are no confounding covariates. Additionally, these simulations choose the effect sizes of causal variants from effect sizes calculated in our platelet count GWAS. As effect sizes calculated in the GWAS were measured in mono-variant regressions against platelet count, they will be mis-estimated according to the corresponding variant’s LD to all causal variants in the region in which it resides.

Further, we note that not recalculating the fine-mapping regions may artificially inflate the rate at which strategy one identifies causal variants, as when causal variants in strategy one were randomly chosen to fall near the edges of the region, there would be fewer variants in LD with those variants and fine-mapping them would be easier. This may contribute to the observation in **Supplementary Table 7** that both fine-mappers select STRs in simulation strategy two much more than in simulation strategy one. We also speculate that STRs truly causal for platelet count would contribute to that observation: if those STRs are well tagged by SNPs, strategy two’s run of SuSiE would likely select those tagging SNPs for causal simulation. Then fine-mapping of those simulated phenotypes would have a relatively high chance of confusing those SNPs with the STRs they tag.

Lastly, we observe that FINEMAP mostly identifies variants with low p-values, while a p-value cutoff is necessary for accurate SuSiE results. Once a p-value cutoff is applied, we see that the fine-mappers’ results are almost entirely consistent with one another, in large distinction from how they perform when applied to real datasets, suggesting that there are some features of the architectures of blood traits are not captured by these simulations.

### Validation of imputation results for fine-mapped STRs using WGS data

We worked with WGS CRAM files for 200,025 UKB participants on the UKB Research Analysis Platform cloud solution provided by DNA Nexus. This data was aligned to reference genome hg38. HipSTR was unable to load the index files for the CRAM files of 10 participants, possibly due to file corruption. Removing those participants left us with 200,015 participants. We inadvertently truncated the participant list, leaving 200,000 participants. From that participant list we called genotypes of the 409 STRs in **Supplementary Table 4** using HipSTR^6^ in batches of 500 participants, using the flag `--min-reads 10` and allowing HipSTR to estimate stutter-error models from the data. We merged batches using MergeSTR^73^. We performed call level filtering using DumpSTR^73^ with the flags `--hipstr-min-call-Q 0.9 --hipstr-min-call-DP 10 --hipstr-max-call-DP 10000 --hipstr-min-supp-reads 2 --hipstr-max-call-stutter 0.15 --hipstr-max-call-flank-indel 0.1`. After calling all 200,000 individuals we summarized their genotypes separately per population, noting that 166,638 individuals were in our set of QC’ed (potentially related) White British UKB participants, accounting for 40.8% of the QC’ed White British participants.

We did not apply any locus-level filters, such as Hardy-Weinberg equilibrium, to our WGS results. We report per-locus WGS call rates for QCed (potentially related) individuals in each population. We used liftOver to lift the hg38 WGS calls to the hg19 reference genome (**URLs**). To compare the WGS calls to the imputed STR calls, we used CompareSTR from TRTools^73^ branch `compareSTR_upgrade` using the flags `--ignore-phasing --balanced-accuracy -- vcf2-beagle-probabilities`. We report multiple metrics at each locus, specifically concordance, the mean absolute summed-length difference, r^2^ and dosage r^2^.

For the following definitions, let *X* be the set of all samples, *A* be the set of all possible STR length alleles at a locus, let *S* = {*a*_1_ + *a*_2_|*a*_1_, *a*_2_ ∈ *A*} be the set of all summed-lengths possible at a locus (including the case of homozygous individuals when *a*_1_ = *a*_2_), for *x* ∈ *X* let *s_x,WGS_* be the summed-length call for sample *x* from WGS data, and for *x* ∈ *X*, *s* ∈ *S* let *Pr_x,imp_*(*s*) be the probability that sample *x* has a summed imputation length of *s* as output by the Beagle AP1 and AP2 FORMAT fields in the imputed VCF file.

We report (summed-length) per-locus concordances as *E_x∈X_*[*Pr_x,imp_*(*s_x,WGS_*)]. This metric has the advantage of being intuitive but is biased upwards for loci with a single very common allele and so should be interpreted cautiously for such loci. We also report mean absolute summed length differences as *E_x∈X_*[Σ*_s∈S_ Pr_x,imp_*(*s*) · |*s_x,WGS_* − *s*|]. This metric has similar caveats as the concordance metric. However, for highly multi-allelic loci where concordance is low, this metric can help quantify how close (or not) imputed calls are to the actual genotypes. We calculated r^2^ as the square of the weighted Pearson correlation between *s_x,WGS_* and *s* for each sample *x* ∈ *X* and all possible summed-lengths *s* ∈ *S* (so that there are |*X*| · |*S*| total values being correlated), weighting by the imputation probabilities *Pr_x,imp_*(*s*). This correlation measure is more comparable across loci with different numbers of alleles than concordance. It has the downside of being less intuitive and of being more sensitive to the WGS-vs-imputation concordance of rare long and short alleles than the WGS-vs-imputation concordance of common average-length alleles. We report dosage r^2^ as the square of the Pearson correlation between *s_x,WGS_* and the dosage Σ*_s∈S_ s* · *Pr*_*x,imp*_(*s*) for each sample *x* ∈ *X*. Dosage r^2^ is strictly greater than or equal to the weighted r^2^ measure. While the weighted r^2^ measure more directly measures the concordance of individual imputation probabilities with the WGS calls, the dosage r^2^ measure better estimates how analyses like GWAS, which condense imputed probabilities into dosages, will perform.

Lastly, at each locus we report the frequency of each summed-length according to WGS calls, and for all samples with each WGS summed-length we report the probability that imputation concurs with that length: *E*_*X*|*s*_*x,WGS*_ =*s*_[*Pr*_*x,imp*_(*s*_*x,WGS*_)].

### Replication in other populations

We separated the participants not in the White British group into population groups using the self-reported ethnicities summarized by UKB showcase data field 21000 (**URLs**). This field uses UKB showcase data coding 1001. We defined the following five populations based on those codings (counts give the maximal number of unrelated QC’ed participants, ignoring per-phenotype missingness):

● Black (African and Caribbean, n=7,562, codings 4, 4001, 4002, 4003)
● South Asian (Indian, Pakistani and Bangladeshi, n=7,397, codings 3001, 3002, 3003)
● Chinese (n=1,525, coding 5)
● Irish (n=11,978, coding 1002)
● Other White (White non-Irish non-British, n=15,838, coding 1003)

Self-reported ethnicities were collected from participants at three visits (initial assessment, repeat assessment, first imaging). The above groups also exclude participants who self-reported ethnicity at more than one visit and where their answers corresponded to more than one population (after ignoring ‘prefer not to answer’ `code=-3` responses). We did not include any participants who were neither in the White British population nor any of the above populations. Unlike for the determination of White British participants, genetic principal components were not used as filters for these categories.

For the association tests in these populations we applied the same procedures for sample quality control, unrelatedness filtering, phenotype transformations, and preparing genotypes and covariates as in the White British group. The only changes in procedure were that (a) we removed categorical covariate values where there were fewer than 50 participants with that value, (in which case we also removed those participants from analysis, as that would be too few to properly control for batch effects), whereas for White British individuals we used a cutoff of 0.1% instead and (b) we also applied this cutoff to the visit of measurement categorical covariate, resulting in some association tests that excluded individuals whose first measurement of the phenotype occurred outside the initial assessment visit. See **Supplementary Table 9** for details.

STRs were marked as replicating in another population (**Fig. 2**) if any of the traits confidently fine-mapped to that STR share the same direction of effect as the White British association and reached association p-value < 0.05 after multiple hypothesis correction (i.e. if there are three confidently fine-mapped traits, then an STR is marked as replicating in the Black population if any of them has association p-value < 0.05/3 = 0.0167 in the Black population).

We validated imputation STR lengths using WGS data in these populations as was done in the White British population, and report these results in **Supplementary Tables 4** and **5**. The number of samples in our QC’ed set that had WGS data were 2,990 Black, 3,373 South Asian, 619 Chinese, 5,174 Irish and 6,428 Other White samples, all roughly 40% of their respective populations.

### Logistic regression analysis of replication direction

We used logistic regression to quantitatively assess the impact of fine-mapping on replication rates while controlling for discovery p-value. For this analysis, to have sufficient sample sizes, we defined that an STR-trait association replicates in another population if it had the same direction of effect in that population as in the White British population, regardless of the replication p-value.

For each of the five replication populations, we compared four categories: all gwsig (genome-wide significant associations in the discovery population, i.e. p-value < 5e-8), FINEMAP (discovery p-value < 5e-8 and FINEMAP CP ≥ 0.8), SuSiE (discovery p-value < 5e-8 and SuSiE CP ≥ 0.8) and confidently fine-mapped STR (STR associations in our confidently fine-mapped set).

For each comparison, we used the function `statsmodels.formula.api.logit` from statsmodels v0.13.2 (**URLs**) to fit the logistic regression model:

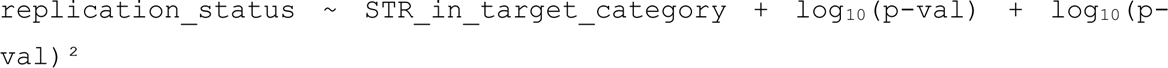

where `replication_status` is a binary variable indicating whether or not the given STR-trait association replicated in the other population, `p-val` is the discovery p-value, and `STR_in_target_category` is a binary variable indicating if the STR is in the target category.

For each replication population, we considered various models:

- All gwsig STRs with either FINEMAP, SuSiE, or confidently fine-mapped STRs as the target category.
- All FINEMAP STRs with confidently fine-mapped STRs as the target category.
- All SuSiE STRs with confidently fine-mapped STRs as the target category.

For each model, we performed a one-sided t-test for the hypothesis that the coefficient for the covariate `STR_in_target_category` was greater than zero, i.e. testing that being in the target category increased the predicted chance of replicating in the chosen population (**Supplementary Table 10**).

### Gene and transcription factor binding annotations

For all analyses not using GTEx data, gene annotations were based on GENCODE 38^75^ (**URLs**). Transcription factor binding sites and DNaseI hypersensitivity regions were identified by ENCODE^76^ overlapping several loci (*TAOK1*, *RHOT1* and *NCK2*) through visual inspection of the “Txn Factor ChIP” and “DNase Clusters” tracks in the UCSC Genome Browser^77^ and using the “Load from ENCODE” feature of the Integrative Genomics Viewer^78^.

### Enrichment testing

We tested the following categories for enrichment in STRs identified by our association testing pipeline:

- <u>Genomic feature</u>: We grouped records by feature type and restricted to features with support level 1 or 2 except for genes which don’t have a support level. We used bedtools^79^ to compute which features intersect each STR and the distance between each STR and the nearest feature of each feature type.
- <u>Repeat unit</u>: unit length and standardized repeat unit were defined as described above. Repeat units occurring in <1000 STRs were grouped by repeat length. Repeats whose unit could not be determined were considered as a separate category.
- <u>Overlap with expression STRs (eSTR)</u>: we tested for overlap with either all eSTRs or fine-mapped eSTRs as defined in our previous study to identify STR-gene expression associations in the Genotype Tissue Expression (GTEx) cohort^13^.

Enrichment p-values were computed using a Chi-squared test (without Yate’s continuity correction) if all cells had counts ≥ 5. A two-sided Fisher’s exact test was used otherwise. Chi-squared and Fisher’s exact tests were implemented using the `chi2_contingency` and `fisher_exact` functions from the Python scipy.stats package v1.7.3 (**URLs**).

### Expression association analysis in GTEx

We had previously analyzed associations^13^ between STRs and gene expression in GTEx V7. Here we reanalyzed those associations using GTEx V8. We obtained 30x Illumina whole genome sequencing (WGS) data from 652 unrelated participants in the Genotype-Tissue Expression project (GTEx)^37^ through dbGaP accession number phs000424.v8.p2. WGS data was accessed using fusera (**URLs**) through Amazon Web Services. We genotyped STRs using HipSTR^6^ v0.5 with HipSTR’s hg38 reference STR set (**URLs**). All individuals were genotyped jointly using default parameters. GTEx’s whole genome sequencing procedure is not PCR-free, which likely contributed to low call rates at long poly-A and GC-rich STRs. The resulting VCFs were filtered using DumpSTR from TRTools^73^, using the parameters --filter-hrun --hipstr-min-call-Q 0.9 --hipstr-min-call-DP 10 --hipstr-max-call-DP 1000 --hipstr-max-call-flank-indel 0.15 --hipstr-max-call-stutter 0.15 --min-locus-callrate 0.8 --min-locus-hwep 0.00001. We also removed STRs overlapping segmental duplication regions (UCSC Genome Browser^80^ h38.genomicSuperDups table). Altogether, 728,090 STRs remained for downstream analysis.

The *TAOK1* STR locus was filtered from this genotyping for having an 11% call rate, so we imputed the genotypes at that locus into the GTEx cohort. GTEx V7 SNP files were downloaded from GTEx data portal (**URLs**). SNPs on chromosome 17 were extracted and filtered to remove using vcftools with the parameters `--maf 0.01 --mac 3 --we 0.00001 --max-missing 0.8 --minQ 30`. We used Beagle v5.2 (beagle.28Jun21.220.jar) with the tool’s provided human genetic maps to impute STRs into the GTEx SNPs using the same reference panel used for imputation in the UKB cohort above^30^. From this imputation we took the best-guess genotypes of the *TAOK1* STR. We lifted the coordinates of the *TAOK1* STR from hg19 to hg38 using liftOver.

For each tissue, we obtained gene-level and transcript-level transcripts-per-million (TPM) values, exon-exon junction read counts, and exon read counts for each participant from GTEx Analysis V8 publicly available from the GTEx project website (**URLs**). Gene annotations are based on GENCODE v26^75^. We focused on 41 tissues with expression data for at least 100 samples (**Supplementary Table 13**). We restricted our analysis to protein-coding genes, transcripts and exons that did not overlap segmental duplication regions.

To control for population structure, we obtained publicly available genotype data on 2,504 unrelated individuals from the 1000 Genomes project^23^ genotyped with Omni 2.5 SNP genotyping arrays. We performed the following principal components analysis jointly on that data and the SNP genotypes based on WGS of the 652 individuals above. We removed all indels, multi-allelic SNPs, and SNPs with minor allele frequency less than 5%. We then used plink v.1.90b3.44 to subset these remaining SNPs to a set of SNPs in approximate linkage equilibrium with the command `--indep 50 5 2`. We excluded any remaining SNPs with missingness rate 5% or greater. We lastly ran principal component analysis using smartpca70^81^ v.13050 with default parameters.

We removed genes with TPM less than 1 in more than 90 percent of individuals. PEER factors^82^ were calculated using PEER v1.0 from the TPM values which remained after filtering. For each gene, we tested for association with each STR within 100kb. For each test we performed a linear regression between the STR’s dosage (sum of allele lengths) and gene expression (TPM). We included the loadings of the top five genotype principal components as computed above and the top N/10 PEER factors as covariates. The number of PEER factors was chosen to maximize the number of significant associations across a range of tissues. We did not include sex or age as covariates.

For each STR we computed Bonferroni-adjusted p-values to control for the number of gene **×** tissue tests performed for that STR. Associations that remained with adjusted p < 0.05 are shown in **Supplementary Table 12**.

We additionally used the GTEx cohort to test for an association between length of the bilirubin-associated dinucleotide repeat identified in *SLC2A2* with splicing efficiency in liver. We obtained exon-exon junction read counts and exon read counts from the GTEx website (**URLs**). We calculated the percent spliced in value for each exon in the manner suggested by Schafer et al.^83^. We performed a linear regression to test between the STR’s dosage and the percent spliced in of each exon within 10kb, using the top 5 ancestry principal components as covariates.

### Methylation association analysis in GTEx

This analysis used the same STR data and genotype principal components as the GTEx expression association analysis above.

We downloaded genome-wide DNA methylation (DNAm) profiling results from the NCBI GEO database under accession number GSE213478. This contained DNA methylation levels from the whole blood of 47 individuals who had been genotyped, including 754,054 autosomal CpG loci which passed quality control checks in that dataset (**URLs**)^84^. We lifted those loci from hg19 to hg38. We performed per-locus inverse-normalization of the DNAm data prior to downstream analysis. We calculated 5 PEER factors from the normalized DNAm data across quality-controlled loci from all chromosomes (including sex chromosomes) using PEER v1.0^82^, choosing 5 factors to match the number of PEER factors used by the methylation study which generated this data^84^.

We tested for associations between the methylation of each autosomal CpG locus and the length of each STR located within 100kb of that locus. For each such pair, we performed a linear regression between the STR’s dosage (sum of allele lengths across both chromosomes) and the inverse-normalized DNAm levels of that CpG locus, including the top five genotype principal components and the 5 PEER factors as covariates. We compared the effect sizes of these associations with those from another paper studying STR-methylation correlations in two separate cohorts in whole blood^17^ and found that they were broadly consistent (r=0.73, p<10^-200^, **Supplementary Fig. 18c**).

For each STR we computed Bonferroni-adjusted p-values to control for the number of CpG tests performed for that STR. Associations that remained with adjusted p < 0.05 are shown in **Supplementary Table 14**.

### Expression analysis of the CBL and RHOT1 STRs in Geuvadis

We applied HipSTR^6^ v0.6.2 to genotype STRs from HipSTR’s hg38 reference STR set (**URLs**) in 2,504 individuals from the 1000 Genomes Project^85^ for which high-coverage WGS data was available. Gene-level reads per kilobase per million reads (RPKM) values based on RNA-seq in lymphoblastoid cell lines for 462 1000 Genomes participants were downloaded from the Geuvadis website (**URLs**). Of these, 449 individuals were genotyped by HipSTR.

Similar to the GTEx analysis, we performed a linear regression between STR dosage (sum of allele lengths) and RPKM, adjusting for the top 5 genotype principal components (computed as above for the GTEx analysis, but only on populations included in Geuvadis and separately for Europeans and Africans) and N/10 (45) PEER factors as covariates. PEER analysis was applied using PEER v1.0 to the matrix of RPKM values after removing genes overlapping segmental duplications and those with RPKM less than 1 in more than 90% of LCL samples. We performed a separate regression analysis for African individuals (YRI) and European individuals (CEU, TSI, FIN, and GBR). After restricting to individuals with non-missing expression data and STR genotypes and who were not filtered as PCA outliers by smartpca^81,86^ included in EIGENSOFT v6.1.4, 447 LCL samples remained for analysis in each case (num. EUR=358, and AFR=89 for *CBL*, EUR=359 and AFR=88 for *RHOT1*).

### URLs

- 1000 Genomes individuals: https://www.internationalgenome.org/data-portal/sample using the “Download the list” tab
- associaTR: https://trtools.readthedocs.io/
- Beagle Human genetic maps: https://bochet.gcc.biostat.washington.edu/beagle/genetic_maps/
- fusera: https://github.com/ncbi/fusera
- GENCODE 38 (hg19): http://ftp.ebi.ac.uk/pub/databases/gencode/Gencode_human/release_38/GRCh37_mapping/gencode.v38lift37.annotation.gff3.gz
- Geuvadis: https://www.ebi.ac.uk/arrayexpress/experiments/E-GEUV-1/files/analysis_results/?ref=E-GEUV-1
- GTEx v8:

- https://www.gtexportal.org/home/datasets
- https://storage.googleapis.com/gtex_analysis_v8/rna_seq_data/GTEx_Analysis_2017-06-05_v8_RNASeQCv1.1.9_gene_tpm.gct.gz
- https://storage.googleapis.com/gtex_analysis_v8/rna_seq_data/GTEx_Analysis_2017-06-05_v8_STARv2.5.3a_junctions.gct.gz
- https://storage.googleapis.com/gtex_analysis_v8/rna_seq_data/GTEx_Analysis_2017-06-05_v8_RNASeQCv1.1.9_exon_reads.parquet
- h5py https://github.com/h5py/h5py
- HDF5: https://www.hdfgroup.org/HDF5/
- HipSTR STR reference https://github.com/HipSTR-Tool/HipSTR-references/raw/master/human/hg38.hipstr_reference.bed.gz
- liftOver:

- https://genome.ucsc.edu/cgi-bin/hgLiftOver, accessed on 2023/03/09
- https://hgdownload.soe.ucsc.edu/goldenPath/hg19/liftOver/hg19ToHg38.over.chain.gz
- ftp://hgdownload.soe.ucsc.edu/goldenPath/hg38/liftOver/hg38ToHg19.over.chain.gz
- Methylation in GTEx:

- Overview: https://www.ncbi.nlm.nih.gov/geo/query/acc.cgi?acc=GSE213478
- CpG locations: https://www.ncbi.nlm.nih.gov/geo/download/?acc=GSE213478&format=file
- NCBI protein database: https://www.ncbi.nlm.nih.gov/protein
- Pan-UKB:

- Overview: https://pan.ukbb.broadinstitute.org/downloads
- manifest: https://docs.google.com/spreadsheets/d/1AeeADtT0U1AukliiNyiVzVRdLYPkTbruQSk38DeutU8
- bilirubin SNP summary statistics: https://pan-ukb-us-east-1.s3.amazonaws.com/sumstats_flat_files/biomarkers-30840-both_sexes-irnt.tsv.bgz and https://pan-ukb-us-east-1.s3.amazonaws.com/sumstats_flat_files_tabix/biomarkers-30840-both_sexes-irnt.tsv.bgz.tbi
- Plink association testing best practices: https://www.cog-genomics.org/plink/2.0/assoc#glm
- PyMOL: https://pymol.org/2/
- rhdf5: https://www.bioconductor.org/packages/release/bioc/html/rhdf5.html
- Scipy.stats: https://docs.scipy.org/doc/scipy/reference/stats.html
- SNP-STR reference panel: https://gymreklab.com/2018/03/05/snpstr_imputation.html
- Statsmodels: https://www.statsmodels.org/stable/index.html
- UKB Data Showcase Search Page: https://biobank.ctsu.ox.ac.uk/crystal/search.cgi

## Supporting information

Margoliash-etal-SuppTables_biorxiv

Margoliash-etal-SuppText_biorxiv

## Acknowledgments

Research reported in this publication was supported in part by NIH/NHGRI grants R01HG010885 (M.G. and A.G.) and 1RM1HG011558 (M.G.). This research has been conducted using the UK Biobank Resource under application number 46122. We thank R. Wachs for helping with illustrations. We also thank K. Frazer and M. D’Antonio for helpful discussions and comments.

## GTEx Project

The Genotype-Tissue Expression (GTEx) Project was supported by the Common Fund of the Office of the Director of the National Institutes of Health (commonfund.nih.gov/GTEx). Additional funds were provided by the NCI, NHGRI, NHLBI, NIDA, NIMH, and NINDS. Donors were enrolled at Biospecimen Source Sites funded by NCI\Leidos Biomedical Research, Inc. subcontracts to the National Disease Research Interchange (10XS170), GTEx Project March 5, 2014 version Page 5 of 8 Roswell Park Cancer Institute (10XS171), and Science Care, Inc. (X10S172). The Laboratory, Data Analysis, and Coordinating Center (LDACC) was funded through a contract (HHSN268201000029C) to the The Broad Institute, Inc. Biorepository operations were funded through a Leidos Biomedical Research, Inc. subcontract to Van Andel Research Institute (10ST1035). Additional data repository and project management were provided by Leidos Biomedical Research, Inc. (HHSN261200800001E). The Brain Bank was supported supplements to University of Miami grant DA006227. Statistical Methods development grants were made to the University of Geneva (MH090941 & MH101814), the University of Chicago (MH090951,MH090937, MH101825, & MH101820), the University of North Carolina - Chapel Hill (MH090936), North Carolina State University (MH101819),Harvard University (MH090948), Stanford University (MH101782), Washington University (MH101810), and to the University of Pennsylvania (MH101822). The datasets used for the analyses described in this manuscript were obtained from dbGaP at http://www.ncbi.nlm.nih.gov/gap through dbGaP accession numbers phs000424.v7.p2 and phs000424.v8.p2.

## Author contributions

J.M. led, designed and performed the analyses and wrote the manuscript. S.F. helped oversee physiological interpretation of individual signals. Y.L. performed expression and methylation analyses of the GTEx data. X.Z. performed analyses of protein-coding STRs with AlphaFold. A.M. assisted with analysis of the *APOB* locus. A.G. and M.G. conceived the study, supervised analyses, and wrote the manuscript. All authors read and approved this manuscript.

## Code availability

The associaTR tool has been published as part of the TRTools^73^ package and tool-suite (https://trtools.readthedocs.io/). Code for performing most of the analyses and generating most of the figures in this paper can be found at https://github.com/LiterallyUniqueLogin/ukbiobank_strs/

## Data Availability

Full results from our analyses can be found at the URL https://gymreklab.com/science/2023/09/08/Margoliash-et-al-paper.html We are in the process of uploading imputed STR calls to the UK Biobank.

