## Supplementary material for "Polymorphic short tandem repeats make widespread contributions to blood and serum traits": Margoliash-etal-SuppText_biorxiv

#### Supplementary Notes

##### *Supplementary Note 1: Summary of fine-mapping models*

We applied two different fine-mapping methods, SuSiE[1] v0.11.42 and FINEMAP[2] v1.4. FINEMAP assumes a priori that each variant has an equal chance of being causal and that each variant's chance of causality is independent from the causal status of the other variants. It then attempts to stochastically walk over all reasonably-probable choices of collections of causal variants and assigns each causal configuration a posterior probability based on the observed associations and that prior. It then calculates posterior inclusion probabilities (PIPs) by summing over the walked configurations. For downstream analyses we required a single measurement of causality for each variable from both fine-mappers, which we called those variables' causal probabilities (CPs). For FINEMAP, we took each variable's PIP to be its FINEMAP CP.

While FINEMAP models each region as a collection of causal variants, SuSiE models each region as a collection of causal signals (called effects in the SuSiE manuscript), enabling SuSiE to study variants' contributions to each signal separately. To fit this model, SuSiE alternates between updating its model of each signal, attempting with each update to improve how the collection of all signals fits the observed data. As SuSiE only allows for the possibility of one variant being considered causal in any signal, if two variants are both estimated to be causal, they are forced during model fitting into different signals from one another. SuSiE calculates a value, alpha, for each variant in each signal – the probability that variant causes that signal – and then calculates a single PIP for each variant which gives the probability that the variant is causal in at least one signal. For reasons we explain below, unlike for FINEMAP, we chose an alpha value (or zero) as the SuSiE CP for each variant, rather than a PIP.

SuSiE reports a purity value for each signal, and we used that value to discard signals which were not well fine-mapped. SuSiE constructs 90%-credible sets for each signal so that the estimated probability of the credible set containing a variant causal for that signal is at least 90% (other values, such as 95%-credible sets, could be constructed similarly). SuSiE defines the purity of the credible set for each signal to be the minimum absolute correlation between any pair of variants in the set. The SuSiE manuscript suggests discarding signals with purity less than 0.5, but also states that the threshold is arbitrary. Looking at the distribution of credible set purities across all of our trait-regions (**Supplementary Fig. 3**) we decided to discard credible sets with purity less than 0.8, reasoning that the upper mode of the distribution is well above that threshold and that a signal containing two variants with correlation less than 0.8 has not been acceptably resolved.

SuSiE's PIPs are calculated across all credible sets regardless of purity, while we wished to conservatively only consider variants which had passed this added layer of scrutiny. Additionally, we saw that the PIP metric is sensitive to values of `L` (the number of signals fit per locus) – for one extreme example, in a locus with 57 variants, SuSiE run with `L=50` assigned each variant a PIP ≥ 0.5, which is unrealistic. So instead of using SuSiE's PIPs, we took each variant's highest alpha score from among credible sets with purity at least 0.8 as its SuSiE CP (or zero if it was in no such credible sets). This choice was uniformly conservative; CPs defined this way must be less than SuSiE's PIPs. We also found it to be less sensitive to `L` – we examine this more thoroughly in **Supplementary Note 3** below, but in the above example we note that there was only one credible set containing less than 50 variants and it was not pure, so each variant in that region has a SuSiE CP of 0. We compared our SuSiE CP metric to SuSiE's PIP metric in **Supplementary Fig. 4** and saw that, among variants residing in pure credible sets, these two measures only strongly differed for variants whose contribution to any single pure signal was small. As our downstream analyses focused on variants with large alpha values in pure credible sets, this means that our use of alpha values instead of PIPs was not strongly impactful in analyzing those variants. The impact is that we conservatively restricted which variants we examined. Lastly, we note that for high purity thresholds such as the one we use, our metric should be very similar to calling SuSiE's `susie_get_pip` function with the flag `prune_by_cs=TRUE`, a method not examined in the SuSiE manuscript and one we did not encounter until after performing this work.

***Supplementary Note 2: Comparing results across fine-mapping methods***

To assess the reliability of our fine-mapping results, we measured how often the two fine-mapping methods agreed with one another, and how sensitive they were to model settings. First, we used SuSiE's credible sets as a proxy for the truly independent signals in our data. We observed that while SuSiE and FINEMAP were in agreement for most of the signals, their results were strongly discordant for a sizable number of signals (**Supplementary Fig. 8**). In particular, for 8.5% of 90%-credible sets returned by SuSiE (which by definition are assigned at least a 90% chance of containing a causal variant), the sum of FINEMAP's assigned CPs for all variants in each of those sets was less than 0.1, indicating that FINEMAP concluded those sets had a < 10% chance of containing a causal variant.

Second, we looked at the variant level and saw that for most variants, the CPs from FINEMAP and SuSiE were similar (**Supplementary Fig. 9**), with FINEMAP assigning slightly higher CPs overall (possibly due to our use of SuSiE alpha values per variant instead of the overall PIPs). However, we again saw that SuSiE and FINEMAP markedly disagree at a subset of loci. For instance, among all SNPs and indels which at least one fine-mapping method assigned a CP ≥ 0.95 and the other method was decisive about their causality (assigning either CP ≥ 0.95 or CP ≤ 0.05), 12.2% of those were assigned a CP ≥ 0.95 by one method and a CP ≤ 0.05 by the other. For STRs, the fine-mapping methods disagreed at nearly half of the loci (43.5%) that were assigned CP ≥ 0.95 by one method and decisively scored by the other, suggesting the CPs for STRs are even less reliable. This highlights the need for additional quality control before stating that variants assigned a high posterior probability by a single fine-mapper are likely to be causal. Without any prior on which fine-mapper to believe when the two disagreed, we focused only on the 167 trait-STR associations for which association p-values were well below the genome-wide significance threshold (p-value<1e-10) and both fine-mappers assigned high CPs (CPs≥0.8) (**Supplementary Fig. 10a;** the 167 associations can be extracted from **Supplementary Table 4**).

***Supplementary Note 3: Assessing robustness of fine-mapping results***

We further assessed how robust our fine-mapping results were to fine-mapping instability and differences in the fine-mapping conditions, data filtering thresholds and algorithm metaparameters used. For SuSiE, we modified the inputs (1) `scaled_prior_variance`, (2) `tol`, (3) `residual_variance`, and (4) `L`, and also (5) changed the input genotypes from dosage genotypes to best-guess genotypes and (6) changed the prior to favor SNPs and indels over STRs as causal variants. For FINEMAP, we modified the inputs (1) `--prior-std` and (2) `--prob-conv-sss-tol` and also (3) filtered input variants with total non-major allele dosage less than 100, (4) filtered variants with p-value ≤ 5e-4, (5) set the prior on the number of causal variants per region to 4, and (6) changed the prior to favor SNPs and indels over STRs as causal variants. Further, we tested the ability to reproduce each fine-mappers results under the same conditions, and found that while SuSiE produced identical results when run on identical initial conditions, FINEMAP did not, so (7) we ran FINEMAP a second time with the same initial conditions.

We tested a few of the SuSiE settings on a subset of mean platelet volume fine-mapping regions before broader testing. We were encouraged that these settings had minimal impact on the results in this initial run and so did not include these conditions in our downstream tests for identifying confidently fine-mapped STRs. These settings were:

- `scaled_prior_variance` – This is the initial value for the estimation of the prior variance of the causal effect sizes relative to the variance of the phenotype. We changed this from the default of 0.2 to 5e-4 which resulted in no change to observed CPs.

- `tol` – This determines what amount of change in the objective function between optimization rounds is small enough to cause SuSiE to terminate. We reduced this from the default of 1e-3 to 1e-4 and saw only miniscule changes in the results (**Supplementary Fig. 11a**).

- `residual_variance` – This is the initial value for the estimation of the residual variance of the phenotype after controlling for all effects at the locus. By default, the `residual_variance` is initialized to the full variance of the phenotype, which in our study was slightly less than 1 (rank-inverse normalization set it to 1, and then regressing covariates out of the phenotype before running SuSiE reduced it slightly). We ran SuSiE with alternate `residual_variance` values of 0.95 and 0.8 and saw small changes in the results (**Supplementary Fig. 11b,c**), while noting that a residual variance value of 0.8 would be unrealistic for the large majority of fine-mapping regions in traits we studied.

- `L` – This is the number of signals SuSiE fits in a region, or equivalently, the upper bound on the number of causal variants SuSiE attempts to find (**Supplementary Fig. 11d**). In our original fine-mapping runs, we ran SuSiE with `L=10`, and only increased `L` in a region if needed (first to 30, and then to 50 if still needed, **Methods**). Below, we compared SuSiE runs with `L=10` in every region to runs with `L=50`. The SuSiE manuscript[1] states that inflated `L` values should not adversely impact model fitting because extraneous signals contribute small probabilities dispersed over many variants, thus not strongly changing any single variant's prediction, and also the learned effect sizes of these extra signals are shrunk towards zero. We see in our comparison that this only induces a large change in CP for a small fraction of variants. Of those, almost all of them are variants with non-zero CP values under the `L=10` case and zero CP in the `L=50` case. Thus, if they have any effect, this indicates that in most cases inflated values of `L` should lead to more conservative fine-mapping results. While overestimating `L` does not seem to harm our CP estimates, we found that when using SuSiE's standard PIP metric, overestimates of `L` may indeed lead to poor performance in some cases (see the discussion in **Supplementary Note 1** above).

On the other hand, the fine-mapping conditions we document below did impact the end results. For each of these conditions, we ran fine-mapping on the trait-regions of the 167 STR-trait associations above, (in the same manner as the fine-mapping section of the main **Methods**), and present supplementary figures showing how the CPs of variants changed under those conditions (**Supplementary Figs. 12, 13a-f**). Due to our lack of confidence in signals that were not robust to these choices, we restricted our set of confidently fine-mapped STR associations to the 119 associations that had CP ≥ 0.8 under each of those conditions (**Supplementary Table 5**). While our focus here was to find confidently fine-mapped STRs, and while the set of trait-regions used for running these tests was chosen for that purpose, **Supplementary Figs 12, 13a-f** identify similar trends for SNPs and indels in those regions. Thus, we hypothesize that these comparisons are relevant for fine-mapping of all variant types.

*SuSiE with best-guess genotypes vs dosage genotypes*

We ran SuSiE with the best-guess genotypes from our imputation pipeline instead of the dosage genotypes from that pipeline (**Supplementary Fig. 12**). Discrepancies in best-guess vs. dosages reflect imputation uncertainty, and contrasting runs under those two conditions allowed us to discard loci where this uncertainty strongly impacted the results.

*FINEMAP under identical conditions*

We reran FINEMAP on the same data with the same conditions and compared the CPs of the two runs (**Supplementary Fig. 13a**).

*FINEMAP with alternative p-value thresholds*

By default, we chose to filter as few variants as possible from our fine-mapping runs while still controlling for computational costs, which meant filtering variants with p>5e-2 from our FINEMAP runs and variants with p>5e-4 from our SuSiE runs, as FINEMAP was less computationally intensive. To check if this difference impacted the fine-mappers' results we ran FINEMAP having filtered all variants with p>5e-4 and compared it to our default FINEMAP runs (**Supplementary Fig. 13b**).

*FINEMAP with alternative choice of non-major allele frequency threshold*

To test whether FINEMAP results were strongly influenced by rare variants, we excluded all variants with total non-major allele dosage < 100 (population frequency less than approximately

0.015%) on top of the filter excluding variants with p-value ≥ 0.05 (**Supplementary Fig. 13c**). (Note that variants with total non-major allele dosage < 20 were excluded from association testing and thus from all fine-mapping runs).

Inadvertantly, when running FINEMAP with the alternative non-major allele frequency threshold, we only applied the threshold to SNPs and indels whose reference allele was the major allele, thus failing to filter out such variants whose reference allele had total dosage < 100 in the tested population. This reduces the useful interpretation of the results from this particular run with FINEMAP. Additionally, as a minor note, in this run we inadvertantly included the few variants with association p-value exactly equal to 0.05, in addition to including all variants with p-value < 0.05 as normal.

*FINEMAP with alternative choice of effect size prior*

FINEMAP's default `--prior-std` value is 0.05 which gives causal variants a default effect size of 0.25% of phenotypic variance. We modified this to `--prior-std 0.0224` to reflect published expected effect sizes for GWAS variants of about 0.05%[3] (**Supplementary Fig. 13d**).

*FINEMAP with alternative prior on the number of causal variants per region*

We ran FINEMAP with the prior of four causal variants per trait-region instead of one (**Supplementary Fig. 13e**). We did this by adding a column prob to the FINEMAP input Z file which contained the value 4/n for each variant, where n was the number of variants in the trait-region, and by running FINEMAP with the `--prior-snps` flag.

*FINEMAP with alternative `--prob-conv-sss-tol` stopping threshold*

We ran FINEMAP with the flag `--prob-conv-sss-tol` 0.0001 (reduced from the default of 0.001) (**Supplementary Fig. 13f**). This reduced what amount of change in the objective function over the last 100 rounds of optimization would be considered small enough to cause FINEMAP to terminate.

*Summary*

To conclude, 48 (28.3%) of the 167 STR-trait associations failed to replicate in one of the above alternate fine-mapping runs. The dosages vs best-guess genotypes choice when running SuSiE was the most impactful of these alternate conditions, accounting for 29 of those 48 cases, 24 of which did not fail any other alternate conditions (**Supplementary Fig. 12**), suggesting that imputation uncertainty has a sizable effect on downstream inferences.

Surprisingly, 7 (4.1%) of the 167 STR associations failed to replicate in the FINEMAP run under identical conditions, indicating that FINEMAP is moderately unstable (**Supplementary Fig. 13a**). This is a concern as FINEMAP has no seed parameter to allow for study reproducibility. Overall, 24 associations failed to replicate in at least one of the 6 FINEMAP alternate runs, 19 of which did not additionally fail the SuSiE best-guess condition. At these rates, it is difficult to distinguish if this is driven by FINEMAP's underlying instability, or if any of these other conditions are impactful by themselves. We suspect that FINEMAP's instability only appears at specific loci due to some nature of their LD patterns and do not currently hypothesize that FINEMAP results at all loci are subject to such instability.

For STR associations which failed to replicate in any of the above conditions, meaning that in the alternate run they had CP < 0.8, the associations rarely failed to replicate because of slight decreases in CP. Rather, there was an average decrease of 0.64 CP across failed replications, suggesting that when fine-mappers are sensitive to modeling conditions (or their own instability), those conditions strongly impact the end results.

Encouragingly, we saw that these comparisons agreed more frequently in regions containing variants which both the default SuSiE and FINEMAP runs agreed had high CPs (both CPs ≥ 0.8, **Supplementary Figs. 12, 13**) than in regions where the default SuSiE and FINEMAP runs disagreed (data not shown). This suggests that concordance between different fine-mapping algorithms may be able to provide security against the instability in the results of any single algorithm. While we focused on fine-mapping results for STRs, which generally showed lower concordance across methods than SNPs, our results suggest similar robustness checks should be performed when fine-mapping SNPs and other variant types.

There were several fine-mapping conditions we tested that strongly impacted the resulting CPs but that we did not use as filters when selecting our causal STR candidates since they represent unrealistic parameter choices. We report their values in **Supplementary Table 4**. Those conditions were:

- We ran FINEMAP with the flag `--prior-std 0.005`, corresponding to an expected effect size 0.0025% (**Supplementary Fig. 13g**). We concluded that this was much lower than the effect sizes we were hoping to detect.

- Both SuSiE and FINEMAP have the default assumption that each variant is as likely to be causal as any other variant (regardless of allele frequency). We instead conservatively ran SuSiE and FINEMAP with the prior assumption that SNPs and indels were 4x more likely to be causal than STRs. For this, we set the prior probability of causality for each SNP or indel to 4/(4*n_SNPs_indels + n_STRs) and for each STR to 1/(4*n_SNPs_indels + n_STRs). For SuSiE we did this by setting the `prior_weights` input to an array containing those probabilities. For FINEMAP we did this by adding a column `prob` to the FINEMAP input Z file which contained those probabilities, and by running FINEMAP with the `--prior-snps` flag. As expected, this resulted in overall decreased STR CPs (**Supplementary Fig. 14**). While we did not filter our candidate STRs based on this setting, we were encouraged to see that a majority of the strongest hits replicated despite this conservative setting.

Finally, we note there are other parameters which were not tested here but that could be tested for robustness. This includes whether FINEMAP results are sensitive to imputation uncertainty or overestimating `--n-causal-snps`, whether SuSiE results are sensitive to a non-major allele frequency threshold, or to increasing p-value thresholds higher than is computationally necessary and testing if either method's results are sensitive to the size of the trait-regions being fine-mapped.

**Supplementary Note 4: Additional details for coding fine-mapped STRs**

<u>*Coding trinucleotide repeat in* APOB</u>: This repeat did not initially appear in our list of confidently fine-mapped STRs due to limitations in our process for filtering indels which are STR alleles (**Methods**). We did not filter an indel imputed by the UKB team that corresponded exactly to the short allele of the STR imputed from our reference panel, since the indel/short allele corresponds to the deletion of an imperfect repeat sequence (GCCAGCAGC for a CAG repeat), and we cautiously only performed automated filtering for indels without imperfections. The presence of this indel alongside the STR during fine-mapping caused SuSiE's (but not FINEMAP's) results to, in some cases, show low confidence as to which of the two variants were causal. Specifically, FINEMAP assigned a CP of 1 to the STR for both traits apolipoprotein B and LDL cholesterol under each FINEMAP run used for filtering down to the confidently fine-mapped set. For the original run for the apolipoprotein B trait and for the best-guess run for both traits, SuSiE created a credible set containing both the indel and the STR and assigned each a CP of less than 0.8, causing the association not to pass our filters for confidently fine-mapped STRs. However, if we sum the SuSiE CPs of both variants in those runs we get a CP of over 0.97 in each case, making the apolipoprotein B association pass our confidently fine-mapped thresholds. Thus, we added this association to our confidently fine-mapped set. We note that the original SuSiE run for the LDL trait assigned low CPs to both the STR and the indel. While that was the only fine-mapping of the eight runs used for filtering that did not assign the pair of variants a combined CP ≥ 0.8 for LDL, it precludes us from adding the LDL association to the confidently fine-mapped set. For both the apolipoprotein B and LDL cholesterol associations, we updated the CPs in **Supplementary Tables 4** and **5** to reflect the combined CPs for both variants.

While we manually resolved this issue for the *APOB* STR, similar issues are likely to have caused other STRs in our set not to fine-map appropriately. We expect the choice of which indel representations to filter and which to treat as distinct variants will be critical for proper analysis of many STR loci in the future.

We used AlphaFold[4] to investigate whether the two common repeat alleles (referred to in previous literature as SP24 and SP27[5]) at *APOB* might have an effect on protein structure. Although analysis of the impact of small mutations has not yet been validated using AlphaFold[6], it could help generate hypotheses regarding the impact of protein-coding repeats on protein function. We restricted analysis to the first 600 amino acids as analysis of the full-length APOB protein (~4500) was computationally prohibitive. The two alleles did not induce any notable changes in predicted structure. Previous work on this repeat[7] suggests this variant, which resides in the signal peptide of the protein, affects secretion efficiency. We hypothesize that this reported change in secretion efficiency, rather than large changes in protein structure, drives this particular signal.

*Coding trinucleotide repeat in* E2F4: We identified a protein-coding (poly-serine) repeat in *E2F4* confidently fine-mapped to multiple red blood cell traits. We similarly used AlphaFold[4] to assess the impact of varying the number of serine repeats on E2F4's protein structure, which did not result in any noticeable difference (**Supplementary Fig. 16a**). However E2F4 is known to form a complex with RBL2, in which RBL2 stabilizes E2F4 and protects it from degradation via the ubiquitin-proteasome pathway[8]. This prompted us to explore the joint structure of E2F4 with RBL2. In addition to not being validated for predicting the effect of mutations, the standard AlphaFold software is also not designed or validated for predicting the structure of protein complexes[6]. Nonetheless, it can be used for such prediction, and that may generate useful hypotheses. In this case, AlphaFold results suggest E2F4 and RBL2 are more tightly bound when E2F4 contains a short vs. long poly-serine track (**Supplementary Fig. 16b**). We therefore hypothesize that long poly-serine alleles destabilize the complex and lead to faster degradation of E2F4. While existing proteomics datasets[9,10] did not allow us to test this hypothesis, it does have the expected direction of association: previous knockout[11] and knockdown[12] experiments show that presence of E2F4 is positively associated with counts of other cell types, and our GWAS shows a negative association between this STR's length and red blood cell counts.

*Methods for 3D structure simulation and analysis using AlphaFold:* We performed protein 3D structure simulations using AlphaFold v2.2.0[4]. The simulations were conducted using 1 NVIDIA V100 SMX2 GPU. We used the settings `--max_template_date=2020-05-14 --use_gpu_relax=True --model_preset=monomer` . The simulated protein structures were aligned and visualized using PyMOL 2.5.4 (**URLs**).

The `NP_000375` fasta file for APOB was downloaded from the NCBI protein database (**URLs**). Due to the large size of the full APOB protein (4563 amino acids) and the focus on the N terminal STR variants, only the first 600 amino acids were used in the structure simulation.

The `NP_001941` fasta file for E2F4 was downloaded from the NCBI protein database. The full length of the E2F4 protein was used for the structure simulation. Among all the poly-serine variants, we selected the shortest (containing a 12 unit repeat) and longest variants (containing a 19 unit repeat, the reference) for comparison. Note that the poly-serine sequence in E2F4 is 2 amino acids longer than the repeat as the last two serines use different codings than the repeat unit AGC. (e.g. the reference is SSSSSSSSSSSSSSNSNSSSSS which is 21 amino acids, while the reference repeat is 19 units long).

For the E2F4_RBL2 complex, we downloaded the `NP_001310537` fasta file for RBL2 from the NCBI protein database. The E2F4 sequences were linked with the RBL2 sequence using a flexible 70 amino acid linker $(GGGGS)_{14}$.

The protein sequences used for the simulations are listed below:

##### APOB longer STR allele

```
MDPPRPALLALLALPALLLLLLAGARAEEEMLENVSLVCPKDATRFKHLRKYTYNYEAESSSGVPGTADSRSATRINCKVELEVPQLCSFILKTSQCTLKEVYGFNPEGKA
LLKKTKNSEEFAAAMSRYELKLAIPEGKQVFLYPEKDEPTYILNIKRGIISALLVPPETEEAKQVLFLDTVYGNCSTHFTVKTRKGNVATEISTERDLGQCDRFKPIRTGI
SPLALIKGMTRPLSTLISSSQSCQYTLDAKRKHVAEAICKEQHLFLPFSYKNKYGMVAQVTQTLKLEDTPKINSRFFGEGTKKMGLAFESTKSTSPPKQAEAVLKTLQELK
KLTISEQNIQRANLFNKLVTELRGLSDEAVTSLLPQLIEVSSPITLQALVQCGQPQCSTHILQWLKRVHANPLLIDVVTYLVALIPEPSAQQLREIFNMARDQRSRATLYA
LSHAVNNYHKTNPTGTQELLDIANYLMEQIQDDCTGDEDYTYLILRVIGNMGQTMEQLTPELKSSILKCVQSTKPSLMIQKAAIQALRKMEPKDKDQEVLLQTFLDDASPG
DKRLAAYLMLMRSPSQADINKIVQILPWEQNEQVKNFVASHIANI
```

##### APOB shorter STR allele

```
MDPPRPALLALPALLLLLLAGARAEEEMLENVSLVCPKDATRFKHLRKYTYNYEAESSSGVPGTADSRSATRINCKVELEVPQLCSFILKTSQCTLKEVYGFNPEGKALLK
KTKNSEEFAAAMSRYELKLAIPEGKQVFLYPEKDEPTYILNIKRGIISALLVPPETEEAKQVLFLDTVYGNCSTHFTVKTRKGNVATEISTERDLGQCDRFKPIRTGISPL
ALIKGMTRPLSTLISSSQSCQYTLDAKRKHVAEAICKEQHLFLPFSYKNKYGMVAQVTQTLKLEDTPKINSRFFGEGTKKMGLAFESTKSTSPPKQAEAVLKTLQELKKLT
ISEQNIQRANLFNKLVTELRGLSDEAVTSLLPQLIEVSSPITLQALVQCGQPQCSTHILQWLKRVHANPLLIDVVTYLVALIPEPSAQQLREIFNMARDQRSRATLYALSH
AVNNYHKTNPTGTQELLDIANYLMEQIQDDCTGDEDYTYLILRVIGNMGQTMEQLTPELKSSILKCVQSTKPSLMIQKAAIQALRKMEPKDKDQEVLLQTFLDDASPGDKR
LAAYLMLMRSPSQADINKIVQILPWEQNEQVKNFVASHIANILNS
```

##### E2F4 longer STR allele

```
MAEAGPQAPPPPGTPSRHEKSLGLLTTKFVSLLQEAKDGVLDLKLAADTLAVRQKRRIYDITNVLEGIGLIEKKSKNSIQWKGVGPGCNTREIADKLIELKAEIEELQQRE
QELDQHKVWVQQSIRNVTEDVQNSCLAYVTHEDICRCFAGDTLLAIRAPSGTSLEVPIPEGLNGQKKYQIHLKSVSGPIEVLLVNKEAWSSPPVAVPVPPPEDLLQSPSAV
STPPPLPKPALAQSQEASRPNSPQLTPTAVPGSAEVQGMAGPAAEITVSGGPGTDSKDSGELSSLPLGPTTLDTRPLQSSALLDSSSSSSSSSSSSSSNSNSSSSSGPNPST
SFEPIKADPTGVLELPKELSEIFDPTRECMSSELLEELMSSEVFAPLLRLSPPPGDHDYIYNLDESEGVCDLFDVPVLNL
```

##### E2F4 shorter STR allele

```
MAEAGPQAPPPPGTPSRHEKSLGLLTTKFVSLLQEAKDGVLDLKLAADTLAVRQKRRIYDITNVLEGIGLIEKKSKNSIQWKGVGPGCNTREIADKLIELKAEIEELQQRE
QELDQHKVWVQQSIRNVTEDVQNSCLAYVTHEDICRCFAGDTLLAIRAPSGTSLEVPIPEGLNGQKKYQIHLKSVSGPIEVLLVNKEAWSSPPVAVPVPPPEDLLQSPSAV
STPPPLPKPALAQSQEASRPNSPQLTPTAVPGSAEVQGMAGPAAEITVSGGPGTDSKDSGELSSLPLGPTTLDTRPLQSSALLDSSSSSSNSNSSSSSGPNPSTSFEPIKA
DPTGVLELPKELSEIFDPTRECMSSELLEELMSSEVFAPLLRLSPPPGDHDYIYNLDESEGVCDLFDVPVLNL
```

##### E2F4-RBL2 longer STR allele

```
MAEAGPQAPPPPGTPSRHEKSLGLLTTKFVSLLQEAKDGVLDLKLAADTLAVRQKRRIYDITNVLEGIGLIEKKSKNSIQWKGVGPGCNTREIADKLIELKAEIEELQQRE
QELDQHKVWVQQSIRNVTEDVQNSCLAYVTHEDICRCFAGDTLLAIRAPSGTSLEVPIPEGLNGQKKYQIHLKSVSGPIEVLLVNKEAWSSPPVAVPVPPPEDLLQSPSAV
STPPPLPKPALAQSQEASRPNSPQLTPTAVPGSAEVQGMAGPAAEITVSGGPGTDSKDSGELSSLPLGPTTLDTRPLQSSALLDSSSSSSSSSSSSSSNSNSSSSSGPNPST
SFEPIKADPTGVLELPKELSEIFDPTRECMSSELLEELMSSEVFAPLLRLSPPPGDHDYIYNLDESEGVCDLFDVPVLNLGGGGSGGGGSGGGGSGGGGSGGGGSGGGGSG
GGGSGGGGSGGGGSGGGGSGGGGSGGGGSGGGGSGGGGSMPSGGDQSPPPPPPPPAAAASDEEEEDDGEAEDAAPPAESPTPQIQQRFDELCSRLNMDEAARAEAWDSYRS
MSESYTLEGNDLHWLACALYVACRKSVPTVSKGTVEGNYVSLTRILKCSEQSLIEFFNKMKKWEDMANLPPHFRERTERLERNFTVSAVIFKKYEPIFQDIFKYPQEEQPR
QQRGRKQRRQPCTVSEIFHFCWVLFIYAKGNFPMISDDLVNSYHLLLCALDLVYGNALQCSNRKELVNPNFKGLSEDFHAKDSKPSSDPPCIIEKLCSLHDGLVLEAKGIK
EHFWKPYIRKLYEKKLLKGKEENLTGFLEPGNFGESFKAINKAYEEYVLSVGNLDERIFLGEDAEEEIGTLSRCLNAGSGTETAERVQMKNILQQHFDKSKALRISTPLTG
```

VRYIKENSPCVTPVSTATHSLSRLHTMLTGLRNAPSEKLEQILRTCSRDPTQAIANRLKEMFEIYSQHFQPDEDFSNCAKEIASKHFRFAEMLYYKVLESVIEQEQKRLGD
MDLSGILEQDAFHRSLLACCLEVVTFSYKPPGNFPFITEIFDVPLYHFYKVIEVFIRAEDGLCREVVKHLNQIEEQILDHLAWKPESPLWEKIRDNENRVPTCEEVMPPQN
LERADEICIAGSPLTPRRVTEVRADTGGLGRSITSPTTLYDRYSSPPASTTRRRLFVENDSPSDGGTPGRMPPQPLVNAVPVQNVSGETVSVTPVPGQTLVTMATATVTAN
NGQTVTIPVQGIANENGGITFFPVQVNVGGQAQAVTGSIQPLSAQALAGSLSSQQVTGTTLQVPGQVAIQQISPGGQQQKQGQSVTSSSNRPRKTSSLSLFFRKVYHLAAV
RLRDLCAKLDISDELRKKIWTCFEFSIIQCPELMMDRHLDQLLMCAIYVMAKVTKEDKSFQNIMRCYRTQPQARSQVYRSVLIKGKRKRRNSGSSDSRSHQNSPTELNKDR
TSRDSSPVMRSSSTLPVPQPSSAPPTPTRLTGANSDMEEEERGDLIQFYNNIYIKQIKTFAMKYSQANMDAPPLSPYPFVRTGSPRRIQLSQNHPVYISPHKNETMLSPRE
KIFYYFSNSPSKRLREINSMIRTGETPTKKRGILLEDGSESPAKRICPENHSALLRRLQDVANDRGSH

E2F4-RBL2 shorter STR allele

MAEAGPQAPPPPGTPSRHEKSLGLLTTKFVSLLQEAKDGVLDLKLAADTLAVRQKRRIYDITNVLEGIGLIEKKSKNSIQWKGVGPGCNTREIADKLIELKAEIEELQQRE
QELDQHKVWVQQSIRNVTEDVQNSCLAYVTHEDICRCFAGDTLLAIRAPSGTSLEVPIPEGLNGQKKYQIHLKSVSGPIEVLLVNKEAWSSPPVAVPVPPPEDLLQSPSAV
STPPPLPKPALAQSQEASRPNSPQLTPTAVPGSAEVQGMAGPAAEITVSGGPGTDSKDSGELSSLPLGPTTLDTRPLQSSALLDSSSSSSNSNSSSSSGPNPSTSFEPIKA
DPTGVLELPKELSEIFDPTRECMSSELLEELMSSEVFAPLLRLSPPPGDHDYIYNLDESEGVCDLFDVPVLNLGGGGSGGGGSGGGGSGGGGSGGGGSGGGGSGGGGSGGG
GSGGGGSGGGGSGGGGSGGGGSGGGGSGGGGSMPSGGDQSPPPPPPPPAAAASDEEEEDDGEAEDAAPPAESPTPQIQQRFDELCSRLNMDEAARAEAWDSYRSMSESYTL
EGNDLHWLACALYVACRKSVPTVSKGTVEGNYVSLTRILKCSEQSLIEFFNKMKKWEDMANLPPHFRERTERLERNFTVSAVIFKKYEPIFQDIFKYPQEEQPRQQRGRKQ
RRQPCTVSEIFHFCWVLFIYAKGNFPMISDDLVNSYHLLLCALDLVYGNALQCSNRKELVNPNFKGLSEDFHAKDSKPSSDPPCIIEKLCSLHDGLVLEAKGIKEHFWKPY
IRKLYEKKLLKGKEENLTGFLEPGNFGESFKAINKAYEEYVLSVGNLDERIFLGEDAEEEIGTLSRCLNAGSGTETAERVQMKNILQQHFDKSKALRISTPLTGVRYIKEN
SPCVTPVSTATHSLSRLHTMLTGLRNAPSEKLEQILRTCSRDPTQAIANRLKEMFEIYSQHFQPDEDFSNCAKEIASKHFRFAEMLYYKVLESVIEQEQKRLGDMDLSGIL
EQDAFHRSLLACCLEVVTFSYKPPGNFPFITEIFDVPLYHFYKVIEVFIRAEDGLCREVVKHLNQIEEQILDHLAWKPESPLWEKIRDNENRVPTCEEVMPPQNLERADEI
CIAGSPLTPRRVTEVRADTGGLGRSITSPTTLYDRYSSPPASTTRRRLFVENDSPSDGGTPGRMPPQPLVNAVPVQNVSGETVSVTPVPGQTLVTMATATVTANNGQTVTI
PVQGIANENGGITFFPVQVNVGGQAQAVTGSIQPLSAQALAGSLSSQQVTGTTLQVPGQVAIQQISPGGQQQKQGQSVTSSSNRPRKTSSLSLFFRKVYHLAAVRLRDLCA
KLDISDELRKKIWTCFEFSIIQCPELMMDRHLDQLLMCAIYVMAKVTKEDKSFQNIMRCYRTQPQARSQVYRSVLIKGKRKRRNSGSSDSRSHQNSPTELNKDRTSRDSSP
VMRSSSTLPVPQPSSAPPTPTRLTGANSDMEEEERGDLIQFYNNIYIKQIKTFAMKYSQANMDAPPLSPYPFVRTGSPRRIQLSQNHPVYISPHKNETMLSPREKIFYYFS
NSPSKRLREINSMIRTGETPTKKRGILLEDGSESPAKRICPENHSALLRRLQDVANDRGSH

#### Supplementary Note 5: Additional details for non-coding fine-mapped STRs

*Dinucleotide repeat in* SLC2A2 (GLUT2): We identified a dinucleotide repeat immediately upstream of exon 4 of *SLC2A2* as a confidently fine-mapped STR for bilirubin. While *SLC2A2* has not previously been causally linked to bilirubin levels, *SLC2A2* mediates glucose transport to hepatocytes, where glucose is stored in the form of glycogen[13]. Glycogen degradation produces intermediates that are substrates in the process that regulates bilirubin conjugation and excretion[14,15] and thus could potentially impact bilirubin levels in the blood. This effect of *SLC2A2* on bilirubin levels may be partially corroborated by a large cohort study on babies with congenital hyperinsulinemic hypoglycemia, a condition that inhibits glycogen breakdown, which reported elevated bilirubin in that population[16].

*Tetranucleotide repeat in* ESR2: We identified a GTTT repeat in an intron of *ESR2* whose length is negatively associated with haemoglobin concentration, red blood cell count, and haematocrit. *ESR2* is known to regulate red blood cell production. Studies conducted in populations chronically exposed to high altitude hypoxia, a driver of erythrocytosis (excess red blood cell production), demonstrated inhibition of erythrocytosis through activation of estrogen beta signaling in ex vivo models[17]. These observations are corroborated by a study of rat models under hypoxia, where beta-estrogen treatment reduced circulating levels of erythropoietin, a kidney-derived factor that stimulates red blood cell production[18].

We additionally identified a negative association between length of this STR and *ESR2* expression. However, the expected direction of association between *ESR2* expression and red blood cell count is unclear. Multiple *ESR2* isoforms exist, either as a result of alternative splicing of the last coding exons (exon 8 and exon 9, respectively), deletion of one or more coding exons, or alternative usage of untranslated exons in the 5′ region[19]. One of the five isoforms found in humans even has an undetectable affinity to estrogen, and instead was found to antagonize estrogen-alpha receptor signaling[20]. Thus any change in overall *ESR2* expression would need to be understood in the context of the isoforms whose expressions are changing and which tissues those isoforms are common in, complicating any mechanistic predictions.

#### Supplementary Figures

##### Supplementary Figure 1: Comparison of SNP alternate allele frequencies between our SNP-STR reference panel and UKB phased hard-called variants

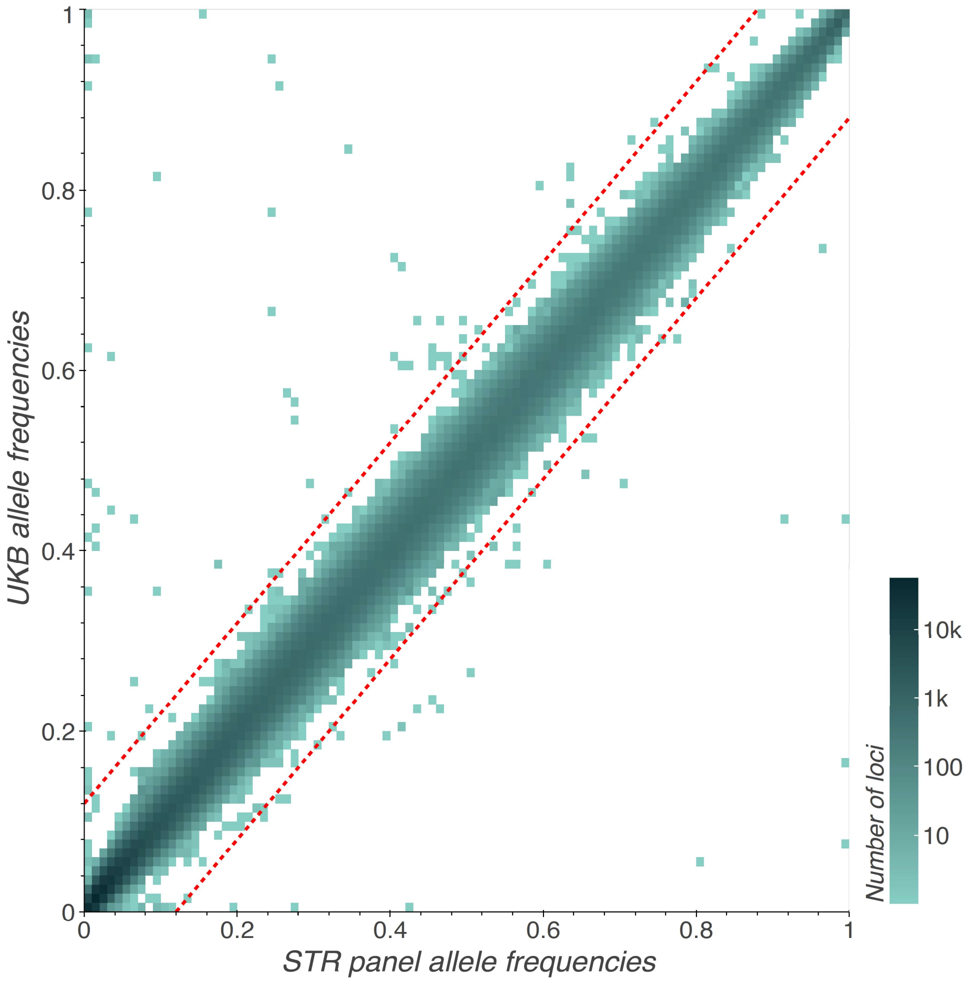


The x-axis indicates the alternate allele frequency of variants calculated from the European individuals in our SNP-STR reference panel[21] (**Main Text URLs**). The y-axis indicates their alternate allele frequency in unrelated participants in the White British population in the UKB. We filtered variants with more than a 12% difference in alternate allele frequency (indicated by the red diagonal lines). The color gradient represents the number of variants ($\log_{10}$ scale) whose p-values fall in each region. White regions contain no variants.

**Supplementary Figure 2: Comparison of association p-values between our pipeline and summary statistics published by Pan UKBB**.

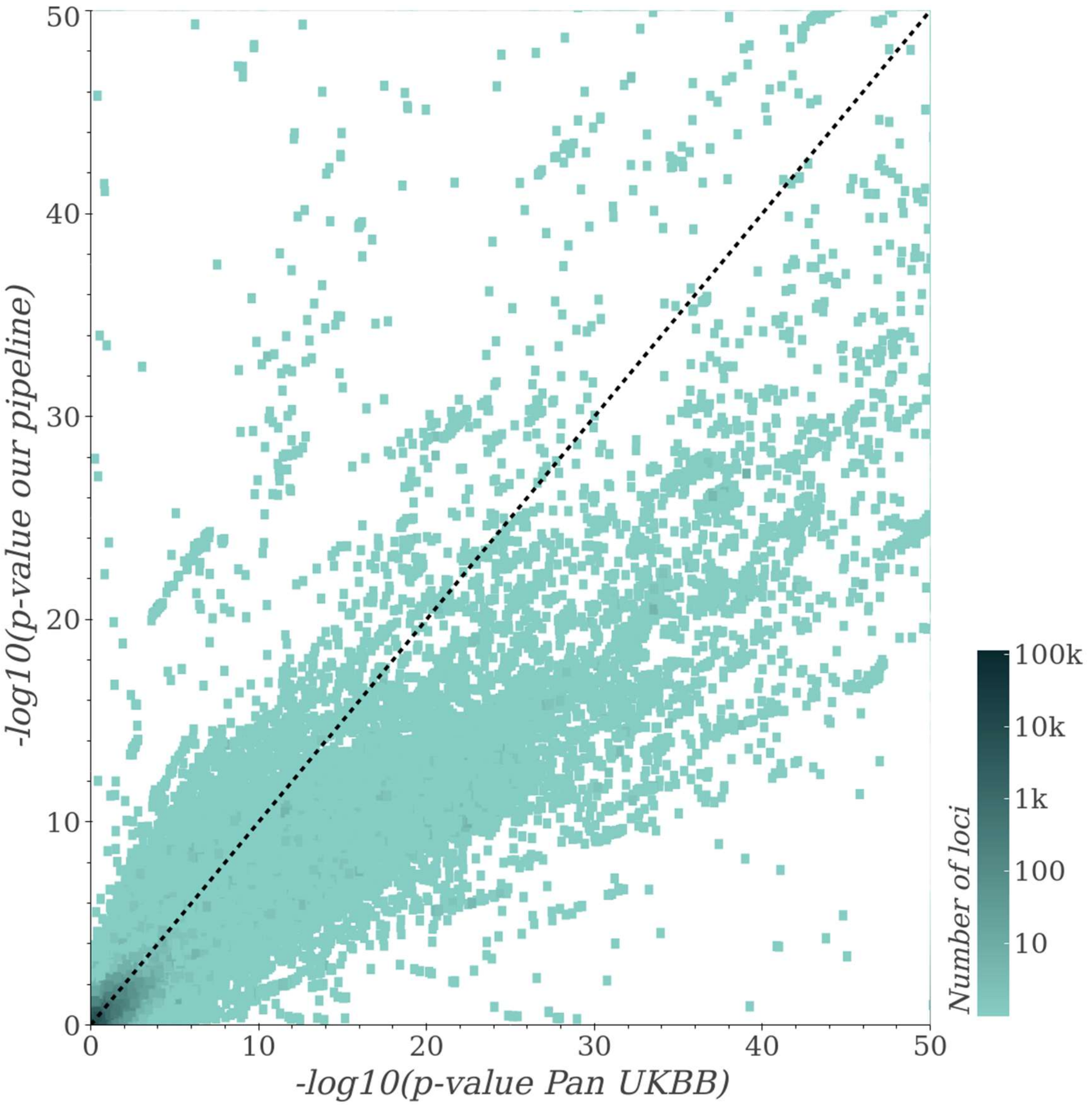


Heatmap of -$\log_{10}$ p-values obtained from the Pan UKBB[22] study of UKB data (x-axis) vs. from our study (y-axis) for total bilirubin associations with SNPs and indels. The color gradient represents the number of variants ($\log_{10}$ scale) whose p-values fall in each region. White regions contain no variants. P-values less than 1e-50 are truncated. Our pipeline's p-values are highly correlated with Pan UKBB's but are overall more conservative, which may be attributable to differences in models used (linear mixed model for Pan-UKB vs. linear model used here).

**Supplementary Figure 3: Distribution of SuSiE 90%-credible set purities**

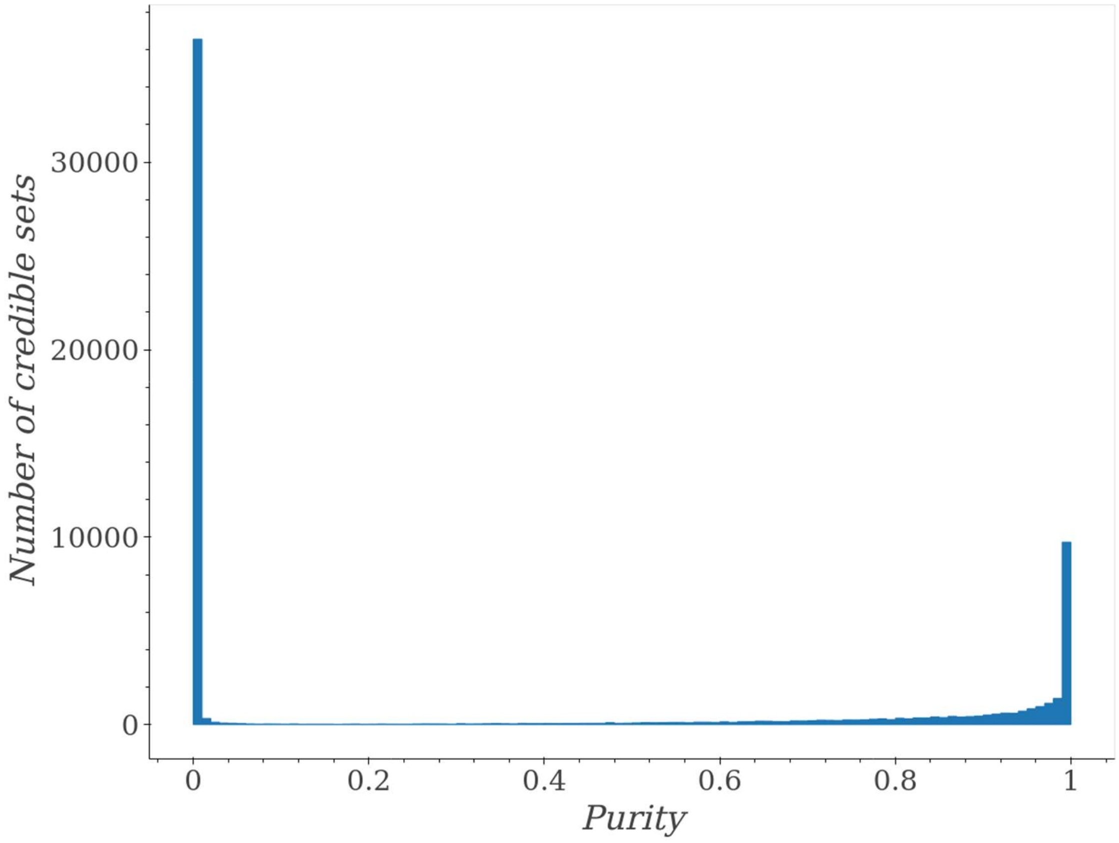


Distribution of SuSiE 90%-credible set purities across all trait-regions. (The rightmost bin is inclusive, containing SuSiE credible sets with purity up to and including 1, e.g. those that consist of a single variant.) Purity is defined as the minimum absolute correlation between any pair of variants in the set. For subsequent analyses, we discarded credible sets with purity < 0.8.

**Supplementary Figure 4: PIP vs alpha values assigned by SuSiE**

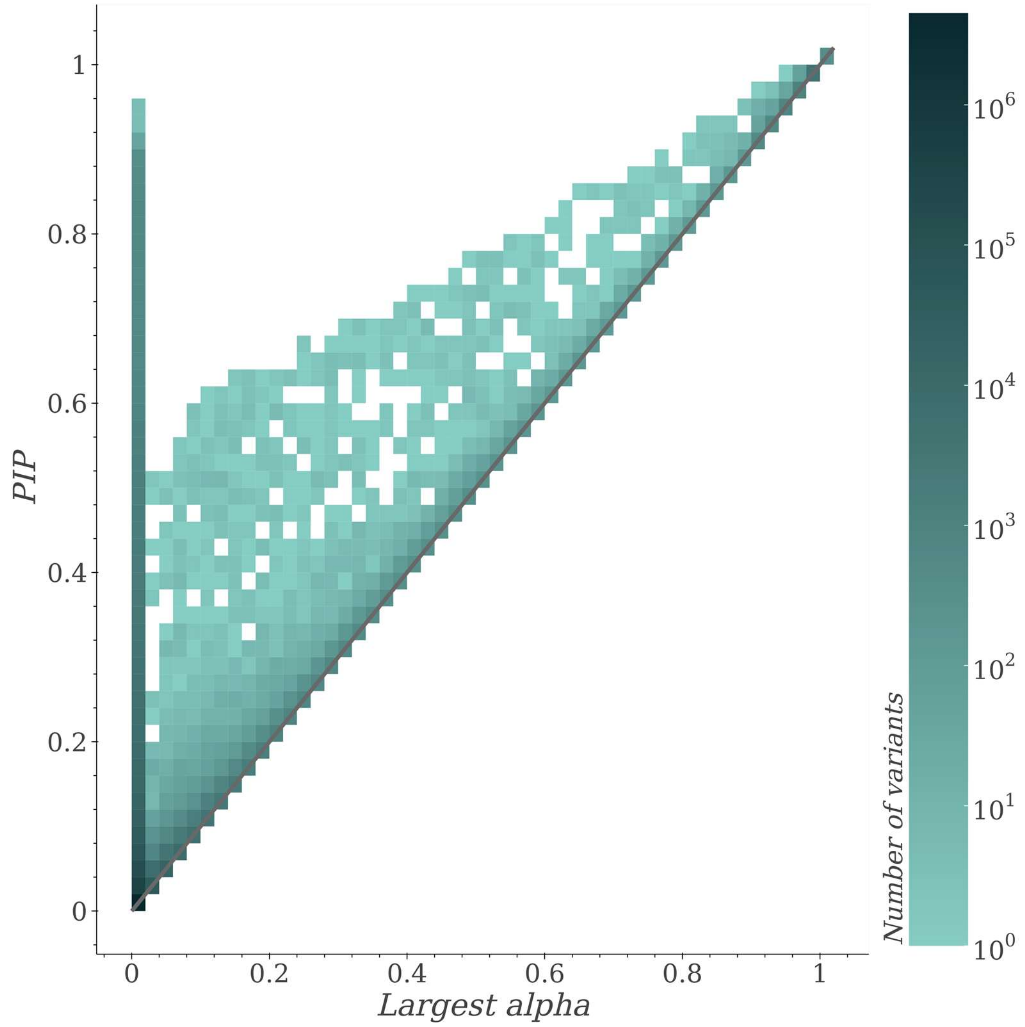


Largest alpha value across pure credible sets (x-axis) vs. PIP (y-axis) for all variants across all trait regions. Color ($\log_{10}$ scale) indicates the number of data points falling in each bin.

**Supplementary Figure 5: Contribution of variants to signals genome-wide by variant CP**

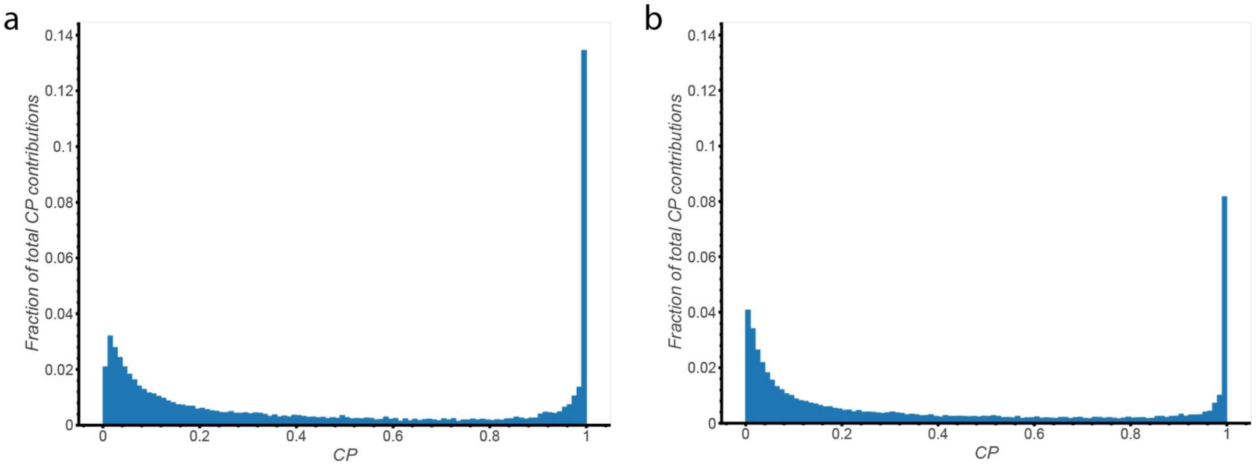


Summed contribution of genome-wide significant variants across all regions binned by variant CP as a fraction of the total CP of all genome-wide significant variants across all regions for **(a)** SuSiE and **(b)** FINEMAP. (The rightmost bin for each graph is inclusive, containing variants with CPs up to and including 1.) The total contribution of all variants across all regions with CP < 0.1 was 29.3% for SuSiE and 35.1% for FINEMAP.

**Supplementary Figure 6: Effect sizes for causal variants in simulation strategy 1**

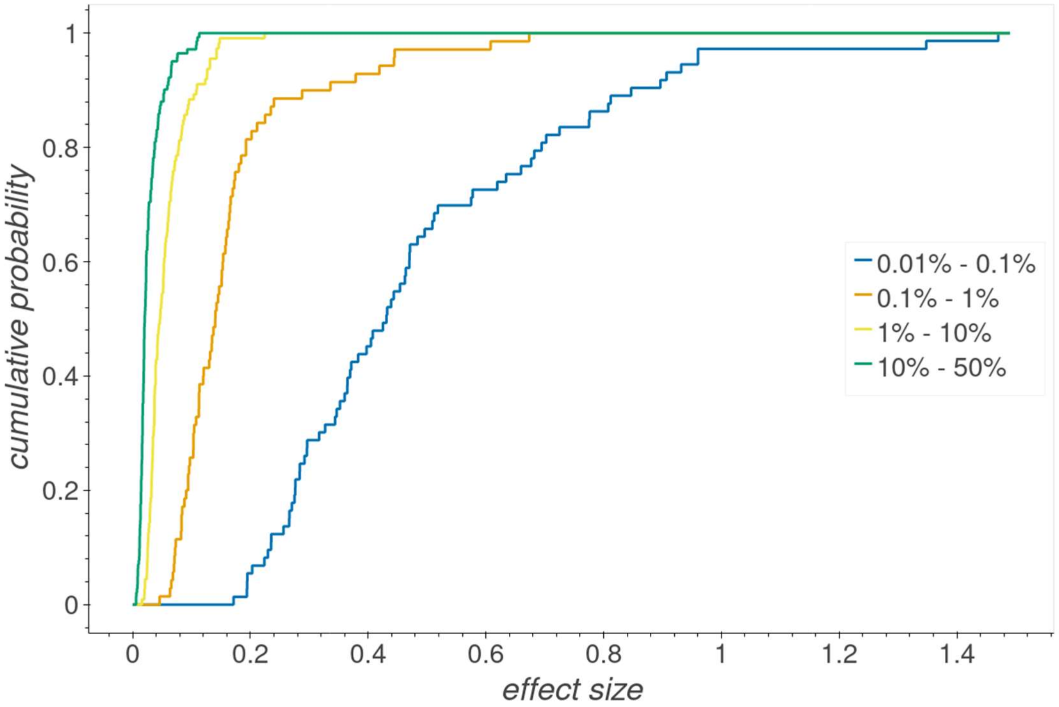


Cumulative distribution functions of the discrete effect size distributions for each minor allele frequency bin for simulation strategy 1. Per the **Methods**, these effect sizes were drawn from all SNPs/indels in all platelet count regions that had either FINEMAP CP ≥ 0.5 or SuSiE CP ≥ 0.5. SNPs/indels chosen to be causal for strategy 1 simulations had their effect sizes drawn from the bin their minor allele frequency corresponded to. As expected, more common variants tend to have smaller effect sizes than rarer variants.

**Supplementary Figure 7: Concordance between STR length imputation and WGS calls**

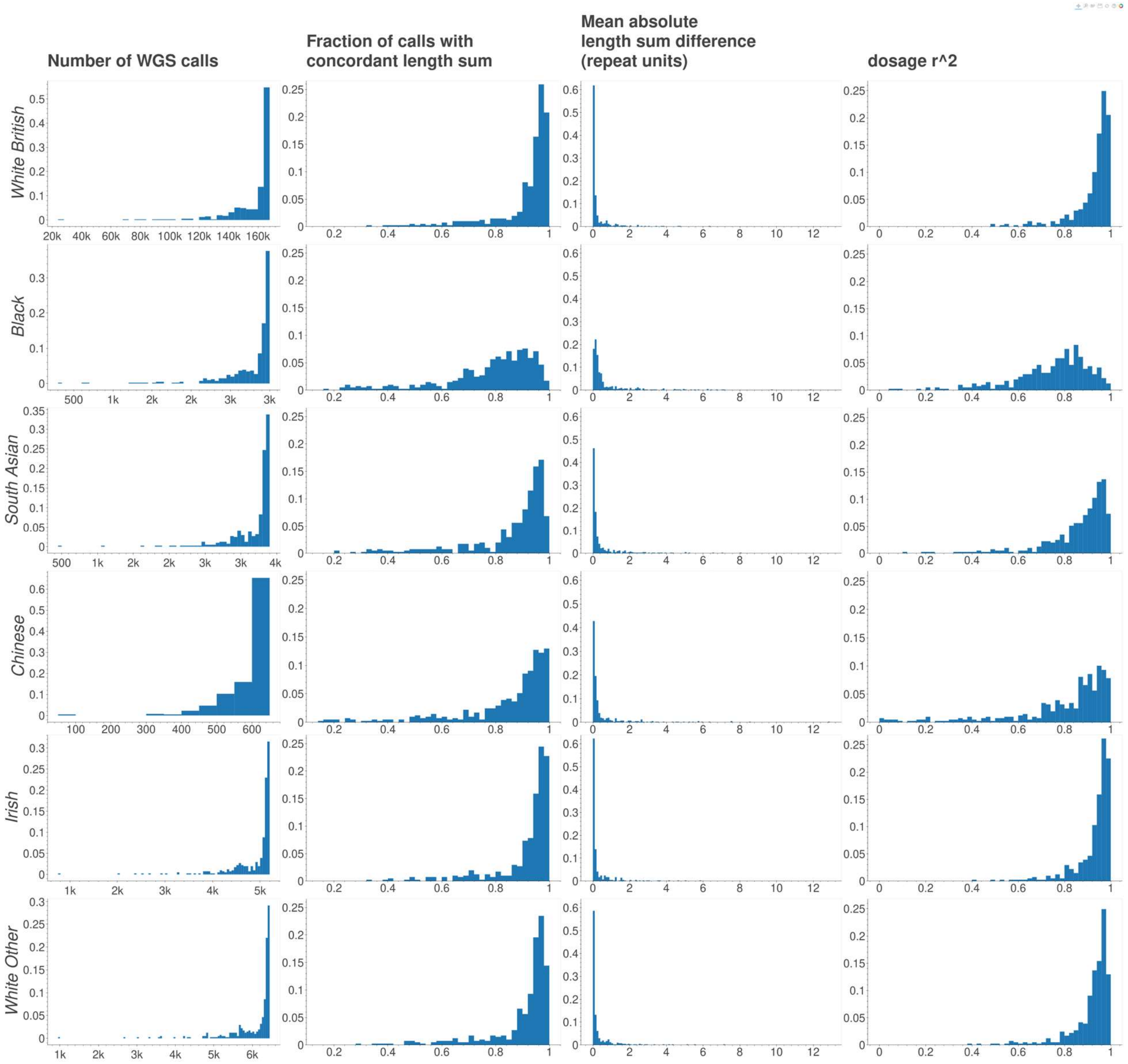


Each histogram is a distribution over the 409 distinct STRs with an association with either FINEMAP or SuSiE CP ≥ 0.8 (**Supplementary Table 4**). Each row represents a different ethnicity group of QCed (potentially related) individuals. The first column displays the number of WGS calls per locus for each group. Each subsequent column is a different measurement of per-locus concordance between STR length calls from imputation and WGS: the fraction of concordant length sums, the mean absolute difference between length sums and the correlation between length sum dosages (**Methods**). For sake of comparison, all histograms in the same column share the same x-axis and the same y-axis bounds, excepting the first column.

**Supplementary Figure 8: Total CPs assigned to SuSiE credible sets by FINEMAP**

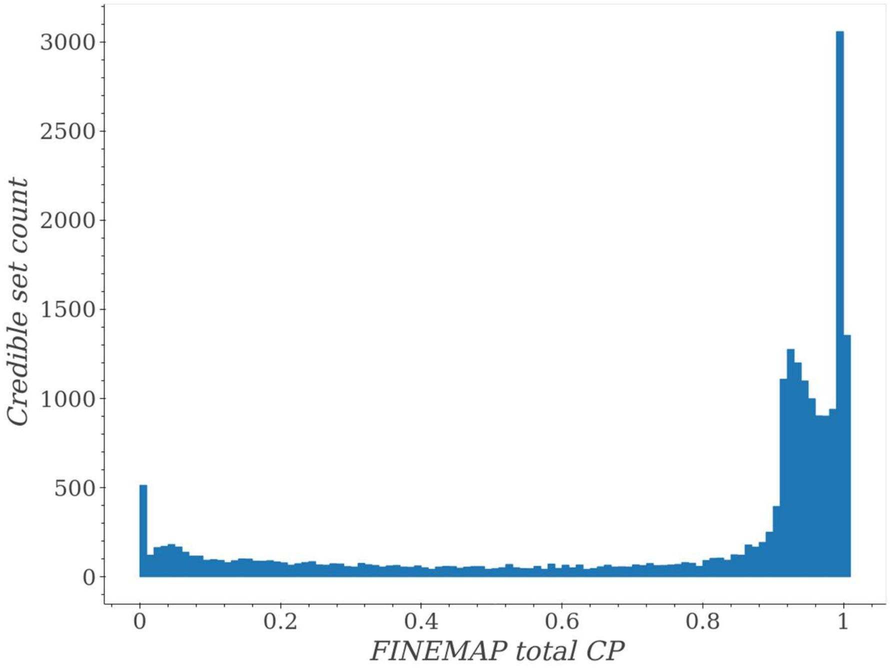


SuSiE 90%-credible sets across all trait-regions (with purity ≥ 0.8) were each binned by the total CP FINEMAP assigned to all variants in that set. Sets in the rightmost bin have FINEMAP total CP between 1 and 1.01 (i.e. FINEMAP predicts them to contain on average between 1 and 1.01 causal variants). FINEMAP assigned 6 SuSiE credible sets to have total CP greater than 1.01 (none of which attained total CP greater than 1.13); those 6 are omitted from the figure. By definition, SuSiE has estimated each 90%-credible set to have between a 90% and 100% chance of containing a single causal variant.

#### Supplementary Figure 9: Discordance between SuSiE and FINEMAP CPs

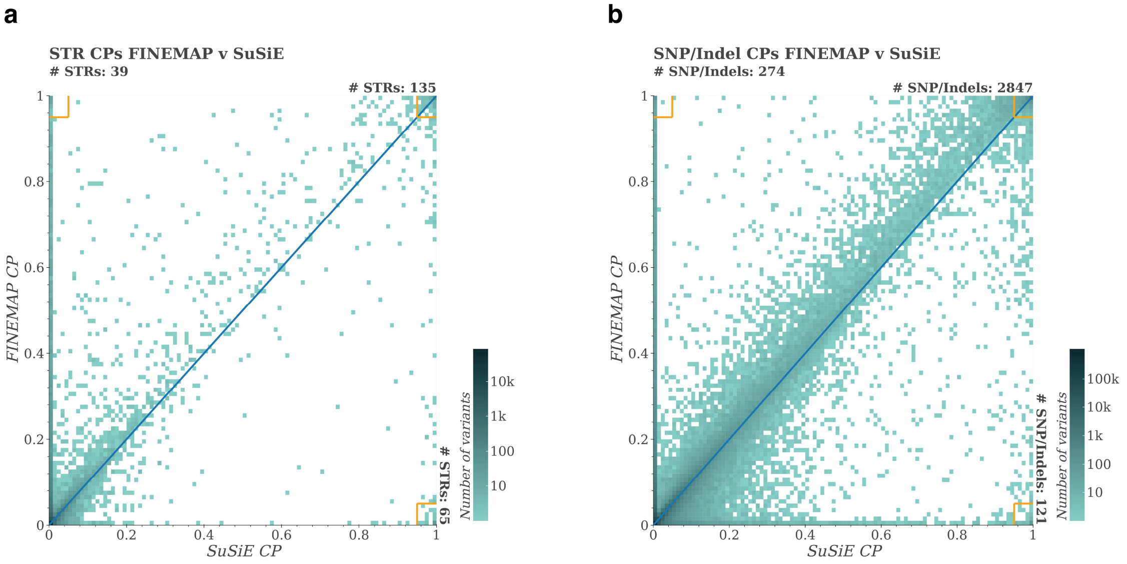


Comparison of CPs across all trait-regions between SuSiE (x-axis) and FINEMAP (y-axis) for genome-wide significant STRs **(a)** and SNPs and indels **(b)**. The blue line denotes equal CP. Yellow boxes in the three extreme corners are summarized by the number of variants residing in those boxes.

**Supplementary Figure 10: Discordance between fine-mappers**

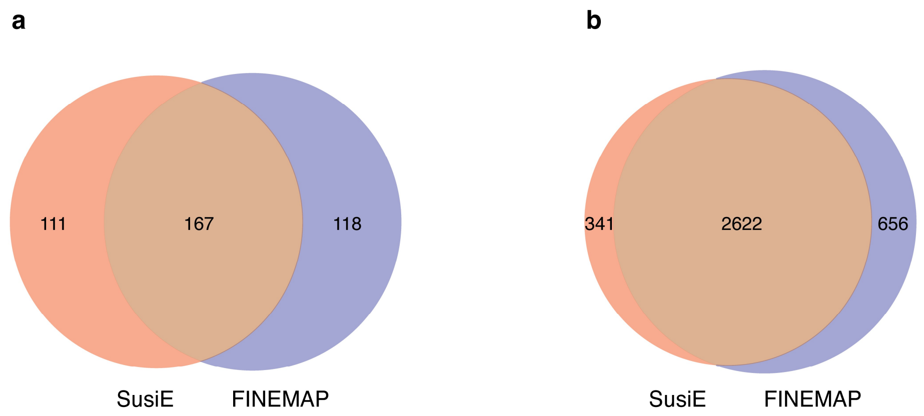


The number of **(a)** STRs and **(b)** SNPs/indels with p-value < 1e-10 assigned a CP ≥ 0.8 by only SuSiE (red), only FINEMAP (purple), or both (brown).

**Supplementary Figure 11: SuSiE settings not used for filtering**

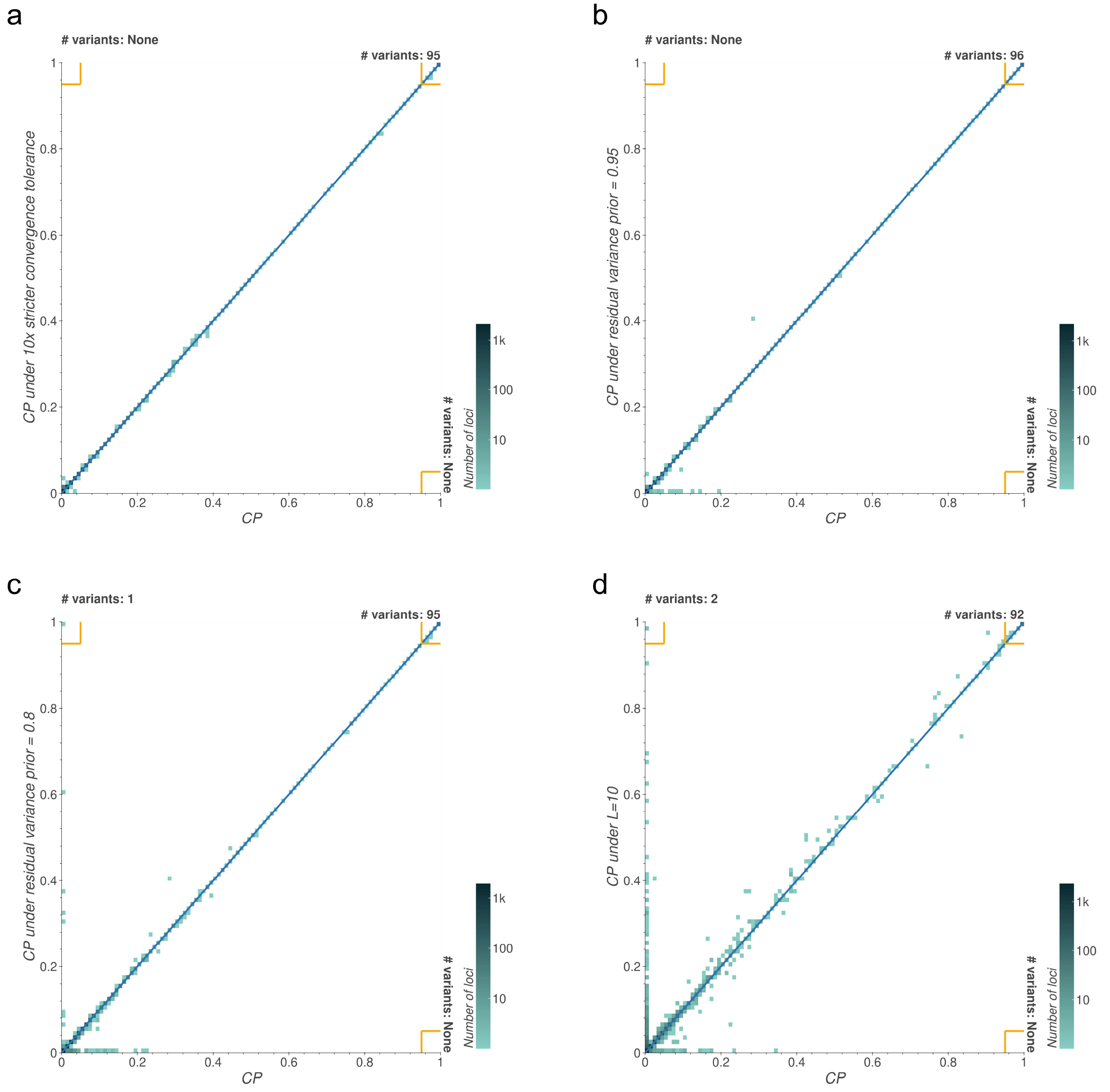


Concordance between SuSiE CPs for all genome-wide significant variants across most small-to-medium sized mean platelet volume fine-mapping regions under default settings on the x-axis (`tol=1e-3`, `residual_variance` slightly less than 1, and `L=50`) vs. a single alternate setting on the y-axis **(a)** `tol=1e-4`, **(b)** `residual_variance=0.95`, **(c)** `residual_variance=0.8` and **(d)** `L=10`. Blue lines denote equal CP. Yellow boxes in the three extreme corners are summarized by the number of variants residing in those boxes.

#### Supplementary Figure 12: Effect of best-guess genotypes on SuSiE results

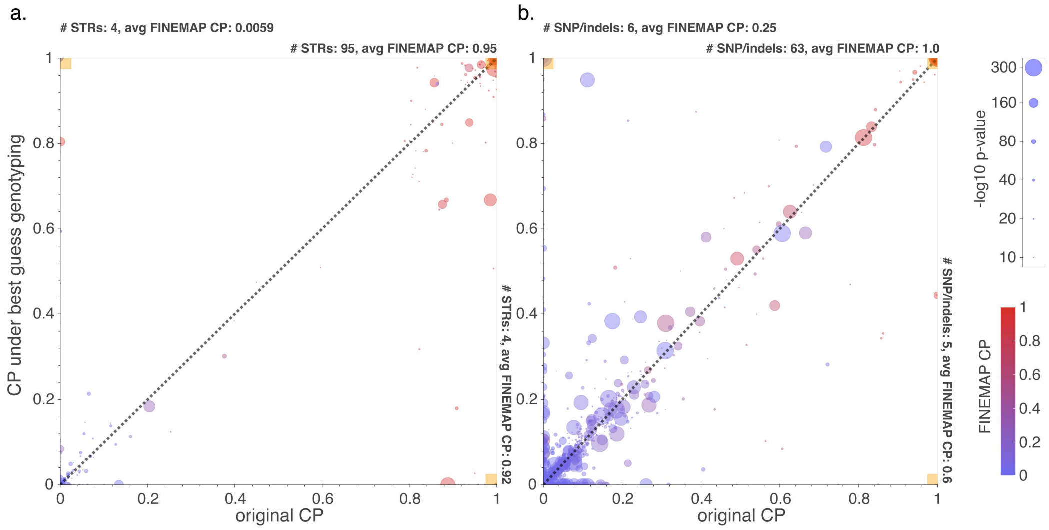


Discordance between SuSiE CPs for variants with p-value < 1e-10 when run with dosage genotypes (x-axis) vs best-guess genotypes (y-axis) among STRs **(a)** and SNPs and indels **(b)**. These data points are taken from running SuSiE on the trait-regions containing the 167 STR-trait associations with p-value < 1e-10 and with both SuSiE and FINEMAP CPs ≥ 0.8. Black lines denote equal CP. Larger circle sizes denote larger variant -$\log_{10}$ association p-values. Circle color denotes the CP of that variant from our default FINEMAP run. Yellow boxes in the three extreme corners are summarized by the number of variants residing in those boxes and the average FINEMAP CP value of those variants.

#### Supplementary Figure 13: Effect of alternate settings on FINEMAP results

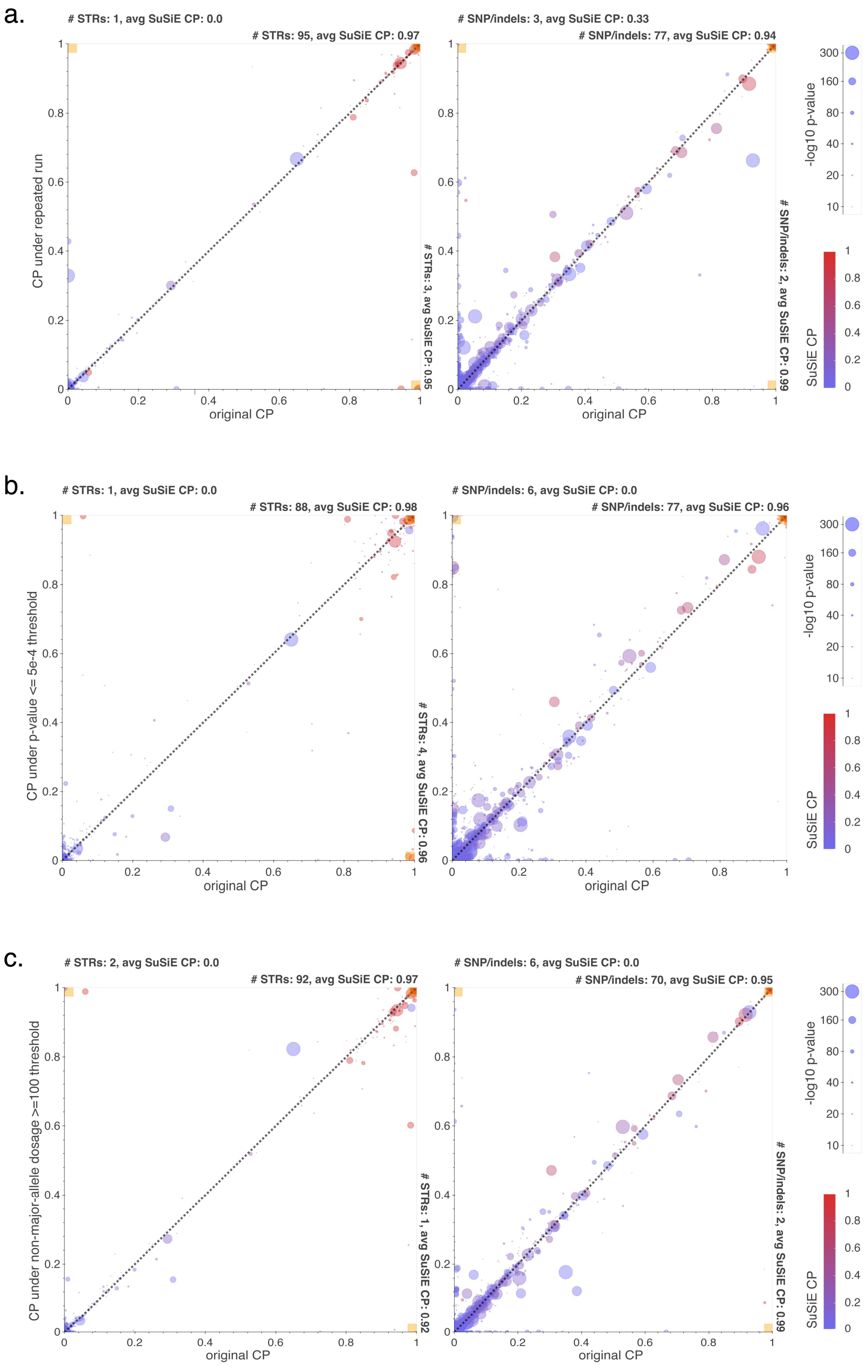

d.

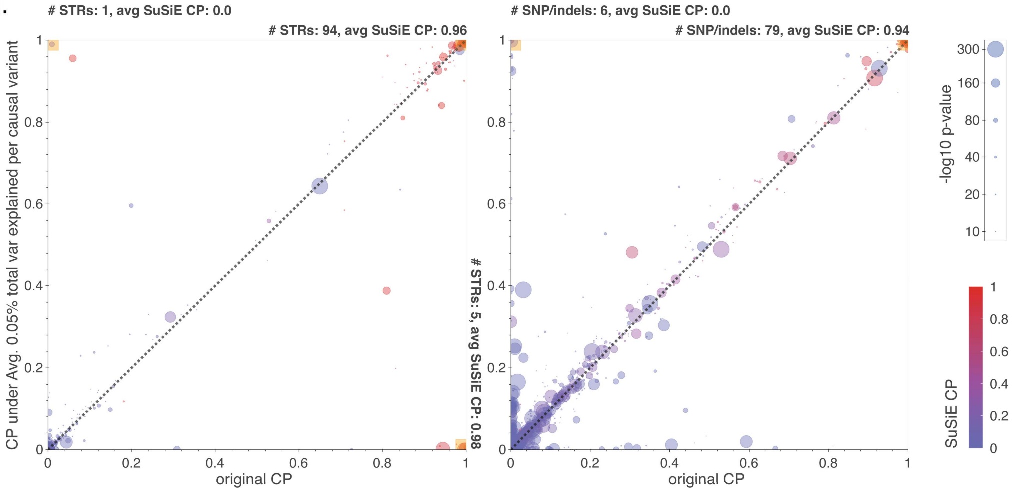

### STRs: 1, avg SuSiE CP: 0.0
### STRs: 94, avg SuSiE CP: 0.96
### STRs: 5, avg SuSiE CP: 0.98
### SNP/indels: 6, avg SuSiE CP: 0.0
### SNP/indels: 79, avg SuSiE CP: 0.94
CP under Avg. 0.05% total var explained per causal variant
original CP
-log10 p-value
300
160
80
40
20
10
SuSiE CP


e.

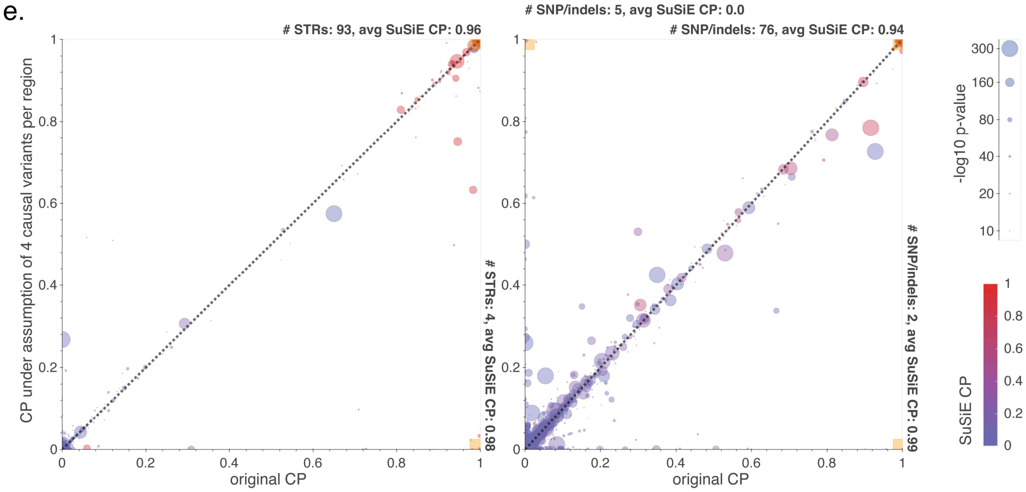

### STRs: 93, avg SuSiE CP: 0.96
### STRs: 4, avg SuSiE CP: 0.98
### SNP/indels: 5, avg SuSiE CP: 0.0
### SNP/indels: 76, avg SuSiE CP: 0.94
### SNP/indels: 2, avg SuSiE CP: 0.99
CP under assumption of 4 causal variants per region
original CP
-log10 p-value
300
160
80
40
20
10
SuSiE CP


f.

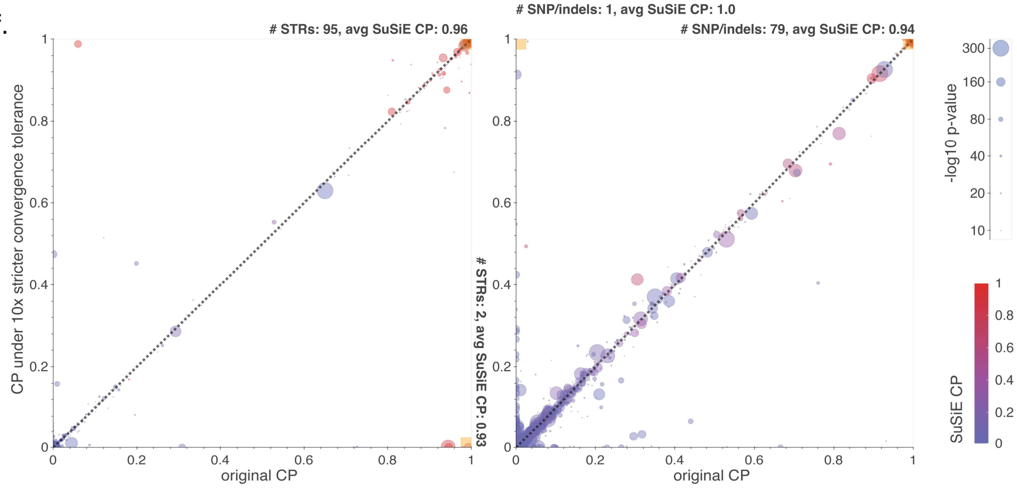

### STRs: 95, avg SuSiE CP: 0.96
### STRs: 2, avg SuSiE CP: 0.93
### SNP/indels: 1, avg SuSiE CP: 1.0
### SNP/indels: 79, avg SuSiE CP: 0.94
CP under 10x stricter convergence tolerance
original CP
-log10 p-value
300
160
80
40
20
10
SuSiE CP

g.

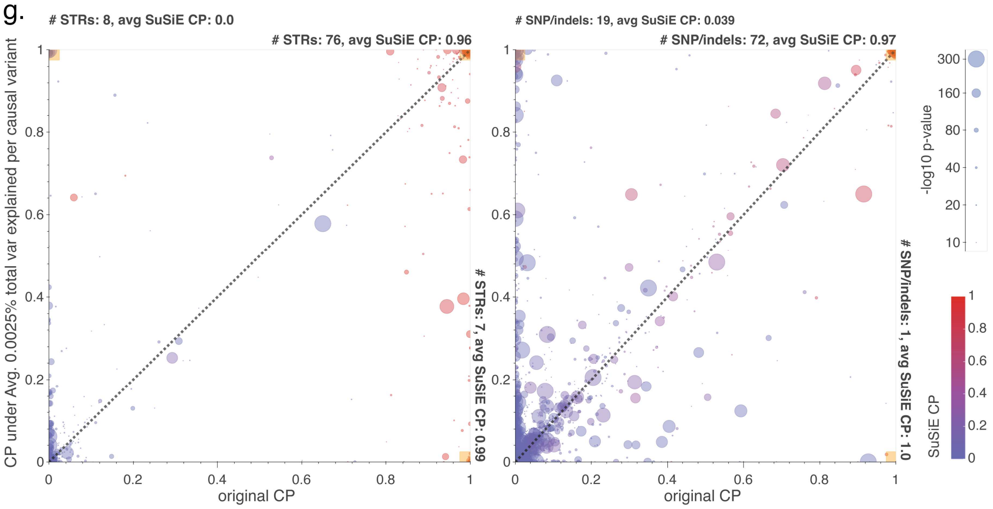


Discordance between FINEMAP CPs for variants with p-value < 1e-10 . x-axis values correspond to CPs from FINEMAP runs using our standard settings: with a p-value > 5e-2 filter, `--prior-std 0.05`, prior of one causal variant per trait-region and `--prob-conv-sss-tol 0.001` . y-axis values correspond to CPs from FINEMAP runs when **(a)** rerun with the same default settings on the same data (to account for random variation in the algorithm) or **(b-g)** run on the same data with a single alternate setting. Those alternate settings were: **(b)** p-value > 5e-4 filter **(c)** additionally filtering those variants with total non-major-allele dosage < 100 **(d)** `-prior-std 0.0224` **(e)** prior of four causal variants per trait region **(f)** `--prob-conv-sss-tol 0.0001` and **(g)** `--prior-std=0.005`. The variants in these plots are all those with p-value < 1e-10 in the trait-regions containing the 167 STR-trait associations with p-value < 1e-10 and with both SuSiE and FINEMAP CPs ≥ 0.8. CP discordance among STRs is plotted on the left, and among SNPs and indels is plotted on the right. Black lines denote equal CP. Larger circle sizes denote larger variant -$\log_{10}$ association p-values. Circle color denotes the CP of that variant from our default SuSiE run. Yellow boxes in the three extreme corners are summarized by the number of variants residing in those boxes and the average SuSiE CP value of those variants.

**Supplementary Figure 14: Effect of conservative prior favoring SNPs and indels on estimated causality of STR variants**

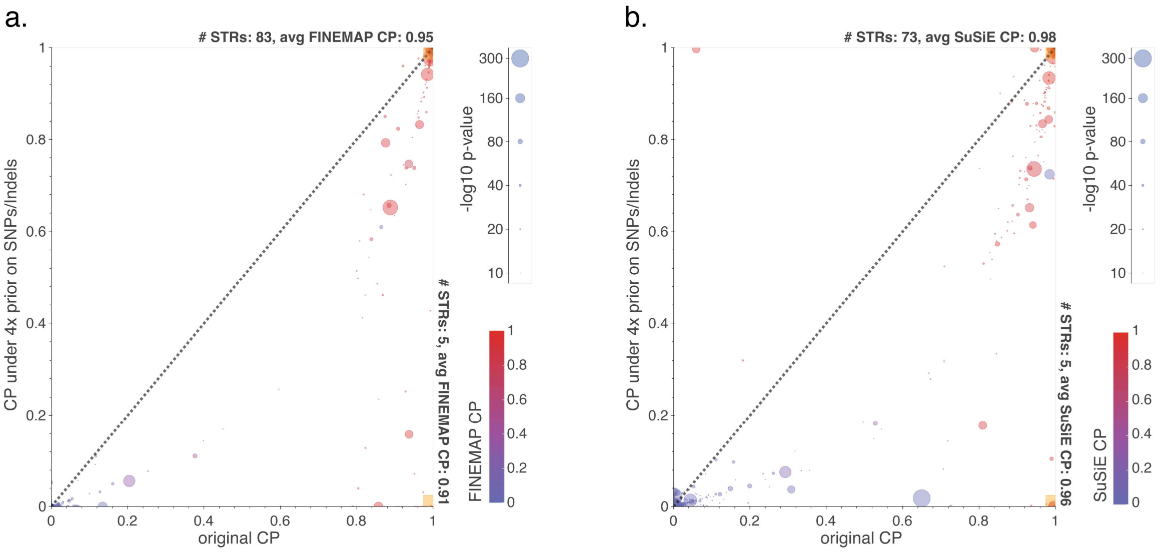


Discordance between CPs for STRs with p-value < 1e-10 when run under default settings (x-axis) vs with a 4x prior of causality for SNPs and indels as compared to STRs (y-axis) in **(a)** SuSiE and **(b)** FINEMAP. These data points are taken from running SuSiE and FINEMAP on the trait-regions containing the 167 STR-trait associations with p-value < 1e-10 and with both SuSiE and FINEMAP CPs ≥ 0.8. Black lines denote equal CP. Larger circle sizes denote larger variant -$\log_{10}$ association p-values. Circle color denotes the CP of that variant from the other fine-mapper's default run. Yellow boxes in the extreme corners are summarized by the number of variants residing in those boxes and the average SuSiE CP value of those variants.

**Supplementary Figure 15: Replication of White British STR associations in other White populations**

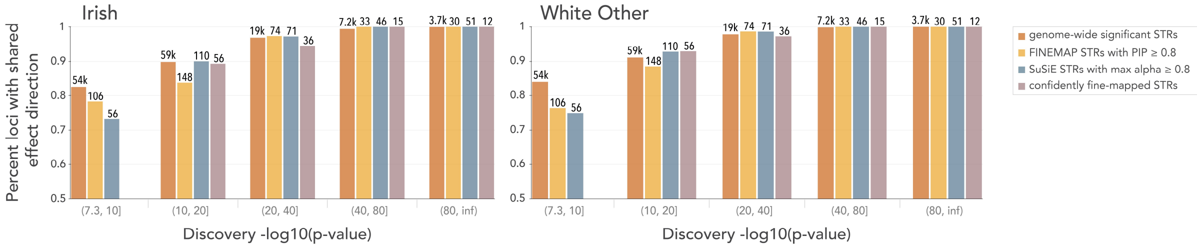


The y-axis gives the fraction of STR associations measured in the discovery cohort that have the same direction of effect when measured in the replication population regardless of p-value (left=Irish, right=White Other, see **Fig. 3** for the non-White populations). Brackets beneath the x-axis denote the binning of discovery -$\log_{10}$ p-values. Brown=genome-wide significant associations (discovery p<5e-8), orange=FINEMAP fine-mapped STR associations (discovery p<5e-8 and FINEMAP CP ≥ 0.8), teal=SuSiE fine-mapped STR associations (discovery p<5e-8 and SuSiE CP ≥ 0.8) and purple=confidently fine-mapped STR associations. Annotations above each bar indicate the number of STR-trait associations considered. We required confidently fine-mapped STR associations to have p-value < 1e-10, thus they do not appear in the left-most bin. The trends in these figures are somewhat sensitive to the choice of p-value bin boundaries so we additionally analyze this data using logistic regression models (**Supplementary Table 10**).

**Supplementary Figure 16: Evaluation of E2F4 protein structure using AlphaFold**

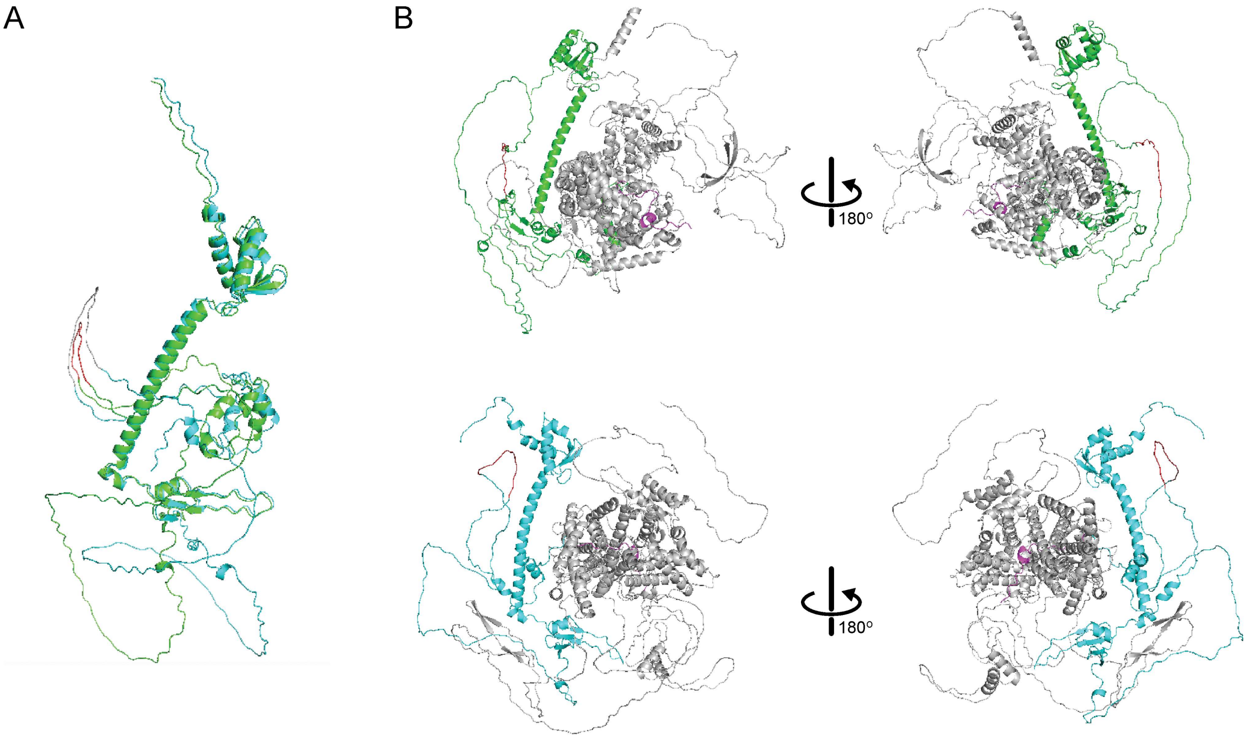


**(a)** depicts the E2F4 protein, while **(b)** depicts the E2F4-RBL2 complex. In both, green denotes the E2F4 protein variant with the shorter STR allele (12 units) while cyan represents the E2FF4 protein containing longer, reference STR allele (19 units). In **(a)** the shorter variant's poly S region coded by the STR is highlighted in red, and the longer variant's poly S region is highlighted in grey. In **(b)**, both poly S regions are highlighted in red, and RBL2 is in grey. Notice in **(a)** that the structure of the two E2F4 variants are highly similar but that in **(b)** the distance between E2F4 and RBL2 is smaller for the shorter-allele variant than the longer-allele variant.

**Supplementary Figure 17: Prevalence of STR features**

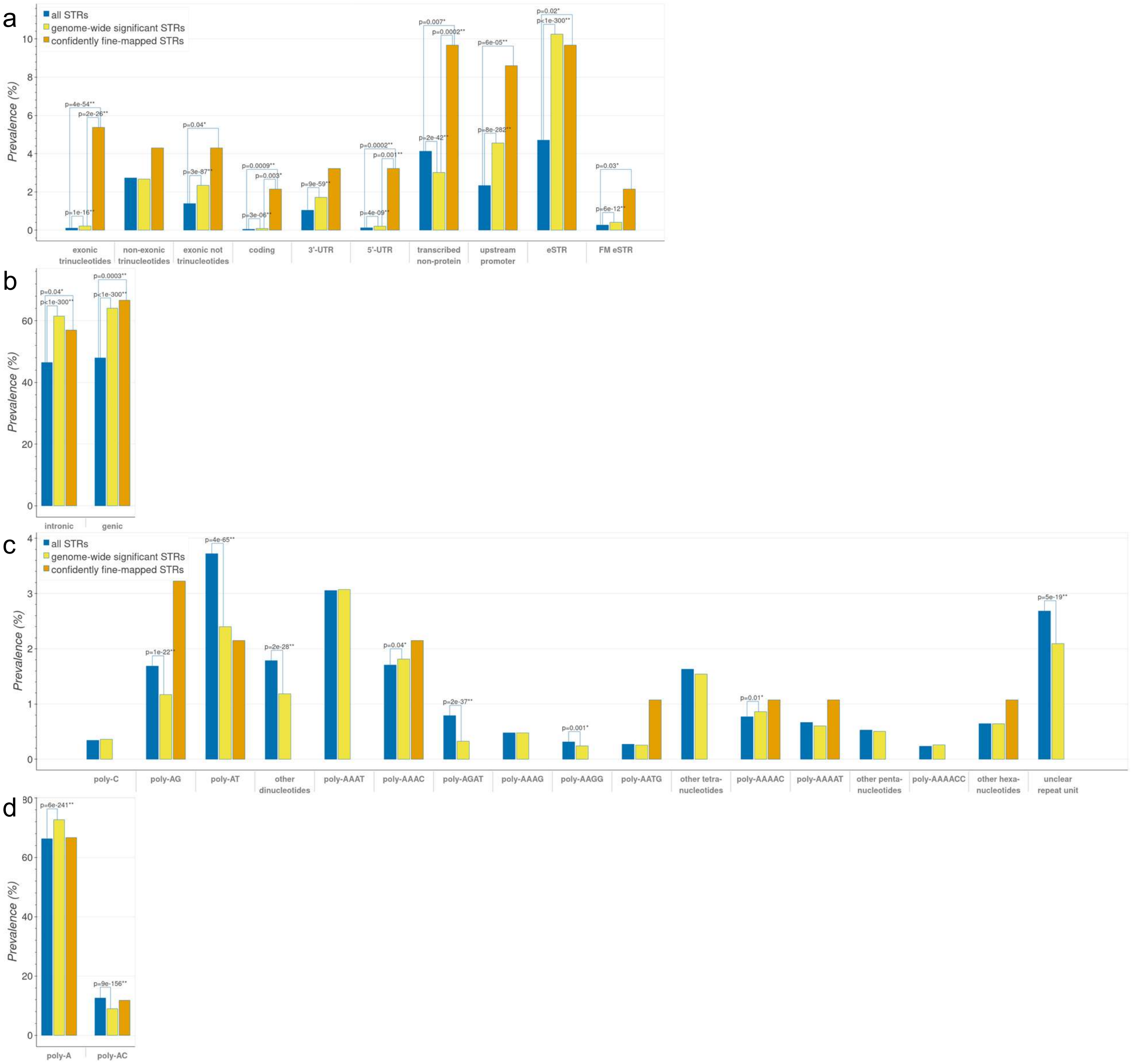


Genomic annotation **(a-b)** and repeat unit **(c-d)** prevalences are shown for different categories of STRs. (Blue=all STRs in our imputation panel, yellow=genome-wide significant STRs for at least one trait, orange=confidently fine-mapped STRs). **(a,c)** show low prevalence categories, **(b,d)** show high prevalence categories. In **(a)**, "upstream promoter" is defined as the region 3kb upstream of a transcription start site, and "eSTR" and "FM eSTR" are categories from our previous study to identify STR-gene expression associations in the Genotype Tissue Expression (GTEx) cohort[23]. **(c-d)** contains all repeat units represented by at least one thousand STRs in our reference panel, except for trinucleotide STR repeat units, as enrichments for those could not be distinguished from the enrichment for exonic trinucleotide STRs as a whole. See the **Methods** for more details. P-values from two-sided tests of difference between proportions are only displayed when $p \leq 0.05$. Note that strong p-values for differences between the all STRs and genome-wide significant STRs categories could often be due to restricting to phenotypically-important genomic regions and not necessarily due to enrichment for causal variants.

**Supplementary Figure 18: Confidently fine-mapped STRs influencing DNA methylation**

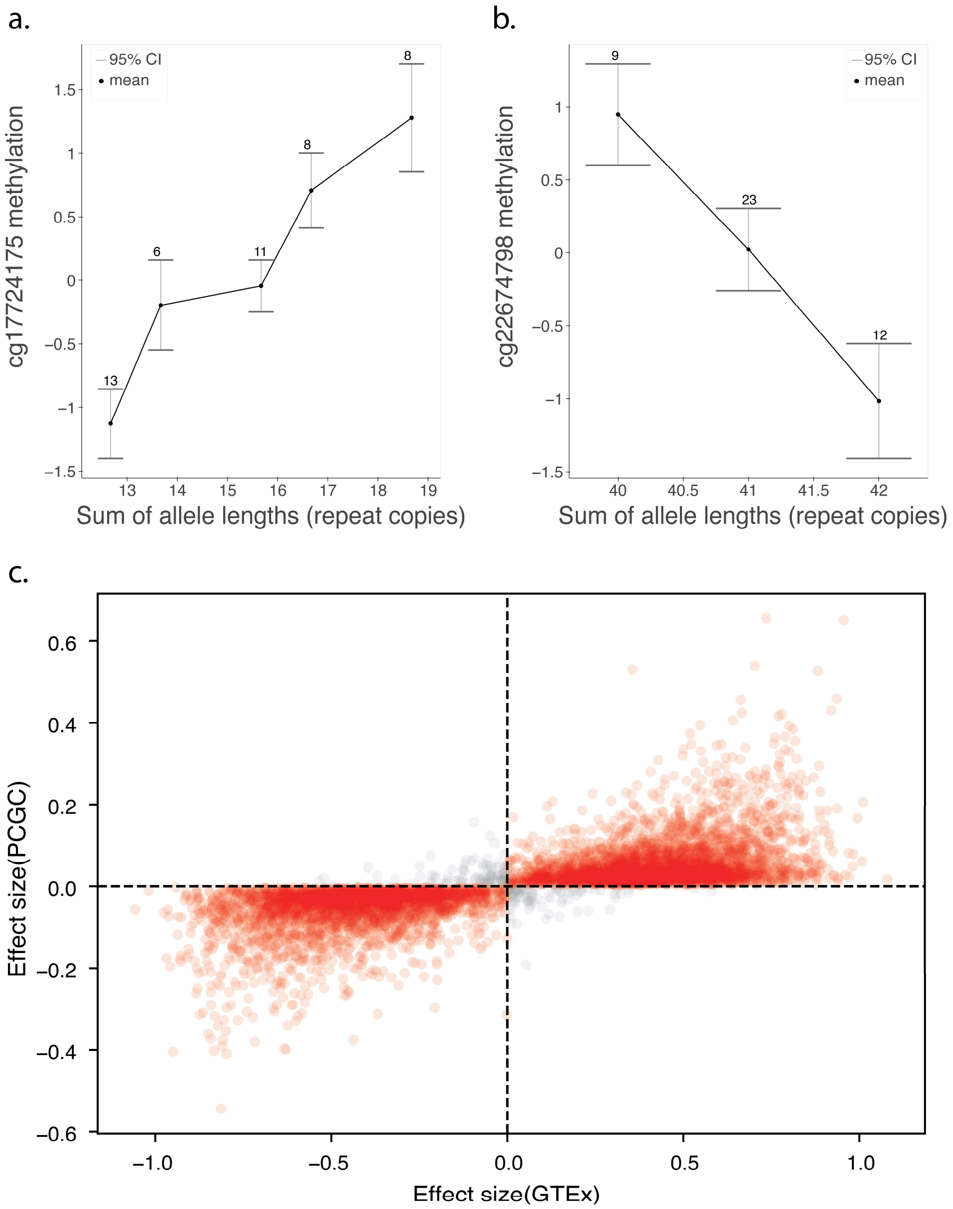


**(a)** STR 1:150579759-150579814 (hg38), residing less than 200bp upstream from the transcription start site of the gene *MCL1* and confidently fine-mapped to mean platelet volume (p=3e-16), is associated with methylation of a CpG site 500bp farther upstream (cg17724175). **(b)** STR 1:3170058-3170078 (hg38), residing in a conserved region of an intron of the gene *PRDM16* and confidently fine-mapped to platelet crit (p=4e-12), is associated with the methylation of a CpG site ~10kb away that resides in the same intron (cg22674798). Summed allele lengths are on the x-axis, and inverse normalized methylation levels are on the y-axis. Box plots indicate

25th, median and 75th percentiles, with whiskers extending up to 1.5 times the interquartile range beyond the 25th and 75th percentiles. Circles denote the mean methylation values for each allele length sum. **(c)** The effect sizes from STR length vs CpG methylation associations as measured here in the GTEx cohort compared to those measured by Martin-Trujillo et al.[24] in the Pediatric Cardiac Genomics Consortium (PCGC) cohort. Each of the n=7661 circles represents one of the associations measured in both cohorts. Red/grey circles denote matching/opposite effect directions in the two cohorts. We see that effects are broadly consistent ($r=0.73$, $p<10^{-200}$) between the two studies.

**Supplementary Figure 19: Association of an STR in *SLC2A2* with bilirubin levels**

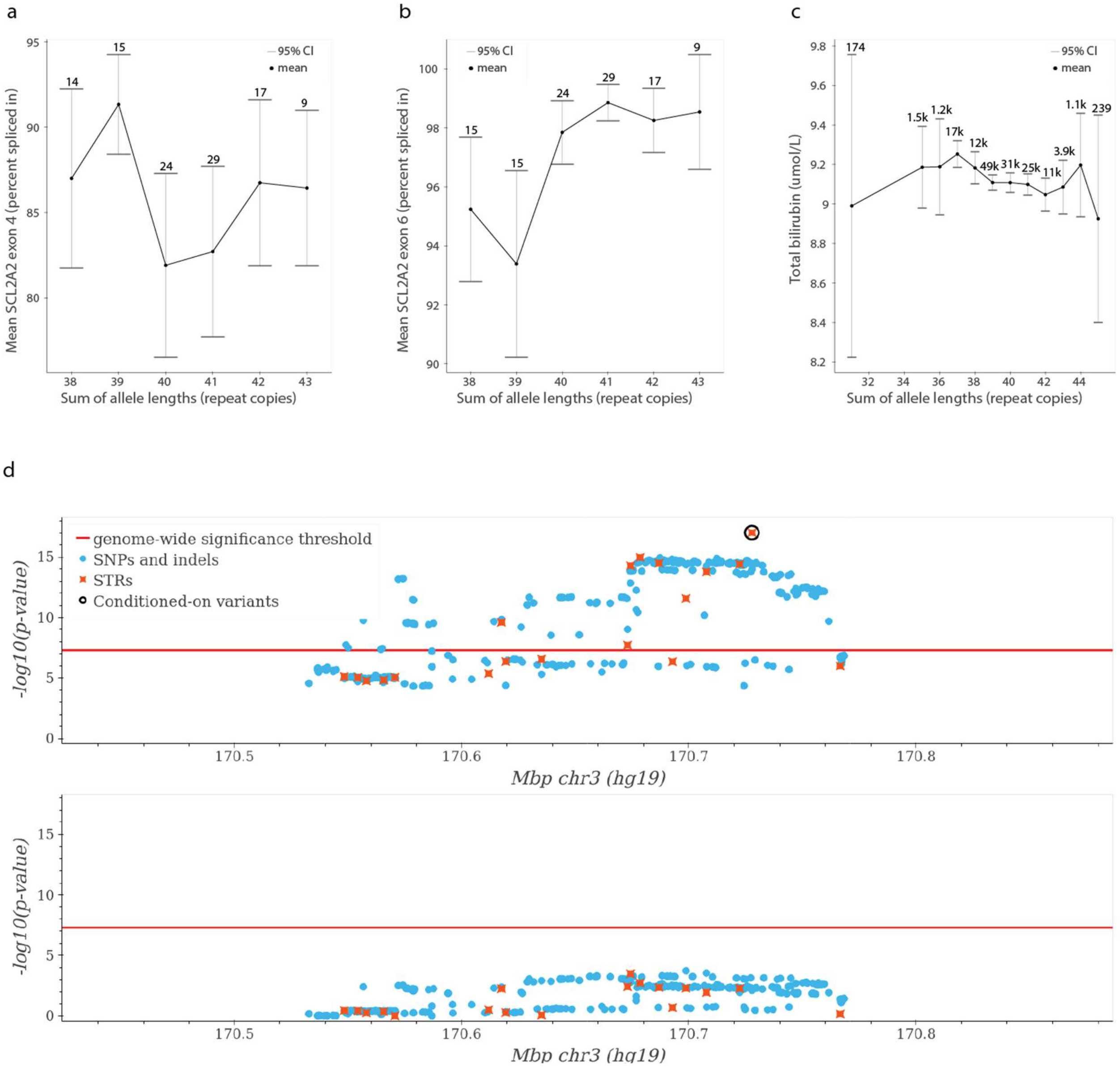


Scatter plots showing the association between the GT repeat at chr3:170727702 (hg19) and the splicing (percent spliced in) of exon 4 (**a**; linear association p=0.77) and exon 6 (**b**; linear association p=8.7e-07) of *SLC2A2* in Liver samples from GTEx[25]. For each plot, the x-axis represents the sum of repeat copies of STR in each individual and the y-axis represents percent spliced in for the indicated exon. The solid line gives the mean percent spliced in. Population counts are displayed for each length sum, only length sums with a count of at least 5 are displayed. **(c)** Association between length (from whole genome sequencing) of the GT repeat and eosinophil percentage in the UKB. The mean trait value for each sum of STR allele lengths was calculated across QCed White British participants. 95% confidence intervals were calculated similarly. Only allele length sums with a population frequency of 0.1% or greater are displayed. Rounded population-wide counts are displayed for each sum. **(d)** Association of variants at the *SLC2A2* locus and total bilirubin levels, before (top) and after (bottom) conditioning on the GT repeat. Light blue=SNPs and indels; orange=STRs. Red line=significance threshold, black circle=the $(GT)_n$ STR.

**Supplementary Figure 20: Associations of an STR in *CBL* with platelet crit and residual platelet volume**

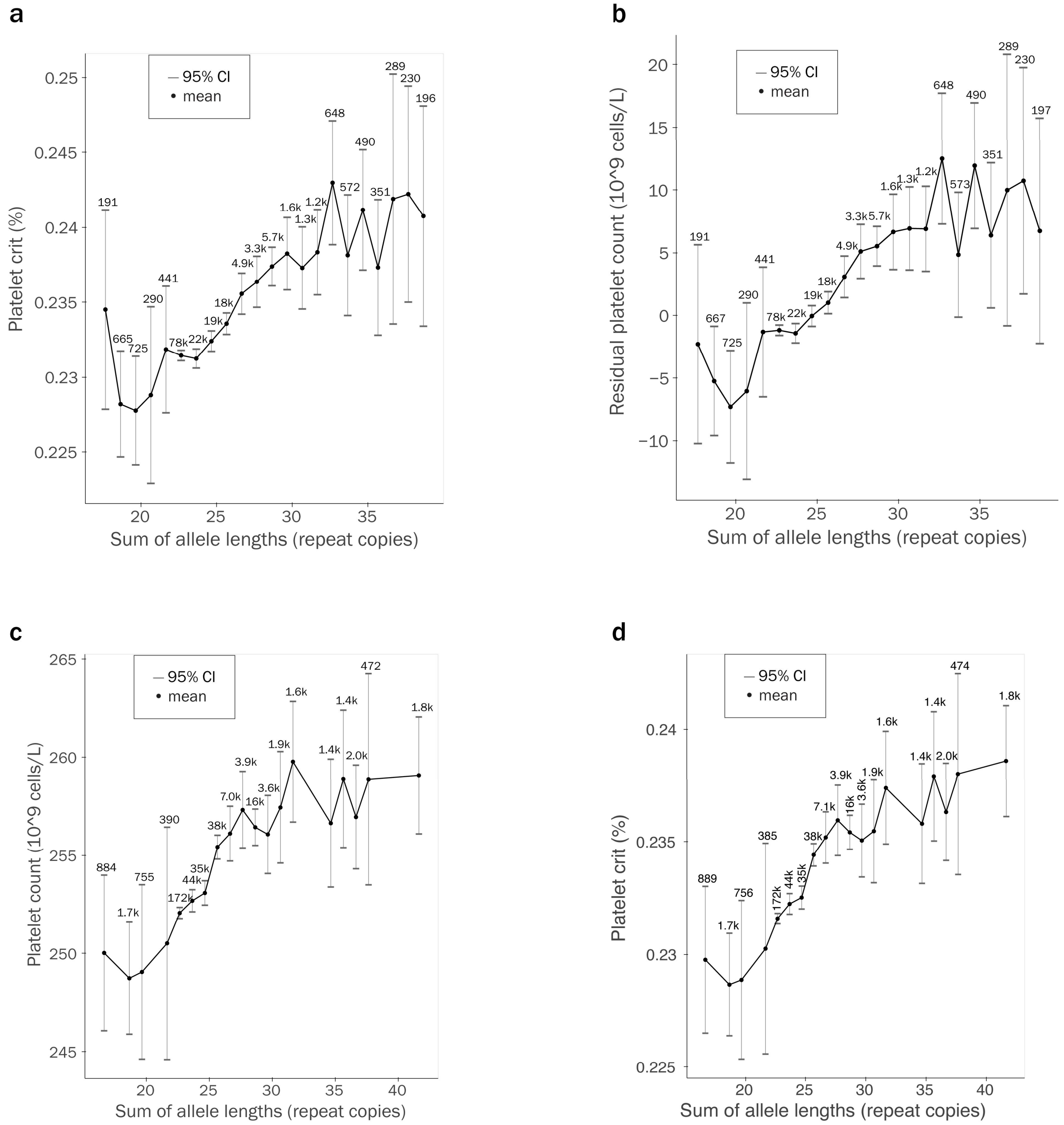


Summed STR length (from whole genome sequencing) vs platelet crit **(a)** and residual platelet count **(b)**. The mean trait value for each sum of STR allele lengths was calculated across QCed White British participants. Rounded population-wide counts are displayed for each sum. Only allele length sums with a population frequency of 0.1% or greater are displayed.

The trends in **(a-b)** are nearly identical to those in **Fig. 4b** for unadjusted platelet count. For **(b)** we calculated residuals by linearly regressing out the same covariates that were used in association p-value calculations (**Methods**), including sex, age, population principal components and categorical covariates for batch effects. We then calculated the mean residual for each allele length sum. Note that in our association pipeline, covariates are included as we test STRs for association with rank inverse normalized phenotypes, while for **(b)** here we did not rank inverse normalize the phenotype values.

**(c-d)** display the associations with platelet count and platelet crit, respectively, of STR lengths derived from imputation. These trends are overall similar to those with genotypes derived from WGS data (**Fig. 4b**, part **(a)** of this figure). For **(c-d)** we calculate the mean trait value for each allele length sum across QCed, unrelated White British participants, where each participant's contribution to each allele length sum's mean is weighted by that participant's imputed likelihood of having that allele length sum genotype. 95% confidence intervals were calculated similarly.

**Supplementary Figure 21: Associations of an STR in *CBL* with mean sphered cell volume**

a

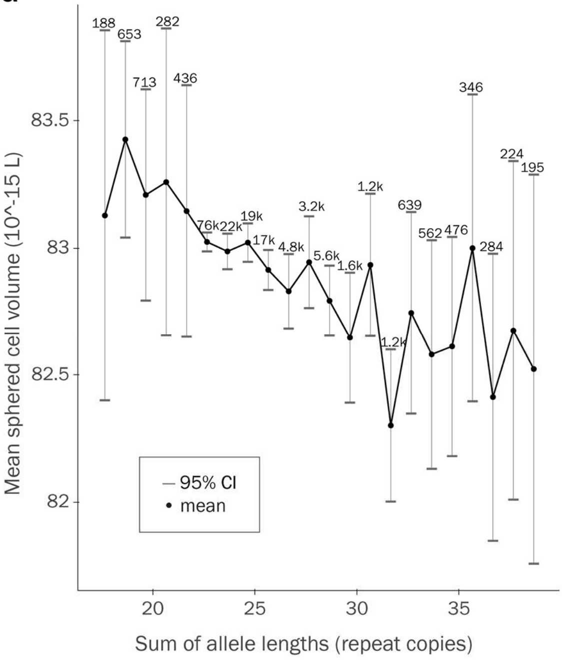


b

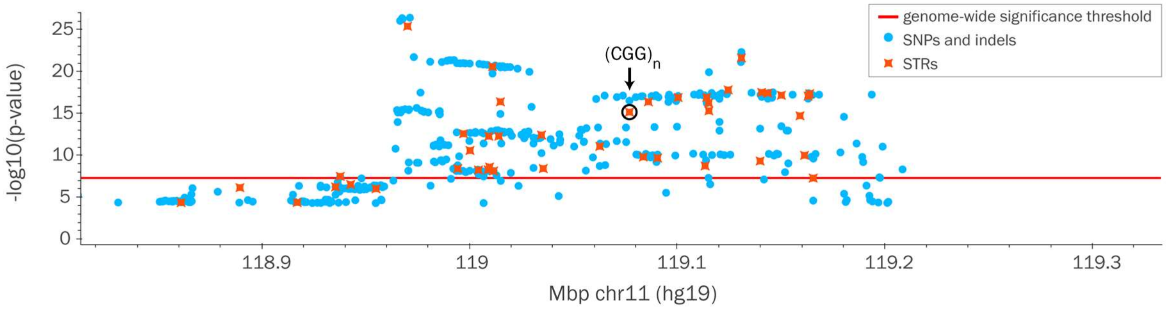


**(a)** Association between length (from whole genome sequencing) of the CGG repeat at chr11:119077000 (hg19) and mean sphered cell volume. The mean trait value for each sum of STR allele lengths was calculated across QCed White British participants. 95% confidence intervals were calculated similarly. Only allele length sums with a population frequency of 0.1% or greater are displayed. Rounded population-wide counts are displayed for each sum. **(b)** Association of variants at the *CBL* locus and mean sphered cell volume. Light blue=SNP and indels; orange=STRs. Red line=significance threshold, black circle=the $(CGG)_n$ STR.

**Supplementary Figure 22: Distribution of alleles of an STR in *CBL* across 1000 Genomes populations**

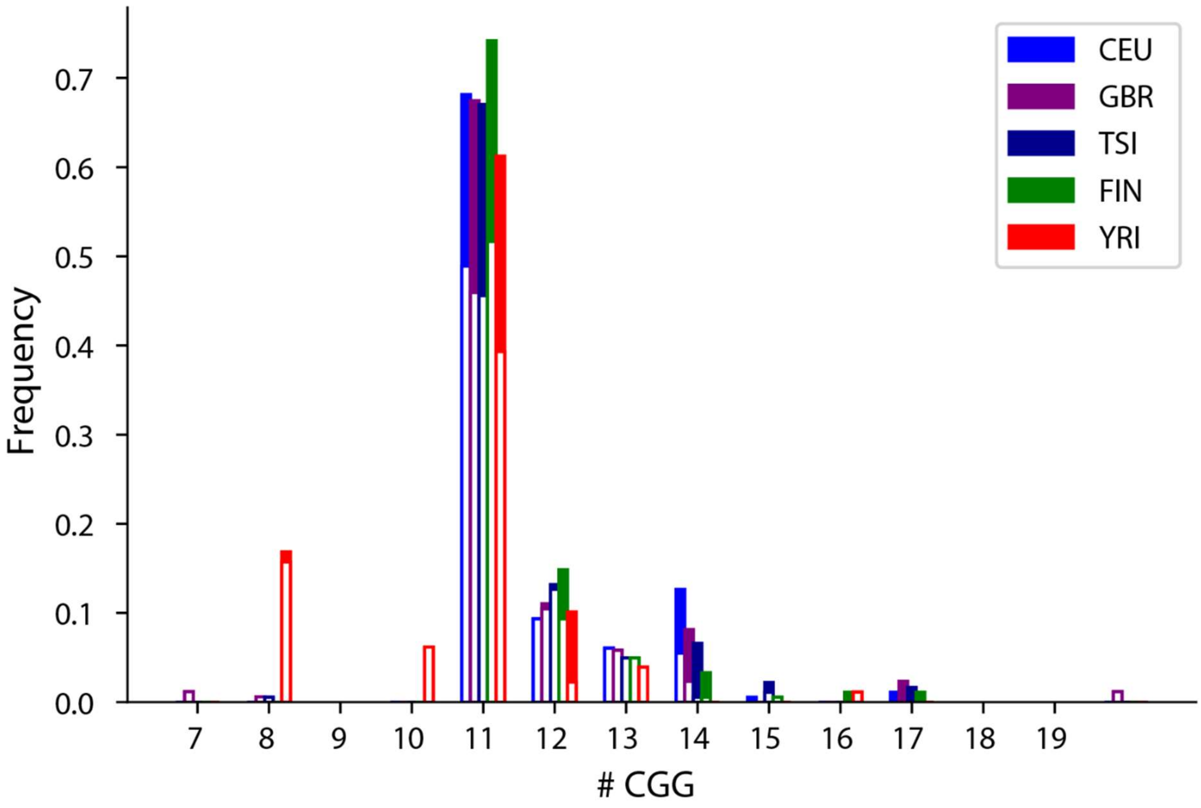


The x-axis gives STR length (number of repeat units) and y-axis gives the population frequency. The solid portion of each bar corresponds to the alleles of that length that include a "TGG" imperfection at the second repeat (rs7108857). Colors denote 1000 Genomes populations that were included in the Geuvadis cohort[26].

**Supplementary Figure 23: Association of an STR in *BCL2L11***

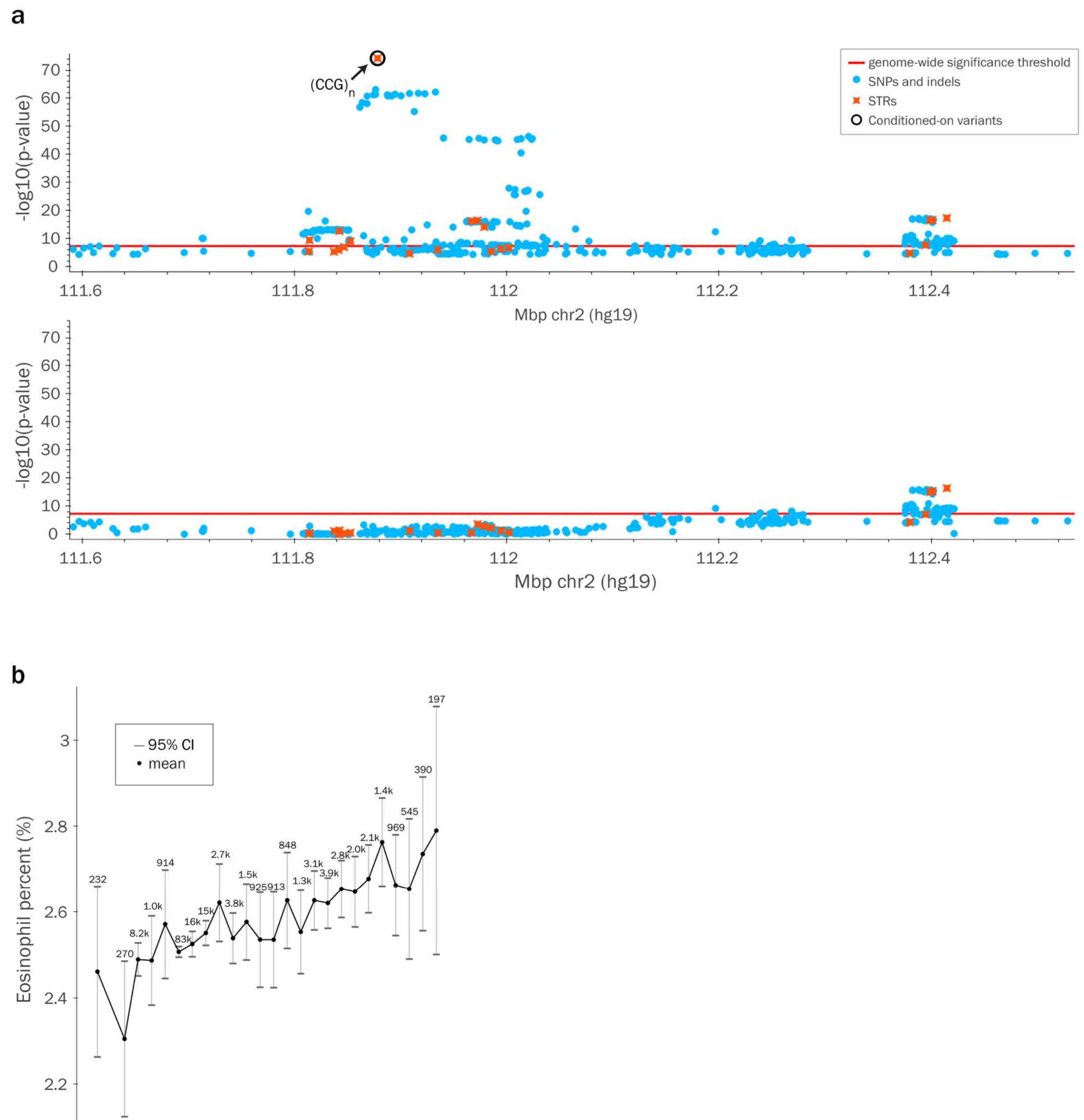


**(a)** Association of variants at the *BCL2L11* locus and eosinophil percent, before (top) and after (bottom) conditioning on the CCG repeat at chr2:111878544 (hg19). Light blue=SNPs and indels; orange=STRs. Red line=significance threshold, black circle=the $(CCG)_n$ STR. **(b)** Association between length (from whole genome sequencing) of the CCG repeat and eosinophil percentage. The mean trait value for each sum of STR allele lengths was calculated across QCed White British participants. 95% confidence intervals were calculated similarly. Only allele length sums with a population frequency of 0.1% or greater are displayed. Rounded population-wide counts are displayed for each sum.

#### Supplementary Figure 24: Association of an STR in *TAOK1*

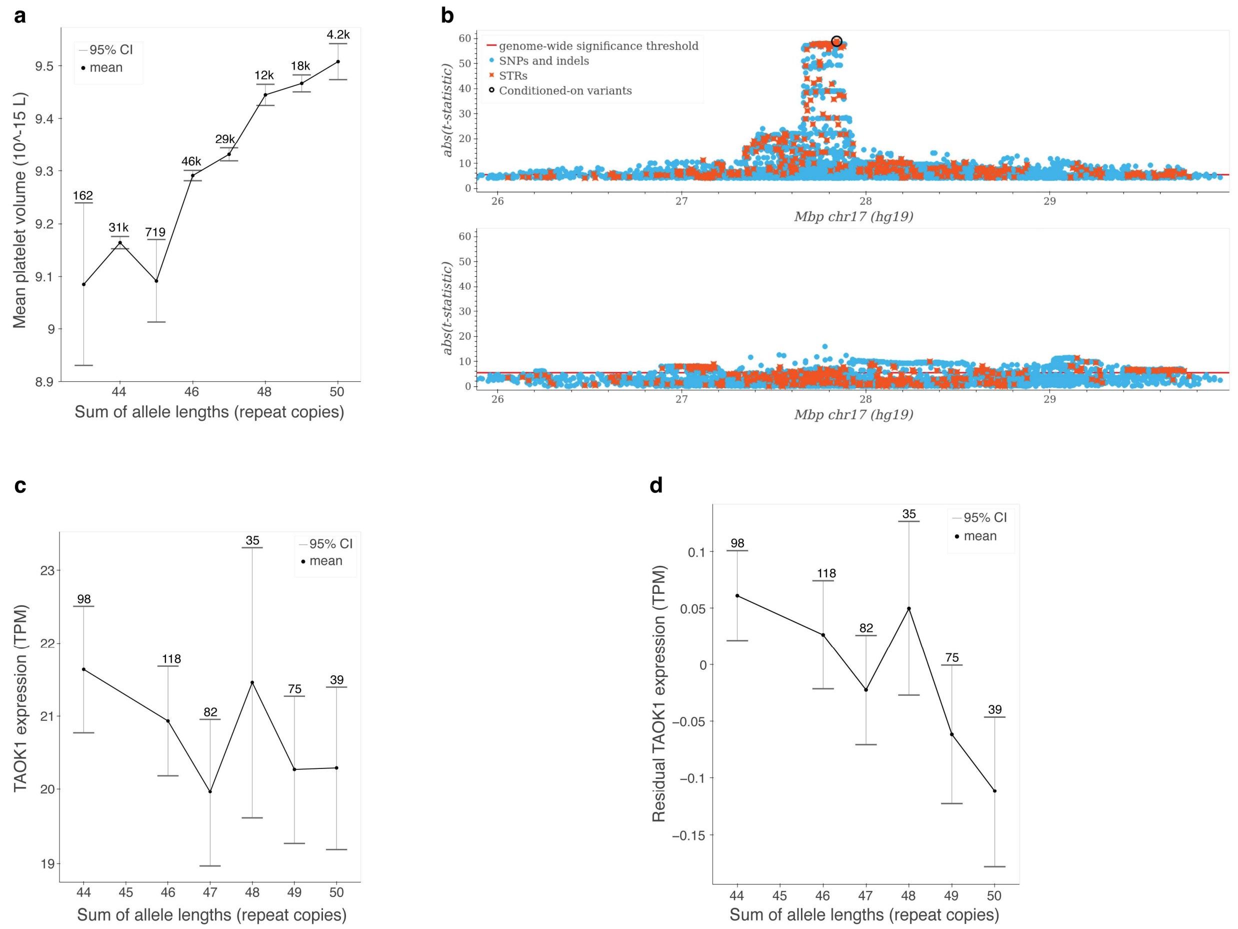


**(a)** Association between length (from whole genome sequencing) of the A repeat at chr17:27842016 (hg19) and mean platelet volume. The mean trait value for each sum of STR allele lengths was calculated across QCed White British participants. 95% confidence intervals were calculated similarly. Only allele length sums with a population frequency of 0.1% or greater are displayed. Rounded population-wide counts are displayed for each sum. **(b)** Association of variants at the *TAOK1* locus and mean platelet volume before (top) and after (bottom) conditioning on the STR. Light blue=SNPs and indels; orange=STRs. Red line=significance threshold, black circle=the $(A)_n$ STR. In this Manhattan plot we display associations according to the absolute value of their t-statistics instead of their -log10 p-values as those p-values exceeded the precision of our software (<1e-300). **(c-d)** Association between imputed best-guess genotypes of the repeat and *TAOK1* gene expression in thyroid tissue in the GTEx cohort[25]. Population-wide counts are displayed for each allele length sum. **(c)** displays the association with *TAOK1* expression

TPM, while **(d)** displays the association with residual TPM values obtained after regressing out genetic principal components and PEER factors.

#### Supplementary Figure 25: Association and allele distribution of an STR in *ESR2*

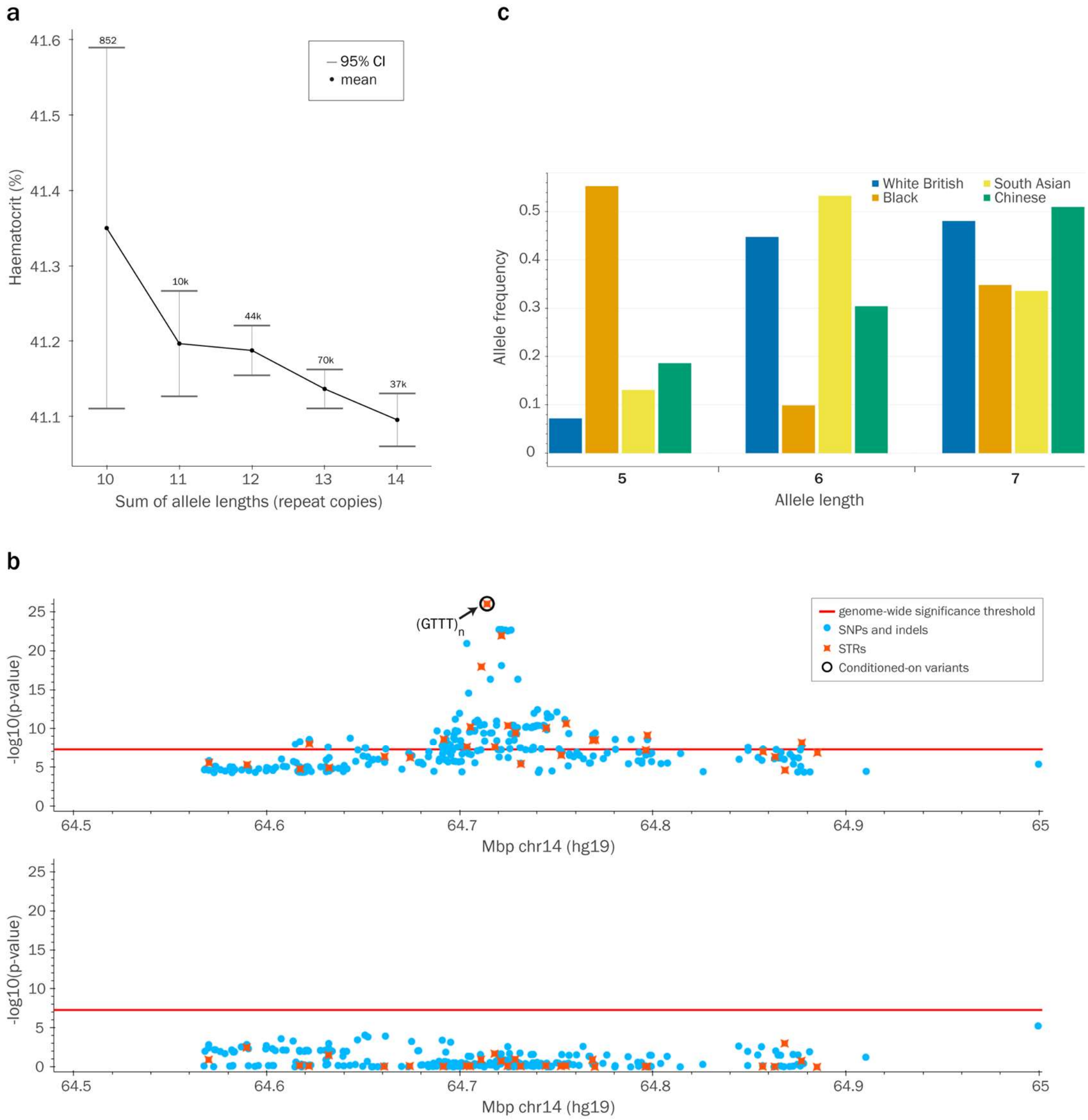


**(a)** Association between length (from whole genome sequencing) of the GTTT repeat at chr14:64714051 (hg19) and haematocrit. The mean trait value for each sum of STR allele lengths was calculated across QCed White British participants. 95% confidence intervals were calculated similarly. Only allele length sums with a population frequency of 0.1% or greater are displayed. Rounded population-wide counts are displayed for each sum. **(b)** Association of variants at the *ESR2* locus and haematocrit. Conditioning on the repeat fully accounts for the signal seen in this region. Light blue=SNPs and indels; orange=STRs. Red line=significance threshold, black circle=the $(GTTT)_n$ STR. **(c)** Distribution of STR length alleles from whole genome sequencing in different populations (blue=White British, orange=Black, yellow=South Asian; green=Chinese). Length alleles with frequency < 0.1% in all populations have been omitted.

**Supplementary Figure 26: Association of an STR in *NCK2***

a

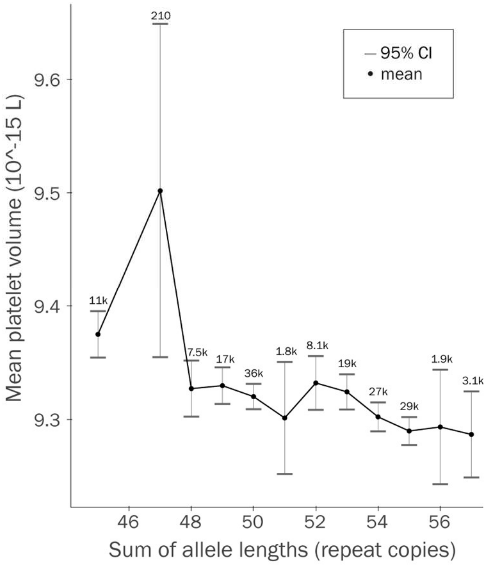


b

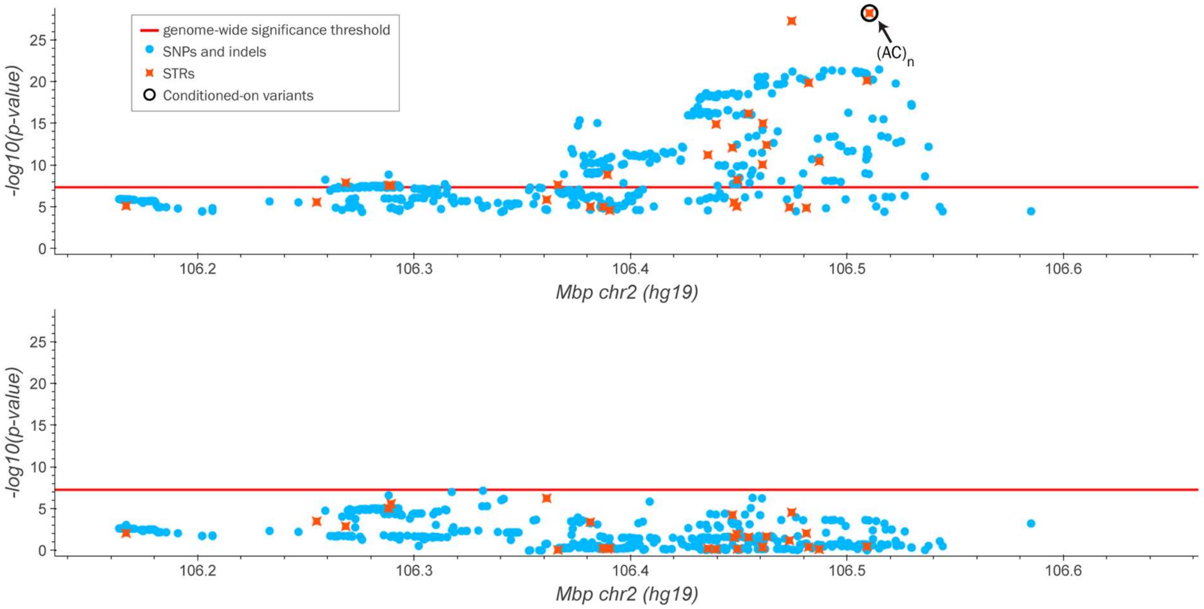


**(a)** Association between length (from whole genome sequencing) of the AC repeat at chr2:106510441 (hg19) and mean platelet volume. The mean trait value for each sum of STR allele lengths was calculated across QCed White British participants. 95% confidence intervals were calculated similarly. Only allele length sums with a population frequency of 0.1% or greater are displayed. Rounded population-wide counts are displayed for each sum. **(b)** Association of variants at the *NCK2* locus and mean platelet volume. Conditioning on the repeat fully accounts for the signal seen in this region. Light blue=SNPs and indels; orange=STRs. Red line=significance threshold, black circle=the $(AC)_n$ STR.

#### Supplementary Figure 27: Associations of an STR in *RHOT1*

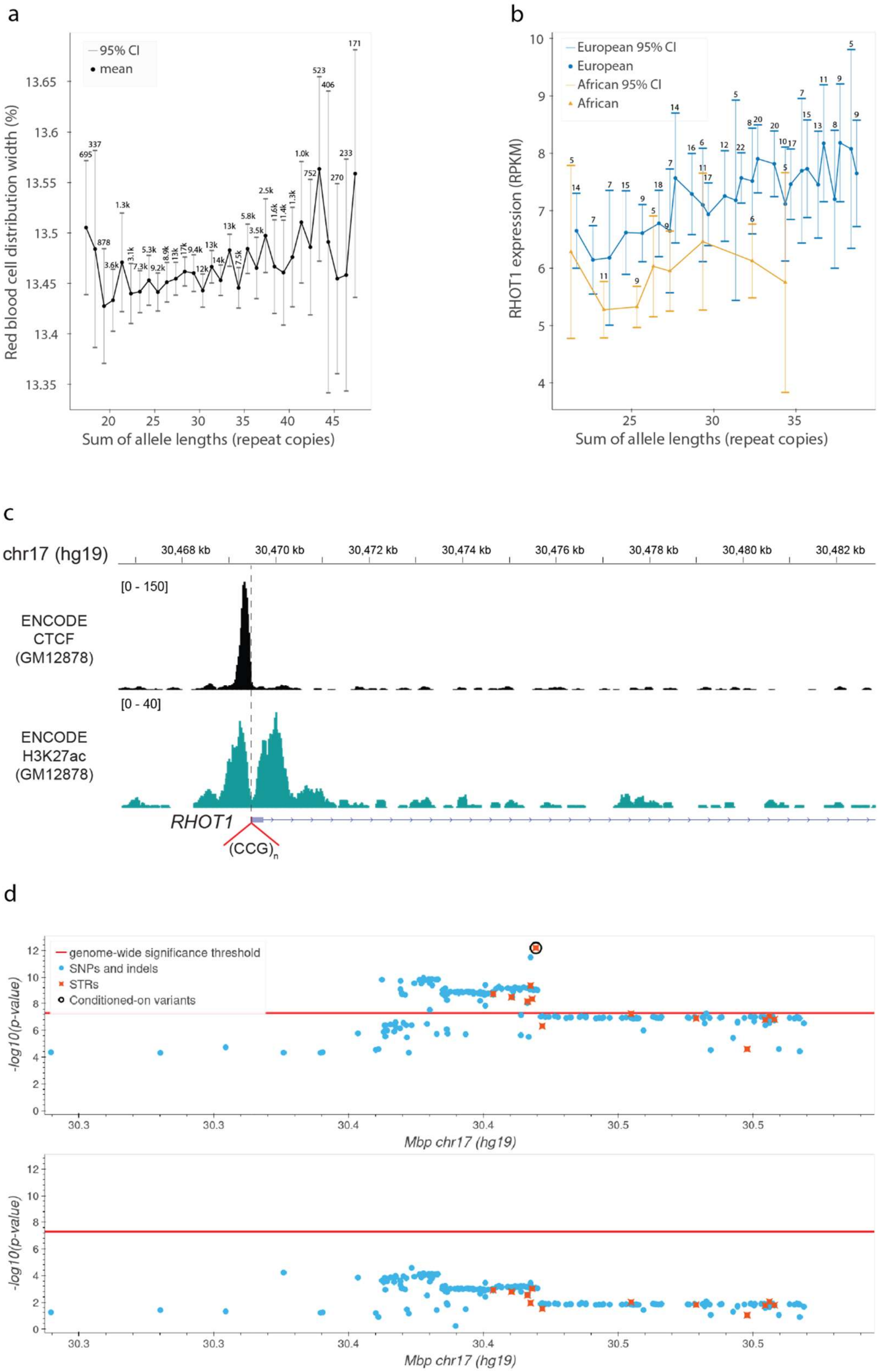

**(a)** Association between length (from UKB whole genome sequencing) of the CCG repeat at chr17:30469471 (hg19) and red blood cell distribution width. The mean trait value for each sum of STR allele lengths was calculated across QCed White British participants. 95% confidence intervals were calculated similarly. Only allele length sums with a population frequency of 0.1% or greater are displayed. Rounded population-wide counts are displayed for each sum. **(b)** Association between dosage of the repeat and *RHOT1* gene expression in the Geuvadis cohort[26] (LCLs; n=447). Solid lines give mean expression values for each STR dosage bin with at least 5% frequency in each group. Dosages were binned into groups spanning 3 repeat copies each since individually each genotype was relatively rare at this locus. **(c)** Positioning of the CCG repeat relative to the H3K27ac signal (note the localization within the nadir of the signal, which indicates a nucleosome depleted region) and a CTCF binding site at the 5' UTR of *RHOT1*. The visualization was generated using the Integrative Genomics Viewer[27] loading the ENCODE[28] data for GM12878 LCLs. The image does not display the gene NR_136413 that also overlaps the STR as it is not expressed in LCLs. **(d)** Association of variants at the *RHOT1* locus and red blood cell distribution width, before (top) and after (bottom) conditioning on the CCG repeat at chr17:30469471 (hg19). Light blue=SNPs and indels; orange=STRs. Red line=significance threshold, black circle=the $(CCG)_n$ STR.
